# Robotic Search for Optimal Cell Culture in Regenerative Medicine

**DOI:** 10.1101/2020.11.25.392936

**Authors:** Genki N. Kanda, Taku Tsuzuki, Motoki Terada, Noriko Sakai, Naohiro Motozawa, Tomohiro Masuda, Mitsuhiro Nishida, Chihaya T. Watanabe, Tatsuki Higashi, Shuhei A. Horiguchi, Taku Kudo, Motohisa Kamei, Genshiro A. Sunagawa, Kenji Matsukuma, Takeshi Sakurada, Yosuke Ozawa, Masayo Takahashi, Koichi Takahashi, Tohru Natsume

## Abstract

Induced differentiation is one of the most experience- and skill-dependent experimental processes in regenerative medicine, and establishing optimal conditions often takes years. We developed a robotic AI system with a batch Bayesian optimization algorithm that autonomously induces the differentiation of induced pluripotent stem cell-derived retinal pigment epithelial (iPSC-RPE) cells. The system performed 216 forty-day cell culture experiments, with a total experimentation time of 8,640 days. From 200 million possible parameter combinations, the system performed cell culture in 143 different conditions in 111 days, resulting in 88% better iPSC-RPE production than that by the pre-optimized culture in terms of pigmented scores. Our work demonstrates that the use of autonomous robotic AI systems drastically accelerates systematic and unbiased exploration of experimental search space, suggesting immense use in medicine and research.

## INTRODUCTION

Automating scientific discovery is one of the grandest challenges of the 21st century (*1*, *2*). A promising approach involves creating a closed loop of computation and experimentation by combining AI and robotics (*3*). A relatively simple form of autonomous knowledge discovery involves searching for optimal experimental procedures and parameter sets through repeated experimentation and result validation, according to a predefined validation method. For example, in material science, the parameters associated with the growth of carbon nanotubes have been explored using an autonomous closed-loop learning system (*4*). In experimental physics, Bayesian optimization has been used to identify the optimal evaporation ramp conditions for Bose–Einstein condensate production (*5*). In 2019, a promoter-combination search in molecular biology was automated using an optimization algorithm-driven robotic system (*6*).

Here, we report the development of a robotic search system that autonomously determines the optimal conditions for cell culture. Cell culture is probably one of the most delicate procedures in two respects. First, the parameters related to physical manipulation can greatly affect the outcome of the experiment (*7*). Secondly, it takes a long time to execute a series of protocols. For example, in regenerative medicine, cells need to be artificially differentiated from embryonic stem cells or induced pluripotent stem cells (ES/iPS cells) through hundreds of experimental procedures that typically last for weeks or months. During these processes, cells are given chemical inputs (e.g., type, dose, and timing of reagents) and physical inputs (e.g., strength of pipetting, vibration during handling of plates, timing of transfer from/to CO_2_ incubator, and accompanying changes in temperature, humidity, CO_2_ concentration etc.). Due to the heterogeneous and complex internal states of cells, suitable culture conditions must be determined for each strain and/or lot (*8*). A small difference in a single chemical stimulus or physical procedure can lead to failure of differentiation or poor quality of the produced cells, and such consequences can often become experimentally detectable only days or weeks after the input is given (*9*). Therefore, manual cell culture, which is highly skill-dependent, is often inefficient, error-prone, and lacks scalability, which is a crucial drawback in medical and industrial applications.

It is advantageous to utilize high-accuracy and programmable robot arms for the search of optimal cell culture parameters. Because programs and sensors describe everything that occurs in the lab, robotization realizes nearly perfect control and parameterization of experimental procedures. Furthermore, unlike human hands, robot arms can repeat the same procedure many times, ensuring reproducibility by keeping all the parameters related to physical procedures constant. Although some automated cell culture machines have already been proposed (*10*), proper formulation of an autonomous search for optimal culture conditions has not yet been determined.

In this study, we combined a Maholo LabDroid (*11*) and an AI system that independently evaluates the experimental results and plans the next experiments to realize an autonomous robotic search for optimal culture conditions. We first created a digital representation of the regenerative medical cell culture protocol used for the induction of retinal pigment epithelial (RPE) cells from iPS cells (iPSC-RPE cells) (*12*), which can be executed by the robot and used as a template for an AI-driven parameter search. We then implemented the experimental protocol on a LabDroid, which is a versatile humanoid robot that can perform a broad range of experimental procedures. Its flexibility allows frequent changes in protocols and protocol parameters, making it suitable for use in experimental parameter searches. The robot has an integrated microscope, providing data for image-processing through AI which evaluates the quality of growing cells. The search process was mathematically formulated as a type of experimental design problem, and a batch Bayesian optimization (BBO; **Figs. 1, S1, S2**) technique was employed as a solver. Finally, we demonstrated that iPSC-RPE cells generated by LabDroid satisfy the cell biological criteria for regenerative medicine research applications.

**Fig. 1:**
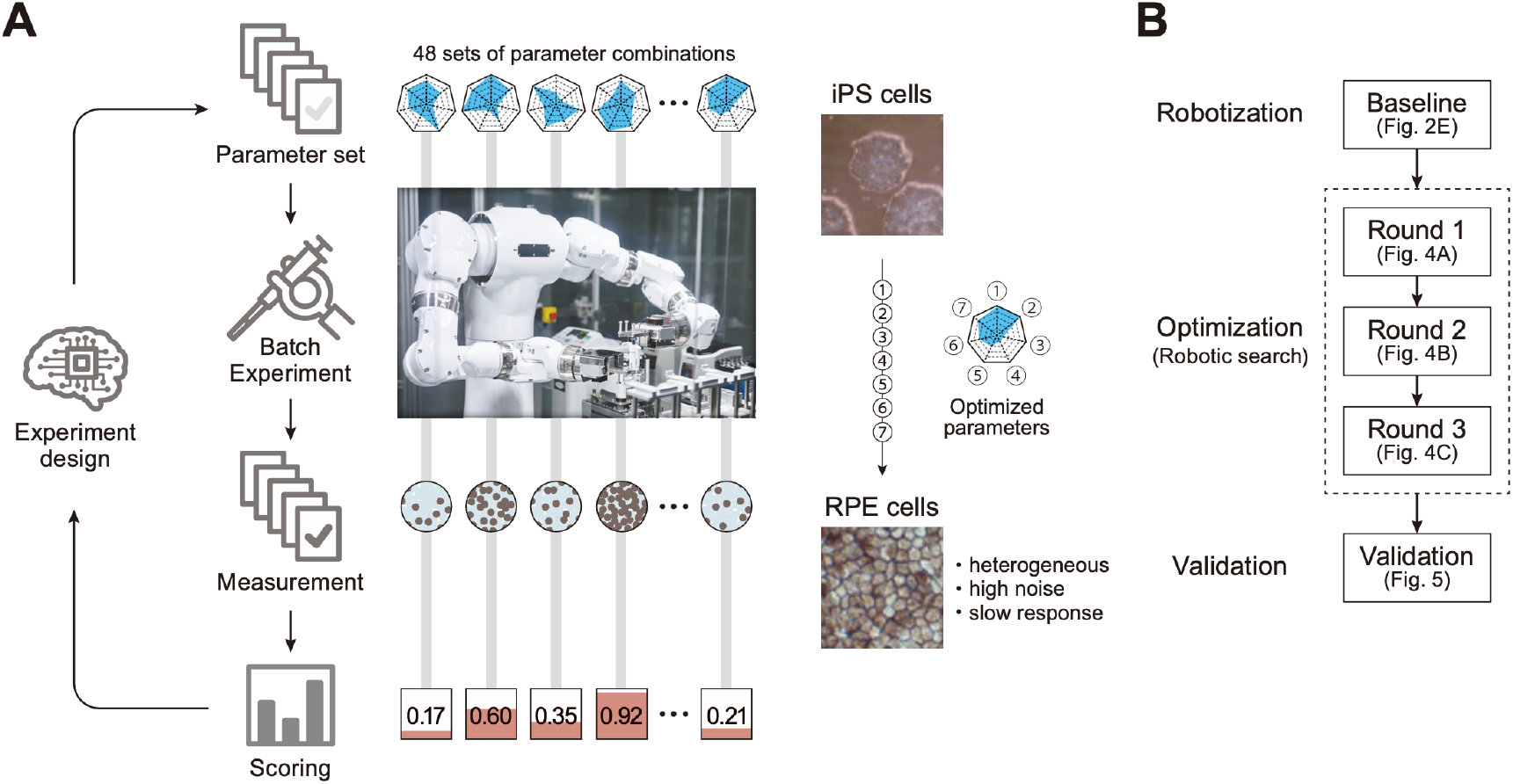
Robotic search for optimal experimental conditions. (**A**) Overall workflow for the optimization of experimental procedures using combined experimental robotics and Bayesian optimization. The user defines the target experimental protocol, subject parameters of the protocol, and the validation function. In this study, we chose the differentiation procedure from iPS to RPE cells as a target protocol and selected the reagent concentration, administration period, and five other parameters (details are shown in **Table 1**). We defined the pigmented area in a culture well, which represents the degree of RPE differentiation induction, as the validation function. The optimization program presented multiple parameter candidates; the LabDroid performed the experiment, and then an evaluation value for each candidate was obtained. Subsequently, the Bayesian optimization presented a plurality of parameter candidates predicted to produce higher validation values. The optimal parameters were searched by repeating candidate presentation, experiment execution, validation, and prediction. The detailed components are shown in **Fig. S2**. (**B**) Workflows performed in this study. First, robotization of the iPSC-RPE protocol was performed as a baseline. Next, the optimization process was conducted in three rounds, followed by statistical and biological validation. The figure numbers in parentheses represents the results shown in the figure.

## RESULTS

### Robotization of the iPSC-RPE differentiation protocol

An overview of the iPSC-RPE differentiation protocol used for optimization is shown in **Figs. 2A** and **S1**. It consists of five steps: seeding, preconditioning, passage, induction of RPE differentiation (induction), and RPE maintenance culture. The day on which the passage was performed was defined as differentiation day (DDay) 0, and the cultured cells were sampled and validated on DDays 33 and 34. To implement this protocol using LabDroid, the necessary peripheral devices were installed on and around LabDroid’s workbench (**Figs. 2B**, **S3**). We designed the system to work simultaneously with eight 6-well plates per batch, for a total of 48 cell-containing wells. LabDroid was programmed for three types of operations: seeding, medium exchange, and passage (**Figs. S4–9**; **Table S1**; **Movie S1**). The steps for the preconditioning and induction, which correspond to the preparation of reagents, were named medium exchange type I, and the step for RPE maintenance culture, which does not involve reagent preparation, was named medium exchange type II (**Figs. 2A**, **S4**).

**Fig. 2:**
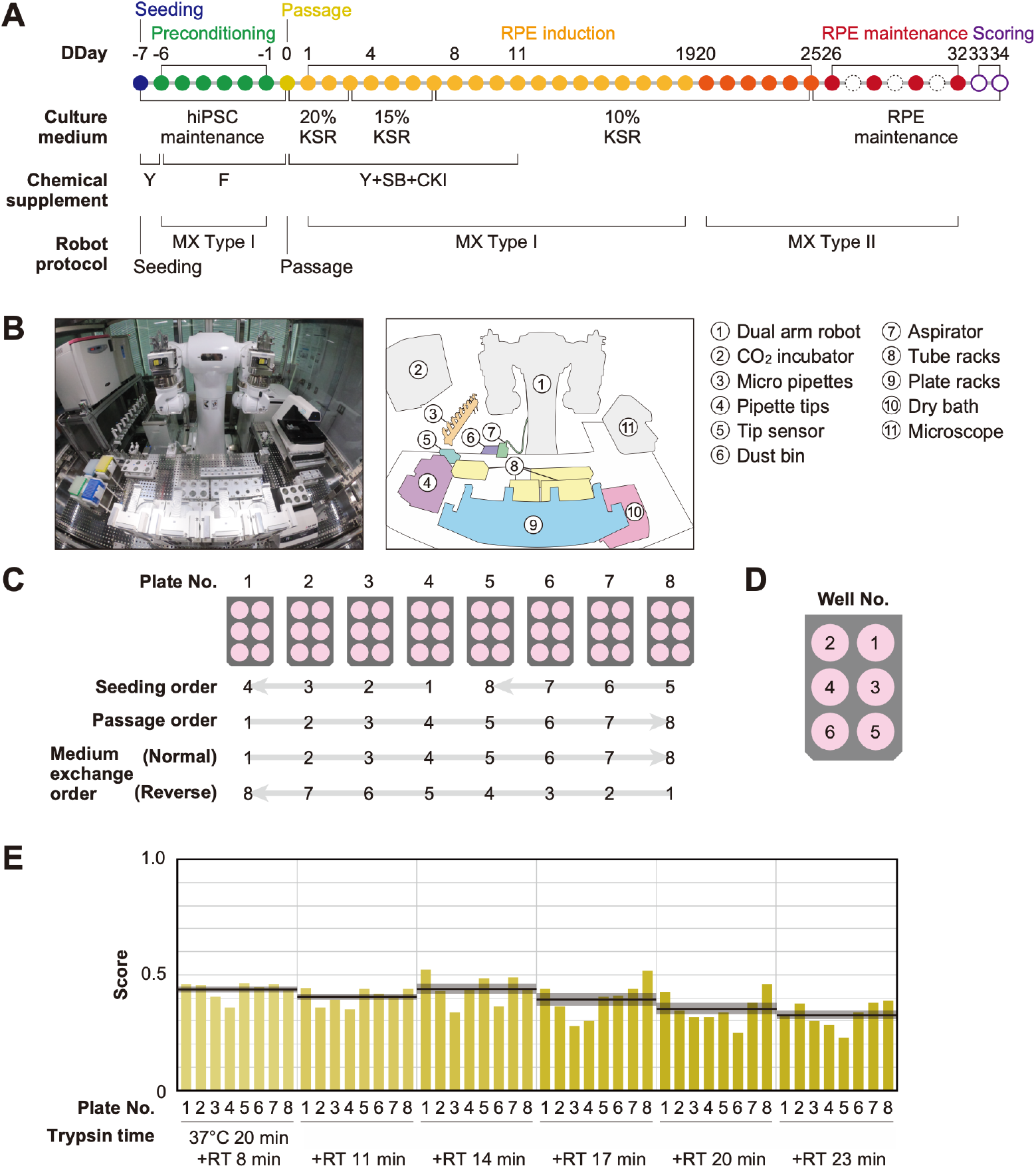
Robotization of iPSC-RPE differentiation protocols. (**A**) Schematic diagram of the standard iPSC-RPE differentiation procedures. DDay indicates the differentiation day. Filled circles represent days when the robot operated, solid circles represent days with human operations only, and dashed line circles represent days when no operations were conducted. F stands for FGF receptor inhibitor; Y for Y-27632, a Rho-kinase inhibitor; SB for SB431542, a TGF-β/Activin/Nodal signal inhibitor; CK for a CKI-7, Wnt signal inhibitor; and MX for medium exchange. (**B**) The LabDroid Maholo including peripheral equipment. (**C**) Plate numbering and the orders of seeding, passage, and medium exchange operations. Eight 6-well plates were used for each experiment. (**D**) Well numbering. (**E**) Scores of the first trial. iPSC-RPE differentiation was conducted under six different trypsin treatment times using the LabDroid. Vertical blue bars represent the pigmented cell area score of each well. The bold black lines and the shaded area around the lines represent the mean score and SEM of eight samples operated at the same trypsin time, respectively. The raw values are shown in **Table S2**.

First, we used LabDroid to perform baseline experiments involving iPSC-RPE induced differentiation under the same conditions as the typical manual operations. Because of the differences in structure and experimental environment between the LabDroid and humans, some operations and movements, such as the use of a centrifuge, the presence or absence of cell counting at the time of passage, and the speed of movement, differed from those of humans. For example, achieving the same time interval for trypsin treatment in all wells of a single plate during cell detachment using LabDroid is difficult. Therefore, the passage operation was performed at six separate time intervals. The cells differentiating into RPEs produce melanin, which causes them to turn brown. Therefore, the area ratio of the total number of pigmented cells on DDay 34 was used to estimate the differentiation induction efficiency and obtain evaluation scores, following the example of previous studies (*10*, *13*) (**Fig. S14**). These validation scores were used to simplify the validation process and do not reflect the entire quality of the RPE.

Baseline experiments were conducted and validated using six trypsin conditions and eight plates (**Figs. 2C–E**, **S15**; **Table S2**). The highest-scoring trypsin treatment was 20 min at 37 °C, followed by 14 min incubation at room temperature, with an eight-plate score of 0.44 ± 0.03 (mean ± SEM, n = 8). The lowest-scoring of the six trypsin conditions was incubated for 20 min at 37 °C followed by 23 min at room temperature, with an eight-plate score of 0.33 ± 0.02 (mean ± SEM, n = 8). LabDroid successfully performed the iPSC-RPE protocol, as evidenced by the detection of pigmented cells in all 48 wells and the lack of errors in the operating process. However, in the naive transplantation of the manual protocol to the robot, the induction efficiency was insufficient. This suggests that it is inherently difficult to describe physical parameters, including unrecorded human movements. Therefore, we attempted to optimize the protocol parameters to further improve the scores using a robotic search.

### Parameterization of the protocol

To improve the pigmentation score, we selected seven parameters for optimization: two from the preconditioning step, three from the passage step, and two from the induction step. Search domains were set for each parameter (**Table 1**; **Fig. 3A, B**).

**Fig. 3:**
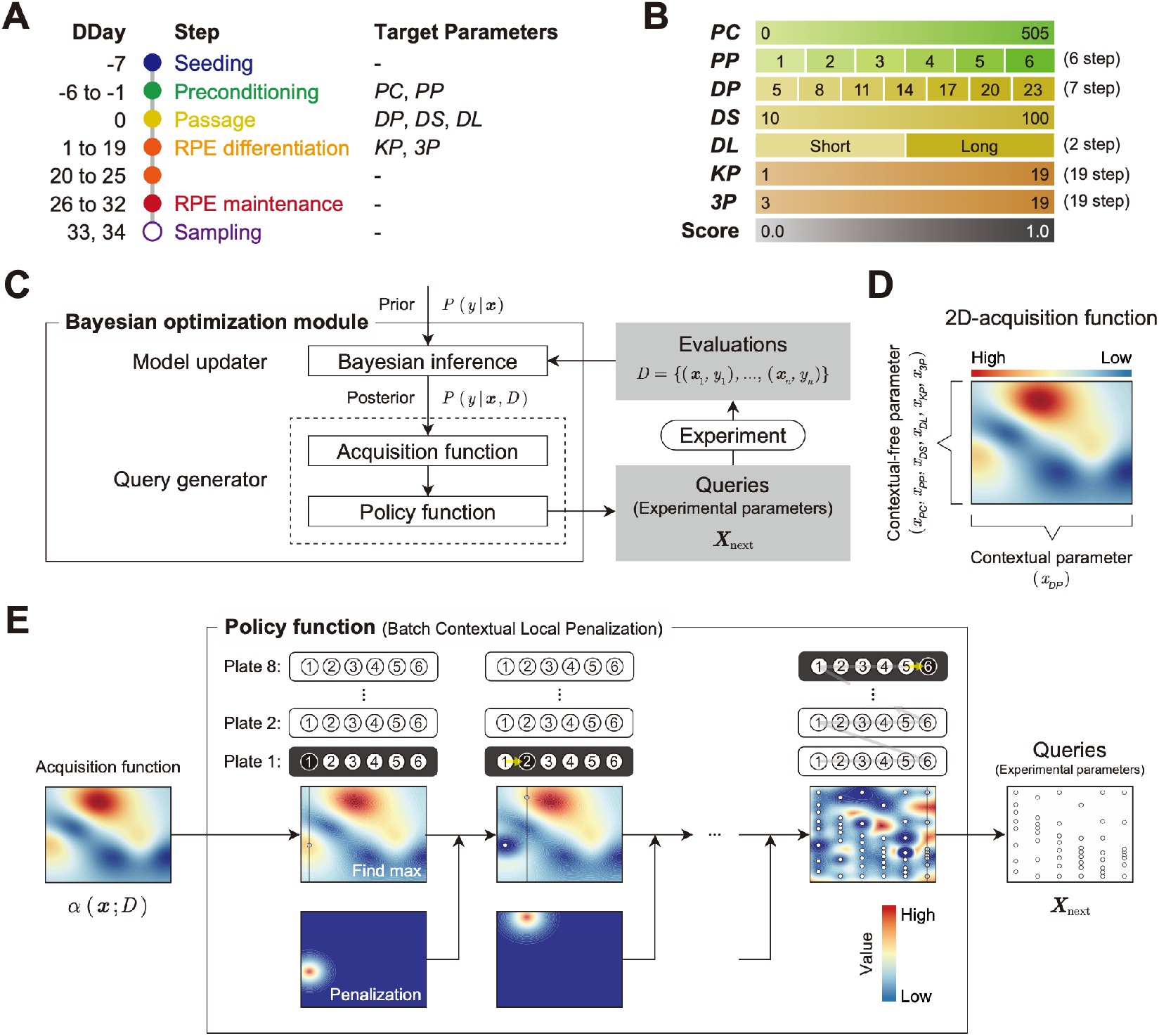
Optimization module. (**A**) Definition of the target parameters and corresponding steps in the protocol: *PC*, preconditioning concentration; *PP*, preconditioning period; *DP*, detachment trypsin period; *DS*, detachment pipetting strength; *DL*, detachment pipetting length; *KP*, KSR concentration reducing period; and *3P*, three chemical (Y, SB, CKI) supplement administration period. (**B**) Ranges and stepping of the parameters. (**C**) The Bayesian optimization module consists of two components: a Model updater and a Query generator. The Model updater updates the Gaussian process posterior on the experiment using all available data 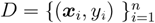, where x indicates experimental parameter, and y indicates corresponding evaluation score. The Query generator calculates the acquisition function *α*(*x*; *D*) for an experiment parameter *x* with the posterior distribution *P*(*y*\*x.D*), and generates the experiment parameter set *X*_next_. for the next 48 points using the policy function with *α*(*x*; *D*). (**D and E**) Test of the query generation process using a two-dimensional toy acquisition function. (**D**) Values of the toy acquisition function given an experimental parameter set. The horizontal axis represents the input values of *^x^DP* (contextual parameter), whereas the vertical axis represents the input values of the other six remaining context-free parameters *X* = (*x_PC_*, *x_PP_*, *x_DS_*, *x_DL_, *x*_3P_*, *x_KP_*) which are collapsed into a single axis. The color of the heatmap indicates the value of the acquisition function. In the heat map, the acquisition value is higher in places where the color is closer to red and lower in places where the color is closer to blue. (**E**) Test of the query generation process for the experimental parameter set *X*_next_ in the next experiment using a batch contextual local penalization policy (BCLP) policy. The heat maps in the upper row show the (penalized) acquisition function values, and the lower row shows the penalization values for the acquisition function. The queries *X*_next_ for 48 wells (right side figure) were iteratively generated from the maximization-penalization loop on the acquisition function.

**Table 1:** Definition of optimized parameters. Parameter names, parameter name codes, description, parameter ranges, parameter units, correspondence between experimental procedure and parameters used (related to **Figs. 2A, 3A, B**).

| Parameter name | Code | Description | Range | Unit | Prptocol step |
| --- | --- | --- | --- | --- | --- |
| Preconditioning concentration | PC | FGFRi concentration in medium | 10–505 | nM | Preconditioning |
| Preconditioning period | PP | FGFRi duration in medium | 1–6 | day | Preconditioning |
| Detachment trypsin period | DP | Trypsin incubation duration at room temperature after incubation at 37°C, 20 min. | 5, 8, 11, 14, 17, 20, 23 | min | Passage |
| Detachment pipetting strength | DS | Pipetting strength during cell detachment | 10–100 | mm/sec | Passage |
| Detachment pipetting length | DL | Bottom surface area to be pipetted | short / long | N/A | Passage |
| KSR period | KP | KSR concentration and duration in medium:<br>KSR concentration is decreased linearly every day so that KSR becomes 10% on DDday of KP value | 1–19 | day | RPE differentiation |
| Three supplements period | 3P | Three chemical supplements duration | 3–19 | day | RPE differentiation |

From the preconditioning step on DDays −1 to −6, we selected two parameters for optimization: the concentration of fibroblast growth factor receptors inhibitor (FGFRi) in the medium (*PC*, preconditioning concentration), and the duration of addition (*PP*, preconditioning period). From the passage step performed on DDay 0, we selected three parameters to optimize: the pipetting strength during cell detachment (*DS*, detachment pipetting strength), the area of the bottom surface to be pipetted (*DL*, detachment pipetting length), and trypsin processing time (*DP*, detachment trypsin period) of a passage. *DP* is a contextual parameter that can only be used to perform experiments at fixed values, owing to the specifications of the experimental system. In this case, *DP* is allowed to take different fixed values at three-minute intervals, corresponding to the number of wells in the plate. From the RPE differentiation induction step on DDays 1 to 25, we selected two parameters to optimize: the concentration of KnockOut Serum Replacement (KSR) in the medium (*KP*, KSR period), and the duration of exposure period of the three chemical supplements (*3P*, three supplement period).

### Optimization of the protocol

To improve the optimization performance, 48 conditions (eight plates × six wells, as shown in **Fig. 2C**) were executed in parallel in each batch. In general, solving a high-dimensional, expensive black-box optimization problem such as the present one, with a limited number of rounds, is challenging. In our case, some 200 million possible parameter combinations existed in the search space, and the point where the pigmented score was optimal in three rounds (144 queries) had to be determined, because one experiment round took 40–45 days. In recent studies, batch Bayesian optimization (BBO) has shown excellent performance in real-world black-box optimization problems (*6*, *14*, *15*). We integrated an experimental design module based on BBO to effectively search for the optimal experimental parameters that maximize the pigmentation scores in the search space defined in (**Fig. 3B**).

The Bayesian optimization module generates queries using two components: the Model updater, which updates the surrogate model that captures the relationship between parameters and the scores using Bayesian inference (**Fig. S10**), and the Query generator that generates the next experimental parameters *X*_next_ using an acquisition function and a policy function (**Figs. 3C**, **S11**; **Algorithms S1–S3**). In the Query generator, the acquisition function estimates the expected progress toward the optimal experimental parameter at a given experimental parameter (**Fig. 3D**). Then, using the acquisition function, the policy function generates the next 48 experimental parameters *X*_next_ considering the context of trypsin processing time ^*x*^DP (**Fig. 3E**).

To test the performance of the Bayesian optimization module in our case, we executed a preliminary performance validation using a toy testing function constructed on domain knowledge (**Figs. S12, S13**).

### Robotic optimization drastically improved pigmented score

In this study, three successive experiments were conducted to optimize the target protocol. In each round, 48 conditions were generated using the Bayesian optimization module and translated into LabDroid operating programs. In accordance with the experimental design, we incorporated the two highest-scoring conditions from the previous experiment (**Fig. 2E**) as control conditions, performed differentiation-inducing cultures with the LabDroid, and validated the area of the colored cells. In round 1, although one condition was found to be experimentally deficient, the other 47 conditions were validated. The highest score was 0.86 (**Figs. 4A**, **S16**; **Table S3**), yielding five conditions that exceeded the mean value (0.39) for all wells in the baseline experiment (**Fig. 2E**). In round 2, 46 conditions were generated, and the two highest-scoring conditions in round 1 were incorporated as control conditions. The highest score was 0.83 (**Figs. 4B**, **S17**; **Table S3**). In round 3, 48 experiments were conducted, yielding an improved highest score of 0.91. We obtained 26 other conditions that were better than the highest in round 2 (**Figs. 4C**, **S18**; **Table S3**). A visualization diagram of a two-dimensional partial least squares regression (PLS) clearly revealed that the overall experimental parameters tended to converge in a higher pigmented score direction from rounds 1 to 3 (**Figs. 4D**, **S20**).

**Fig. 4:**
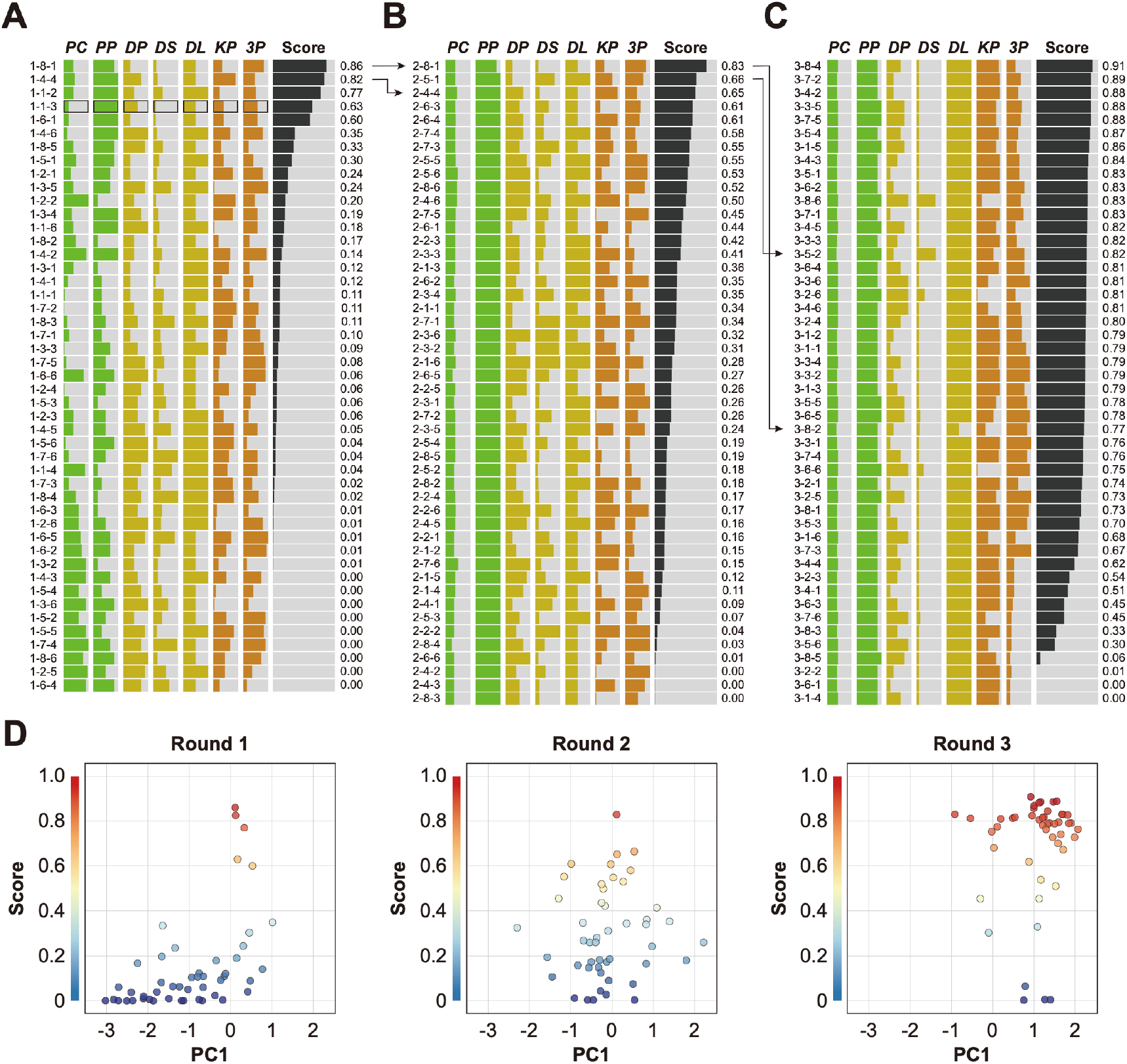
Robotic search for optimal parameters in iPSC-RPE differentiation. (**A–C**) Parameter candidates sorted in order of the pigmentation score in optimization rounds 1 (**A**), 2 (**B**), and 3 (**C**). The ID label on the left represents ‘Round No. - Plate No. - Well No.’. For example, “1-2-3” means “(Round) 1-(Plate) 2-(Well) 3.” The parameter values and resulting pigmentation scores are plotted as horizontal bars. The parameter candidate with black frames (1-1-3) in (**A**) is the standard condition. Arrows indicate the control experiments; the top two conditions in round 1 were included in round 2, and the top two conditions in round 2 were implemented in round 3. The raw values are shown in **Table S3**. (**D**) Visualization of the parameter set and pigmentation score distributions using partial least squares regression (PLS) in each round. The horizontal axis PC1 shows the values of the parameter candidates that are projected onto the first component of the PLS. The vertical axis shows the pigmentation score for each candidate parameter. As the rounds progressed, the overall score tended to converge in a higher direction. A full visualization of the experimental results using a parallel coordinate plot (PCP) is shown in **Fig. S20**.

To determine whether the optimized conditions were statistically improved over the pre-optimized conditions, a multi-well validation experiment was conducted using the top five conditions in round 3 and the pre-optimized conditions. The validation values, ordered by place, were 0.71 ± 0.06, 0.72 ± 0.03, 0.76 ± 0.02, 0.79 ± 0.02, and 0.81 ± 0.02 (mean ± SEM, n = 3 each). All scores were statistically significantly higher than the pre-optimization scores of 0.43 ± 0.02 (mean ± SEM, n = 3) (**Figs. 5A, B**, **S19**; **Table S4**).

**Fig. 5:**
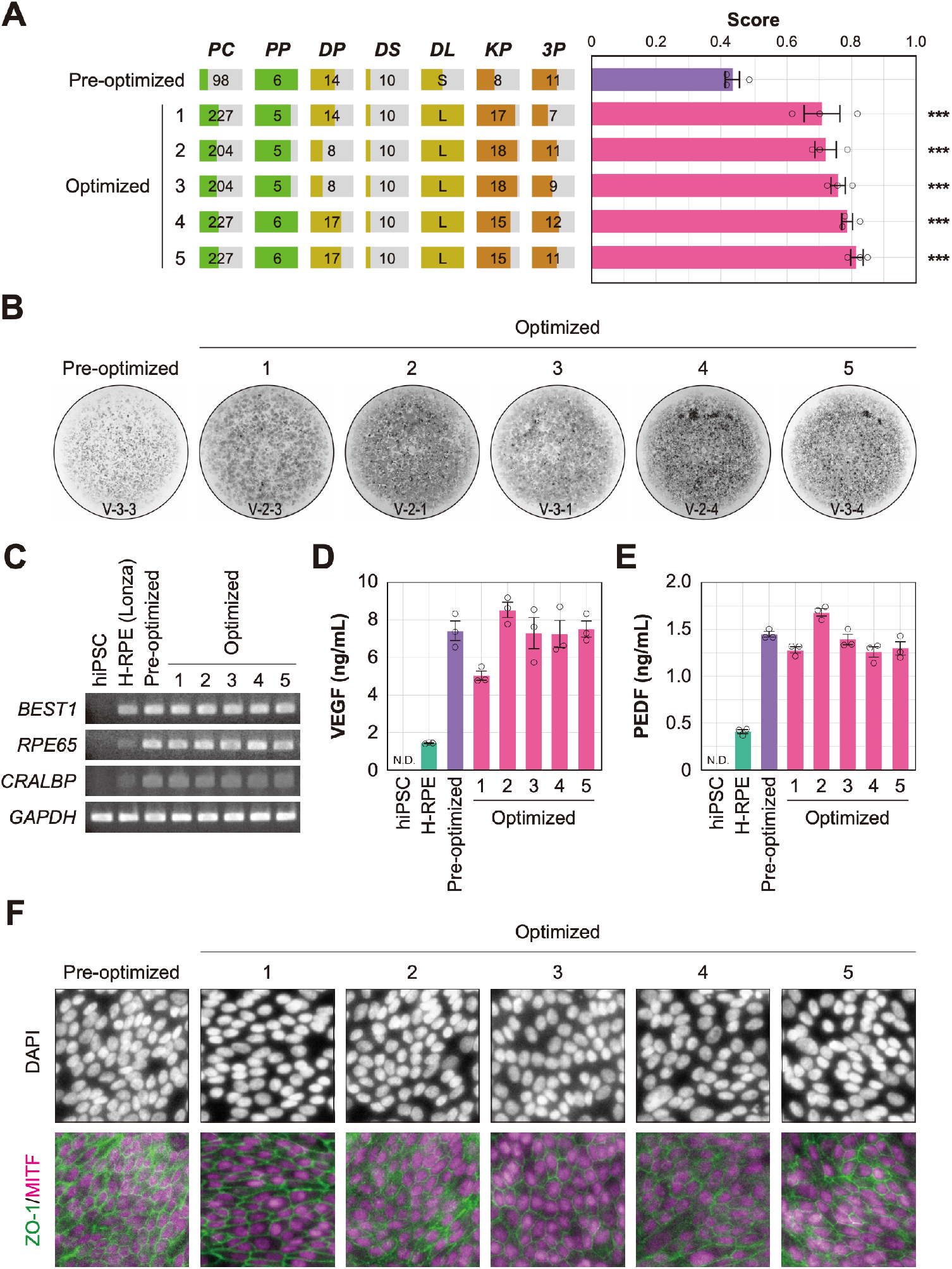
Quality evaluation of robot-induced RPE cells. (**A**) Pigmentation score evaluation of the pre-optimized (n = 3) and top five conditions (n = 3 each) from round 3. Error bars represent the standard error of the mean (SEM). The numbers 1–5 in the optimized group represent the first to fifth place conditions for round 3 (**Fig. 4C**). Circles represent an individual score, bars represent the mean score, and error bars represent the SEM. Statistical significance was examined using two-way ANOVA and SNK *post-hoc* tests. *P* < 0.05 was considered significant. \*\*\**P* < 0.001 versus pre-optimized. In all other combinations no statistical significance was detected. Raw values are shown in **Table S4**. (**B**) Representative pigmented images of the pre-optimized and five optimized iPSC-RPE cells. Images acquired on DDay 34. ID labeling on the bottom reads ‘V (validation) - Plate No. - Well No.’. The other images are shown in **Fig. S19**. (**C–F**) Cell biological validation of the robot-induced RPE cells. After DDay 34, cells were purified, stocked, initiated, maintained for four weeks, and analyzed (**Fig. S1B**). (**C**) Representative marker gene expression in RPE cells by RT-PCR. iPSC, undifferentiated iPSC; H-RPE (Lonza), Clonetics H-RPE (Lot #493461, Lonza, USA); pre-optimized and optimized LabDroid-induced RPE. (**D–E**) Quantification of representative secreted proteins from iPSC-RPE cells using ELISA. The supernatants were collected and the amount of VEGF (**D**) and PEDF (**E**) in the culture medium was analyzed 24 h after medium exchange (n = 3 wells each). Circles represent individual scores, bars represent the mean score, and error bars represent SEM. n.d. = not detected. The raw values are shown in **Table S6**. (**F**) Co-staining of ZO-1 (green) and MITF (magenta) using immunohistochemistry. Nuclei were stained with DAPI.

In summary, we conducted 216 forty-day cell culture experiments with a total experimentation time of 8640 days. We accelerated the search using a BBO technique, compressing the search time to 185 days with a cumulative robot operating time of 995 h (**Table S5**; **Figs. S21**, **S22**; **Movies 2–6**).

In this study, we succeeded in replacing part of the process of moving from iPS cells to the production of RPE cells for transplantation using robots and demonstrated an effective optimization method (**Fig. S2**). However, it was not obvious that robot-manufactured RPE cells would produce the cell characteristics required for transplanted cells obtained using manual preparation. Therefore, we purified the differentiation-inducing cells of the validation round, prepared the cells just before transplantation, and performed a biological quality evaluation (**Fig. S1B**). The analyzed iPSC-RPE cells expressed *BEST1*, *RPE65*, and *CRALBP* (**Fig. 5C**), which are characteristic marker genes of RPE cells. Secretion of VEGF and PEDF into the culture medium, a characteristic of RPE cells, was observed (**Fig. 5D, E; Table S6**). The expression of tight junction-associated factor ZO-1 was examined using immunohistochemistry, and a ZO-1-derived fluorescence signal was observed in microphthalmia-associated transcription factor (MITF)-positive cells, which play a central role in RPE cell function (**Fig. 5F**). These results indicated that the robot-manufactured iPSC-RPE cells had the characteristics of RPE cells, and fulfilled the criteria for use in regenerative medicine research using the type of analysis measured in a previous clinical study (*12*).

## DISCUSSION

Laboratory automation is a recently developing technology that transfers human skills to machines. Although some robotic systems for cell culture have already been developed (*16*–*24*), many of these fixed-process automation apparatuses lack the flexibility and precision necessary to execute comprehensive parameter searching. Biological cells are physical systems with rich internal dynamics (*25*) and have lower tolerance for differences in manufacturing processes than products derived from metal fabrication, chemical synthesis, or other similar approaches (*9*), underscoring the need for closed-loop optimization.

By combining the LabDroid and BBO algorithm, our robotic search system autonomously discovered optimal conditions that improved the efficiency of differentiation induction in iPSC-RPE production by up to 88% (**Fig. 5A**). We chose the iPSC-RPE differentiation protocol for three reasons. First, melanin pigmentation is a single, easily measurable morphological indicator of the quality and quantity of successfully differentiated RPE cells. The pigmentation score is a well-established measure of RPE differentiation quality that can be easily verified with the naked eye. Integrating other modalities of data, such as the RNA expression and secretory protein data shown in **Fig. 5C–F** to the scoring function will greatly improve the accuracy of iPSC-RPE cell quality estimation. It should be noted that, in the analysis described in **Fig. 5C–F**, there was no change in cell quality indicators before and after optimization, because these cells were analyzed after the purification process. In clinical transplantation of iPSC-RPE, appropriate cells among induced cells are picked up and cultured to prevent contamination of cells that did not induce properly (purification process; **Fig. S1A**). For a patient, even if the efficiency of induction is extremely low, the appropriate cells can be selected for transplantation. By contrast, for transplantation to a large number of patients, it is necessary to increase the final cell number by increasing the efficiency of induction as much as possible. Second, the operating accuracy and repeatability of the robotic system used in this study were satisfactory for the efficient completion of the search process. The iPSC-RPE cell differentiation protocol requires 40 days to run, and a single mis-operation, error, or inaccuracy can deteriorate the search efficiency if the entire process is not destroyed. Third, iPSC-RPE cells have already been clinically transplanted into human patients, and a well-established protocol is available (*12*). Data and expertise accumulated in manual differentiation induction experiments provide useful information for establishing the general structure of the protocol and in defining its parameter search space. However, many other cell types and differentiation targets lack established protocols, some of which probably demand a more sophisticated optimization technique that can deal with categorical values and their combinations. Such a technique could optimize the structure of the protocol simultaneously with continuous parameter values while minimizing the execution costs incurred by a large number of possible combinations (*26*).

## ACKNOWLEDGEMENTS

We thank T. Mitsuyama, T. Iwata, and N. Yachie for their insightful comments; E. Takagi and H. Hirabayashi for research support; H. Uchida and A. Kato for illustrations; and J. Freeman for carefully proofreading the manuscript. We also thank all the laboratory members at RIKEN BDR, in particular, N. Koide, Y. Shibata, A. Maeda, and T. Maeda for their kind help in the preparation of the materials, their support in the experiments, and their insightful discussions. This work was supported by a grant from AMED (JP20bm0204002, AMED, to M. Takahashi), a project subsidized by the New Energy and Industrial Technology Development Organization (NEDO, to M. Takahashi), JST-Mirai Program (JPMJMI18G4, JST, to K. T.), an internal grant from the RIKEN Center for Biosystems Dynamics Research (RIKEN, to M. Takahashi and K. T.), RIKEN Engineering Network (RIKEN, to M. Takahashi), RIKEN Junior Research Associate program for graduate students (RIKEN, to N. M.).

## AUTHOR CONTRIBUTIONS

K.T., and T.N. conceived the idea. G.N.K., T.T., T.S., Y.O., M. Takahashi, and T.N. designed the project. G.N.K., M. Terada, N.S., N.M., T.M., M.N., and G.A.S. performed the biological experiments. T.T., C.T.W., T.H., S.H., T.S., and Y.O. performed software development and informatics experiments. T.K., M.K., K.M., and T.N. performed robot teaching and maintenance. T.S. performed project management. G.N.K performed the project administration. M. Takahashi and T.N. supervised the project. G.N.K., T.T., and Y.O. wrote the manuscript. All authors discussed the results and commented on the manuscript.

## DECLARATION OF INTERESTS

G.N.K., T.K., M.K., K.M., and T.N. are employees, executives, or stakeholders of the Robotic Biology Institute, Inc., which may benefit financially from the increased scientific use of LabDroid Maholo. T.T., C.T.W., T.H., S.H., T.S., Y.O., and K.T. are employees, shareholders, or stakeholders of Epistra Inc., which may benefit financially from the increased scientific use of the developed software. All the other authors declare no competing interests.

## Notes

### Competing Interest Statement

G.N.K., T.K., M.K., K.M., and T.N. are employees, executives, or stakeholders of Robotic Biology Institute Inc., which may benefit financially from the increased scientific use of LabDroid Maholo. T.T., C.T.W., T.H., S.H., T.S., Y.O., and K.T. are employees, shareholders, or stakeholders of Epistra Inc., which may benefit financially from the increased scientific use of developed software. All other authors declare no competing interests.

### Summary of Updates

Supplemental files updated.

