## Supplementary Materials for "Robotic Search for Optimal Cell Culture in Regenerative Medicine"

### Table of Contents

1. Materials and Methods
2. System Outline
    2.1. A target process of our robotic search
    2.2. System components
3. Robotic Components
    3.1. Robotic setup
    3.2. Robotic workflows
    3.3. Micropipette combination for the transfer of a wide range of volumes with LabDroid
4. Informational Components
    4.1. Compiling experimental design table to robot protocols
    4.2. Parameterization of iPSC-RPE protocol
    4.3. Gaussian process overview
    4.4. Bayesian optimization overview
    4.5. Bayesian optimization for the iPSC-RPE differentiation protocol
    4.6. Testing optimization in simulation
5. Supplementary Results
    5.1. Image analysis for scoring
    5.2. Score data visualization by PCP
6. Experimental Operations
    6.1. Daily robot experimental operation
    6.2. 24/7 monitoring and recording
    6.3. Execution logs and errors
7. Supplementary Table Legends
8. Supplementary Movie Legends
9. Supplementary References

### 1. Materials and Methods

*Guidelines*

All experiments that involved the use of human-derived samples were reviewed and approved by the institutional review board of the Institutional Committee of RIKEN Kobe Branch.

*Reagents*

hiPSC maintenance medium: 80% StemFit Basal Solution A and 20% StemFit iPS Expansion Solution B (#AK02N, Ajinomoto Co., Inc., Japan)

RPE differentiation medium (20% KSR): 0.10 mM MEM Non-Essential Amino Acids Solution (NEAA) (#11140050, Thermo Fisher Scientific Inc., MA, USA), 1.0 mM Sodium pyruvate (#S8636, Merck & Co., Inc., NJ, USA), 19% KnockOut Serum Replacement (KSR) (#10828028, Thermo Fisher Scientific Inc., MA, USA), 0.0007% 2-mercaptoethanol (#139-06861, FUJIFILM Wako Pure Chemical Corporation, Japan), 78 U/mL Benzylpenicillin Sodium, and 78 µg/mL Streptomycin sulphate (#15140122, Thermo Fisher Scientific Inc., MA, USA). All diluted in GMEM (#11710035, Thermo Fisher Scientific Inc., MA, USA)

RPE differentiation medium (15% KSR): 0.10 mM MEM Non-Essential Amino Acids Solution (NEAA) (#11140050, Thermo Fisher Scientific Inc., MA, USA), 0.99 mM sodium pyruvate (#S8636, Merck & Co., Inc., NJ, USA), 15% KnockOut Serum Replacement (KSR) (#10828028, Thermo Fisher Scientific Inc., MA, USA), 0.0007% 2-mercaptoethanol (#139-06861, FUJIFILM Wako Pure Chemical Corporation, Japan), 82 U/mL benzylpenicillin sodium, and 82 µg/mL streptomycin sulphate (#15140122, Thermo Fisher Scientific Inc., MA, USA). All diluted in GMEM (#11710035, Thermo Fisher Scientific Inc., MA, USA)

RPE differentiation medium (10% KSR): 0.094 mM MEM Non-Essential Amino Acids Solution (NEAA) (#11140050, Thermo Fisher Scientific Inc., MA, USA), 0.94 mM sodium pyruvate (#S8636, Merck & Co., Inc., NJ, USA), 10% KnockOut Serum Replacement (KSR) (#10828028, Thermo Fisher Scientific Inc., MA, USA), 0.0007% 2-mercaptoethanol (#139-06861, FUJIFILM Wako Pure Chemical Corporation, Japan), 85 U/mL benzylpenicillin sodium, and 85 µg/mL streptomycin sulphate (#15140122, Thermo Fisher Scientific Inc., MA, USA). All diluted in GMEM (#11710035, Thermo Fisher Scientific Inc., MA, USA)

RPE maintenance medium: 29% Nutrient Mixture F-12 (#N6658, Merck & Co., Inc., NJ, USA), 1.9 mM L-Glutamine (#G7513, Merck & Co., Inc., NJ, USA), 1.9% B-27 supplement, serum free (#17504044, Thermo Fisher Scientific Inc., MA, USA), 96 U/mL benzylpenicillin sodium, and 96 µg/mL streptomycin sulphate (#15140122, Thermo Fisher Scientific Inc., MA, USA). All diluted in DMEM (Low glucose) (#D6046, Merck & Co., Inc., NJ, USA)

FGF receptor inhibitor (FGFRi) stock: PD 173074 (#P2499-5MG, Merck & Co., Inc., NJ, USA) diluted in DMSO (#D2650-5X5ML, Merck & Co., Inc., NJ, USA)

Rho-kinase inhibitor (Y) stock (8–10 mM): CultureSure Y-27632 (#036-24023, FUJIFILM Wako Pure Chemical Corporation, Japan) diluted in distilled water (Otsuka Pharmaceutical Factory, Japan) to a final 10 µM concentration when added to the cell culture medium

TGF-β/Activin/Nodal signal inhibitor (SB) stock (4–5 mM): SB 431542 hydrate (#S4317-5MG, Merck & Co., Inc., NJ, USA) diluted in DMSO (#D2650-5X5ML, Merck & Co., Inc., NJ, USA) to a final 5 µM concentration when added to the cell culture medium

Wnt signal inhibitor (CK) stock (2.4–3 mM): CKI-7 dihydrochloride (#C0742-5MG, Merck & Co., Inc., NJ, USA) diluted in distilled water (Otsuka Pharmaceutical Factory, Japan) to a final 3 µM concentration when added to the cell culture medium

RPE adhesion medium: DMEM/F12 (D8437, Merck & Co., Inc., NJ, USA), 10% FBS (12007C, Nichirei Corporation, Japan)

RPE washing solution: 98% DMEM/F12 (D8437, Merck & Co., Inc., NJ, USA), 1 mM Sodium pyruvate (S8636, Merck & Co., Inc., NJ, USA), 2 mM L-Glutamine (G7513, Merck & Co., Inc., NJ, USA)

*Labware*

For human use: micropipette tip, 2140-05-HR/2149P-05/61849, Thermo Fisher Scientific Inc. (MA, USA); micropipette tip, 30389165, Mettler Toledo (OH, USA); micropipette tip, 737251, Greiner Bio-One International GmbH (Germany); disposable pipette, 356507, Corning Incorporated (NY, USA); disposable pipette, 606160/607160/760160/768160, Greiner Bio-One International GmbH (Germany); filtration, SLGVJ13SL, Merck & Co., Inc. (NJ, USA); filtration, SS-10LZ, Terumo Corporation (Japan); filtration, 431096/430281/431097/430282, Corning Incorporated (NY, USA); 1.5-mL tube, 72.692MS, Sarstedt K.K. (Japan); 15-mL tube, 352096, Corning Incorporated (NY, USA); 50-mL tube, 352070, Corning Incorporated (NY, USA).

For LabDroid use: 6-well plate, 353046, Corning Incorporated (NY, USA); 50 mL tube, MS-58500, Sumitomo Bakelite Co., Ltd. (Japan); micropipette tip, 3511-05-HR/3512-05-HR/94410313/94410713/94052550, Thermo Fisher Scientific Inc. (MA, USA).

*LabDroid Maholo booth*

LabDroid including peripheral equipment were placed inside a booth made of acrylic walls and a stainless steel frame with three fan-filter-units. The LabDroid booth included a dual-arm humanoid (Robotic Biology Institute Inc., Japan), a $CO_2$ incubator (APC-30D, ASTEC Co., Ltd., Japan), micropipettes (4641110N/4641030N/4641230N/4641210N, Thermo Fisher Scientific Inc., MA, USA), a tube rack (Robotic Biology Institute Inc., Japan), a plate rack (Robotic Biology Institute Inc., Japan), a dry bath (EC-40RA, AS ONE Corporation, Japan), a tip sensor (Robotic Biology Institute Inc., Japan), an aspirator (SP-30, Air Liquide, Italy), a dust bin (EPD3S, Sekisui Techno Moulding Co., Ltd., Japan), and a microscope (EVOS FL Auto 2, Thermo Fisher Scientific Inc., MA, USA).

*hiPSC culture — initiation and preparation of cell suspensions (human part)*

The hiPSC line 253G1 (*27*), made from human dermal fibroblast, was obtained from RIKEN BRC (HPS0002). The hiPSCs were cultured and differentiated with the method described previously (*28–30*).

On DDay-14, frozen hiPSCs were initiated by the following procedures: first, laminin-coated 6-well plates were prepared. Final concentration of 0.5 μg/cm$^2$ of iMatrix-511 (Matrixome Inc, Japan) diluted by PBS (-) was then added to each well of the four 6-well plates and incubated for a minimum of 60 min at 37˚C and 5% $CO_2$, after which 0.75 mL/well of hiPSC maintenance medium was added. The supernatant was then removed. Next, 1 mL/well of hiPSC maintenance medium containing Rho-kinase inhibitor (final 10 μM concentration) was added and the coated plates were incubated at 37°C and 5% $CO_2$ until further use.

For hiPSCs initiation, frozen vials of hiPSCs stored in liquid nitrogen were thawed in a water bath set at 37°C and suspended in 5 mL of hiPSC maintenance medium. After centrifugation (160 × *g*, 22°C, 4 min), the supernatant was removed and an appropriate volume of hiPSC maintenance medium with a final 10 μM Rho-kinase inhibitor concentration was added. After counting the cells with a hemocytometer, the cells were seeded into laminin-coated 6-well plates at 43,300–45,000 cells/1.5 mL medium/well.

On DDay -13, the medium was replaced with hiPSC maintenance medium without Rho-kinase inhibitor. On DDays -12 to -8, the medium was replaced with the same medium composition at 24–72 h intervals. On DDay -7, cells were collected from the plate, and cell suspensions were delivered to the LabDroid booth. The medium was aspirated, 2 mL/well of PBS (-) was gently added and then aspirated for washing, and then 1 mL of 0.5 x TrypLE Select CTS (#A12859-01, Thermo Fisher Scientific Inc., MA, USA) diluted in 0.5 mM EDTA/PBS (-) was added, followed by incubation at 37°C and 5% $CO_2$ for 10 to 20 min. Then, cells were detached by pipetting and collected into a 50-mL tube, to which 1 mL of hiPSC maintenance medium and 3 mL of PBS (-) were added. After centrifugation (160 × *g*, 22°C, 4 min), the supernatant was removed, 0.75 mL of hiPSC maintenance medium with 10 μM Rho-kinase inhibitor was added, and the cells were resuspended. The cell suspension was filtered through a 40-μm cell strainer (#352340, Corning Incorporated, USA) with an additional 0.75 mL of hiPSC maintenance medium. After counting the cells with a hemocytometer, the cell suspension was set to 133,400 cells/20 mL with hiPSC maintenance medium containing 10 μM Rho-kinase inhibitor in eight 50-mL tubes. To prepare the cell suspensions, eight 6-well plates coated with laminin were prepared. A final concentration of 0.5 μg/cm2 of iMatrix-511 (Matrixome Inc., Japan) diluted in PBS (-) was added to each well of four 6-well plates and incubated for a minimum of 60 min at 37°C and 5% $CO_2$.

*iPSC-RPE differentiation (LabDroid part)*

On DDay -7, the hiPSC suspension was seeded into eight 6-well plates by coating eight 6-well plates with laminin, and placing eight tubes of the iPSC suspension and labware in the appropriate positions. The task of seeding was initiated, and the robotic operation was performed by LabDroid (**Figs. S4A**, **S5**). After the robotic operation, the eight cell-seeded plates were exported and incubated at a $CO_2$ incubator outside the LabDroid booth.

On DDay -6, the eight seeded plates were imported into the $CO_2$ incubator of the LabDroid booth. The users prepared eight 50-mL tubes of hiPSC maintenance medium with a final 10 μM Rho-kinase inhibitor concentration and two 50-mL tubes of hiPSC maintenance medium with final 5 μM FGFRi and 10 μM Rho-kinase inhibitor concentrations. The reagents and labware were placed in the appropriate positions. The task of preconditioning was then initiated, and the robotic operation was performed by LabDroid (medium exchange type I; **Figs. S4B**, **S6**) .

On DDays -5 to -1, the users prepared eight 50-mL tubes of hiPSC maintenance medium without Rho-kinase inhibitor and two 50-mL tubes of hiPSC maintenance medium with a final 5 μM FGFRi concentration. The reagents and labware were placed in the appropriate positions. The task of preconditioning was initiated, and the robotic operation was performed by LabDroid (medium exchange type I; **Figs. S4B**, **S6**).

On DDay 0, the following procedure was used for the operation of four plates: the users prepared four 6-well plates coated with laminin. A final 0.5 μg/cm2 concentration of iMatrix-511 (Matrixome Inc., Japan) diluted in PBS (-) was added to each well of the four 6-well plates and then the plates were incubated for a minimum of 60 min at 37°C and 5% $CO_2$. The users also prepared two 50-mL tubes of PBS (-), two 50-mL tubes of 0.5 x TrypLE Select CTS (#A12859-01, Thermo Fisher Scientific Inc., MA, USA) diluted in 0.5 mM EDTA/PBS (-), and four plates with RPE differentiation medium (20% KSR) with final 10 μM Rho-kinase inhibitor/3 μM Wnt signal inhibitor/5 μM TGF-β/Activin/Nodal signal inhibitor (4 mL/well each). The cell plates, laminin-coated plates, plates with medium, reagents, and labware were placed in the appropriate positions. The task of passage was initiated, and robotic operations were performed by LabDroid (**Figs. S4D**, **S7**). After performing this operation twice (four plates each), the eight cell-passaged plates were exported and incubated in a $CO_2$ incubator outside the LabDroid booth.

On DDay 1, the eight cell-passaged plates were imported into the $CO_2$ incubator of the LabDroid booth. Users prepared eight 50-mL tubes of RPE differentiation medium (10% KSR), two 50-mL tubes of 100% KSR, one 50-mL tube of 4 mM Rho-kinase inhibitor stock/1.2 mM Wnt signal inhibitor stock, and one 50-mL tube of 4 mM TGF-β/Activin/Nodal signal inhibitor stock. The reagents and labware were placed in the appropriate positions. The task of RPE differentiation was initiated, and the robotic operation was performed by LabDroid (medium exchange type I; **Figs. S4B**, **S8**).

On DDays 2 to 19, the users prepared eight 50-mL tubes of RPE differentiation medium (10% KSR), two 50-mL tubes of 100% KSR, one 50 mL tube of 4 mM Rho-kinase inhibitor stock/1.2 mM Wnt signal inhibitor, and one 50-mL tube of 4 mM TGF-β/Activin/Nodal signal inhibitor. The reagents and labware were placed in the appropriate positions. The task of RPE differentiation was initiated, and the robotic operation was performed by LabDroid (medium exchange type I; **Figs. S4B**, **S8**).

On DDays 20 to 32, the users prepared eight 50-mL tubes of RPE differentiation medium (10% KSR; DDays 10 to 25) or RPE maintenance medium (DDays 26 to 32). The reagents and labware were placed in the appropriate positions. RPE differentiation and maintenance were initiated and the robotic operations were performed by LabDroid (medium exchange type II; **Figs. S4C**, **S9**).

*Scoring — sampling*

On DDay 33, the cell plates were exported and the cell culture medium was replaced with fresh RPE maintenance medium. After 24 h (DDay34), the medium was collected for ELISA analysis. The remaining media were aspirated, 2 mL of PBS (-) were added, and were then aspirated again for washing. After that, photographic images were acquired for the calculation of scoring values.

*Scoring — image analysis*

Images were acquired using a digital camera (PSG7X MARKII, Canon Inc., Japan): ISO 500; focal length F=9.00, 50 mm; exposure time, 1/1250 sec. The camera was set in the same position throughout all experiments. The acquired images were automatically processed by filtering with Gaussian blur, subtracting the background, binarizing by thresholding with a constant value, and cropping with a constant pixel value. The colored cell area was then calculated (**Fig. S14**).

*Purification and storage*

Purification of iPSC-RPE cells was conducted using the same protocol described in a study previously reported (*12*). When the RPE colonies reached an appropriate size, the cells were suspended in RPE maintenance medium and kept as a floating culture for about 10 days in a low cell adhesion plate (MS-90600Z, Sumitomo Bakelite Co., Ltd., Japan). Under the microscope, colonies consisting only of black RPE cells were selected. Then, they were transferred to 12-well plates coated with iMatrix, and cultured in RPE adhesion medium/RPE maintenance medium (1:1). Once the RPE cell colonies became attached to the dish, they were cultured in RPE maintenance medium with basic fibroblast growth factor (bFGF), which was changed every 2 to 3 days.

After 10–12 days of cell selection, unsuitable cells were removed, and the cells were passaged. The medium was aspirated and 1 mL of RPE washing solution was added and aspirated again for washing. Then, 0.5 mL of RPE washing solution was added and atypical cells were eliminated using micropipette tips under microscope observation. After the removal process, the medium was aspirated, 1 mL/well of PBS (-) was added and aspirated for washing, and then 0.5 mL of Trypsin-EDTA solution (203-20251, FUJIFILM Wako Pure Chemical Corporation, Japan) was added, followed by incubation at RT (approximately 25°C) and 5% $CO_2$ for 8–10 minutes. Cells were detached by pipetting and collected into a 50-mL tube. After centrifugation (280 × *g*, 25°C, 4 min), the supernatant was removed, and the pellet was resuspended in 1 mL/plate of RPE adhesion medium/RPE maintenance medium (1:1) and filtered through a 40-µm cell strainer (352340, Corning Incorporated, NY, U.S.A.). After counting the cells with a hemocytometer, the cells were seeded into 12-well plates. The medium was changed to RPE maintenance medium with bFGF.

After 1–3 days of cell passage, the medium was aspirated, the cells were washed with 0.5 mL of RPE maintenance medium, and 1 mL of RPE maintenance medium containing 10 ng/mL bFGF and 0.5 µM SB431542 was added. This medium was exchanged every 2–3 days.

The cells were stored when they formed hexagonal shapes after sufficient confluency. For that, the medium was aspirated, 1 mL/well of PBS (-) was added and then aspirated for washing, and 0.5 mL of Trypsin-EDTA solution (203-20251, FUJIFILM Wako Pure Chemical Corporation, Japan) was added, followed by incubation at 37 ˚C and 5% $CO_2$ for 10–15 minutes. After adding > 0.5 mL of RPE adhesion medium, the cells were detached using a cell scraper (MS-93100, Sumitomo Bakelite Co., Ltd., Japan). The cell suspension was filtered through a 40-µm cell strainer (352340, Corning Incorporated, NY, USA) and then centrifuged for 4 min at 280 × *g* to obtain a cell pellet. The pellet was resuspended in 1 mL of RPE adhesion medium/RPE maintenance medium (1:1) and filtered through a 40-μm cell strainer. After counting the cells with a hemocytometer, the cell suspension was centrifuged for 4 min at 280 × *g* to obtain a cell pellet. Then, STEM-CELLBANKER (CB047, Zenoaq Resource Co., Ltd., Japan) was added until a cell concentration of 500,000 cells/0.5 mL/tube, and the cell suspensions were dispensed into cryovials. The cryotubes were placed in a cell freezing container at -80°C for 3–24 h, and then stored at -150°C.

*Initiation of iPSC-RPE stock and recovery culture*

Frozen vials of RPE cells were thawed in a 37°C water bath and suspended in 4.5 mL of RPE adhesion medium. After centrifugation (280 × *g*, 25°C, 4 min), the supernatant was removed and RPE adhesion medium/RPE maintenance medium (1:1) was added. After counting the cells with a hemocytometer, the cells were seeded into 24-well plates (0.5 mL/well).

After 1–3 days of cell seeding, the medium was aspirated, the cells were washed with 0.25 mL of RPE maintenance medium, and 0.5 mL/well of RPE maintenance medium containing 10 ng/mL bFGF and 0.5 μM SB431542 was added. This same type of medium was exchanged every 2–3 days.

Two weeks after seeding, the RPE cells were passaged. Two weeks after cell passage, the RPE cells were used for cell biological validation processes (RT-PCR, ELISA, and immunohistochemistry).

*Validation — RT-PCR*

Total RNA was extracted from transfected cells using RNeasy Micro Kit (#74004, QIAGEN, Germany). First-strand cDNA synthesis was performed on 500-1000 ng of total RNA, using SuperScript III (#18080-044, Thermo Fisher Scientific Inc., MA, USA) according to the manufacturer's instructions. Each mRNA transcript was amplified using PCR with the following primers:

*BEST1* (+), 5′-dTAGAACCATCAGCGCCGTC
*BEST1* (−), 5′-dTGAGTGTAGTGTGTATGTTGG
*RPE65* (+), 5′-dTCCCCAATACAACTGCCACT
*RPE65* (−), 5′-dCCTTGGCATTCAGAATCAGG
*CRALBP* (+), 5′-dGAGGGTGCAAGAGAAGGACA
*CRALBP* (−), 5′-dTGCAGAAGCCATTGATTTGA
*GAPDH* (+), 5′-dACCACAGTCCATGCCATCAC
*GAPDH* (−), 5′-dTCCACCACCCTGTTGCTGTA

*Validation — ELISA*

The collected media were centrifuged (90 × g, 4°C, 1 min), and the supernatant was collected and stored at −80°C. The amount of VEGF contained in the thawed medium was measured using the protocols and reagents from the VEGF Human ELISA Kit (BMS277-2, Thermo Fisher Scientific, USA), and the amounts of PEDF were measured using a Human ELISA Kit (RD191114200R, BioVendor, Czech Republic).

*Validation — Immunohistochemistry*

Cells were washed with PBS (-), fixed in 15% paraformaldehyde for 1 h at RT (approximately 25°C), and stored at 4°C after removal of PFA and addition of PBS (-). After removal of the solutions, cells were treated with 50 μL/well of 0.2% Triton X-100/PBS (-), incubated for 30 min at RT, washed with PBS (-), blocked with 50 μL of Blocking One (03953-95, Nacalai Tesque Inc., Japan), and incubated for 1 h at RT. After removal of the solutions, cells were stained at 4°C o/n in 50 μL of the 1st antibody diluent (rabbit anti-ZO-1, 61-7300, Thermo Fisher Scientific Inc., MA, USA; anti-MITF, mouse anti-MiTF, ab80651, Abcam plc., Britain; antibody diluent, S2022, Agilent Technologies Inc., USA). After removal of the solutions, cells were washed with PBS (-) and then stained at RT for 1 h in 50 μL of the 2nd antibody diluent (Alexa Fluor 546 Goat Anti-mouse IgG, A-11030, Thermo Fisher Scientific Inc., MA, USA; Alexa Fluor 488 Goat Anti-rabbit IgG, A-11034, Thermo Fisher Scientific Inc., MA, USA; antibody diluent, S2022, Agilent Technologies Inc., USA) with DAPI (1 μg/mL, D1206, Thermo Fisher Scientific Inc., MA, USA). After removal of the solutions, cells were washed with PBS (-), and then 50 μL of PBS (-) was added. Images of immunohistochemistry samples were acquired using an IX73 inverted microscope (Olympus, Japan).

*Bayesian optimization module*

When no prior experimental results exist, the Bayesian optimization module generates the next query from random uniform sampling. When past experimental results are available, the Bayesian optimization module generates queries using two components: the Model updater and the Query generator (**Fig. 3C**).

The Model updater updates the surrogate model to predict the experimental results given past experimental results: $D = \{(\boldsymbol{x}_i, y_i)\}_{i=1}^{n}$. We adopted Gaussian process regression (GPR, **Fig. S10**) with the ARD-RBF kernel as the surrogate model to estimate the expected score and confidence level for all unevaluated experimental parameters. Based on the experimental results shown in **Fig. 2E**, the observation noise was assumed to follow a zero-mean Gaussian noise with a variance of 0.0039 at all points in the search space. By using the surrogate model, the Query generator generates the next queries in two steps. In step 1, the Query generator constructs an acquisition function that estimates the expected progress toward the optimal experimental parameter at a given experimental parameter $\boldsymbol{x}$ in the search space. We adopted the Expected improvement (EI) (*31*), a commonly used acquisition function in BO. EI estimates how much improvement over the current best score is expected from each point in the search space. In step 2, by using the acquisition function, the Query generator decides where to evaluate next, and our problem required the simultaneous performance of 48 experiments corresponding to 8 plates x 6 wells in each round. In addition, because the trypsin processing time (DP) is a batch contextual parameter as described herein, a policy function that generates parameter sets taking such structural context into account must be incorporated. Therefore, we developed the Batch Contextual Local Penalization (BCLP) as a policy function to generate multiple points with context in parallel. The BCLP is a batch generation policy that extends the local penalization (*32*) to be applied to cases where complex structural context parameters exist. As shown in **Fig. 3E**, for each value of the contextual parameter DP in ascending order, BCLP iteratively generated the parameter by maximizing and penalizing the acquisition function 48 times to obtain the next experimental parameters $\boldsymbol{X}_{\mathrm{next}}$ for each subsequent well (**Algorithm S2**). In addition, after each round, the more promising KP intervals were reconfigured by calculating the integral value of the acquisition function (**Algorithm S3**). We also replaced the queries that corresponded to the place of the top two pigmented scores in the previous experiments with the parameter of the top two pigmented scores in the previous experiments as a positive control.

*Statistical analysis*

Statistical analyses were performed by Wolfram Mathematica version 11.2.0.0. In this study, $P < 0.05$ was considered significant (*$P < 0.05$, **$P < 0.01$, ***$P < 0.001$, and n.s. = not significant).

*Data and code availability*

All code that supports the findings of this study is available at https://github.com/labauto/LabDroid_optimizer. This code is based on GPyOpt (*33*).

### 2. System Outline

*2.1. A target process of our robotic search*

In this study, the differentiation induction process of iPSC-RPE is the object of the robotic search. This process corresponds to part of the whole process of preparing RPE cells from iPS cells, and to surgical transplantation (**Fig. S1A**). The effect measurement of the robotic search was verified by the score of the colored cell area, and the validation of the robot-produced cells was verified by a cell biological quality control test before transplantation, in a previous clinical study (*12*) (**Fig. S1B**).

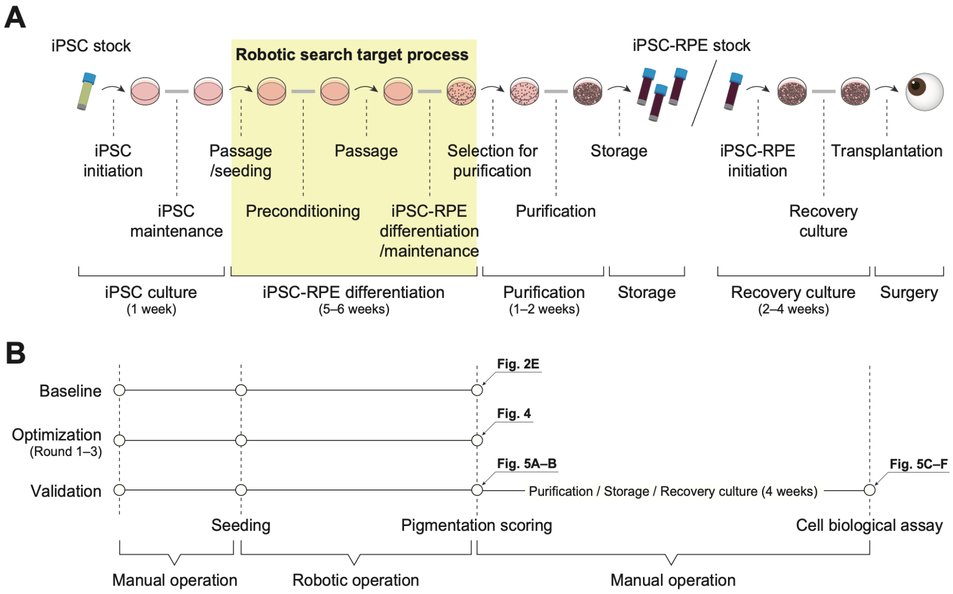


**Fig. S1: Schematic diagram of iPSC-RPE transplantation**

(**A**) Schematic diagram of the process from iPSC stock to iPSC-derived RPE cell transplantation. The steps are roughly divided into iPSC culture, iPSC-RPE differentiation, purification, storage, recovery culture, and surgery. Arrows indicate daily operations, and thick grey lines indicate multi-day operations.

(**B**) Timeline of the baseline, optimization and validation experiments. The baseline and optimization experiments were completed with scoring. The validation experiments, however, were performed by generating an iPSC-RPE stock through a purification process and carrying out a cell biological analysis. The arrows represent the figure number in which the result from that process is displayed.

*2.2. System components*

The system platform is shown in **Fig. S2**. The robotic search using this system starts with the provision of queries for round 1 from the optimizer to the protocol compiler and the provision of the base protocol from the user to the protocol compiler.

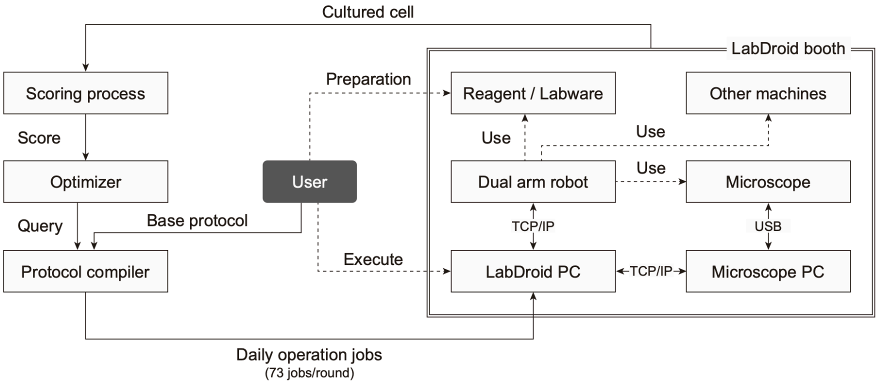


**Fig. S2: System components**

Each rectangle represents a system component. Solid lines represent the movement of data (including cells), and dotted lines represent physical interactions.

### 3. Robotic Components

*3.1. Robotic setup*

LabDroids, including peripheral equipment, were placed inside a booth made of acrylic walls and a stainless steel frame with three fan-filter-units (**Fig. S3A**). LabDroid booths are equipped with humanoid robots and workbenches, as well as the same kind of experimental equipment that humans use. See **Materials and Methods** for the model numbers of the machines and **Fig. S3B–F** for the layout of the equipment.

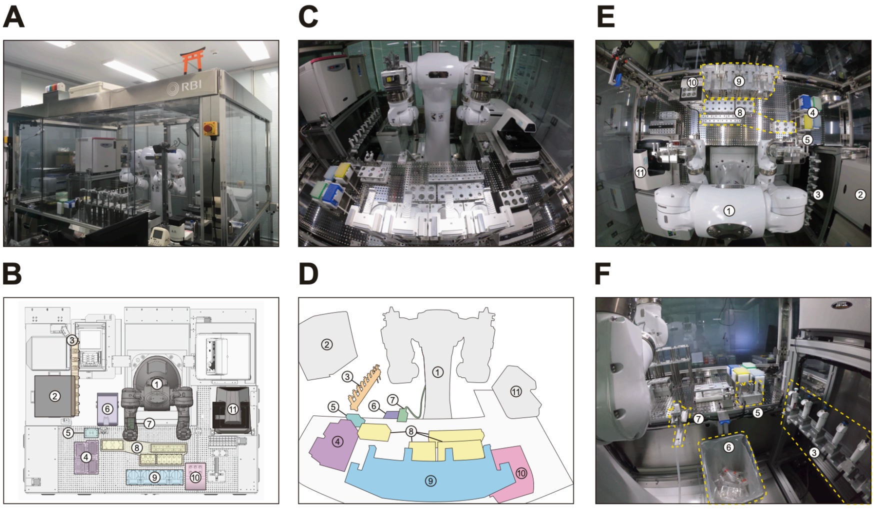


**Fig. S3: LabDroid Maholo booth**

(**A**) Exterior photograph of the LabDroid booth. The LabDroid consists of an acrylic box of W2500 x D2000 x H2200 (mm).

(**B**) Plan view of the LabDroid booth (3D-CAD) and layout of the equipment.

(**C–D**) Front view photograph of the LabDroid booth and schematic drawing of the equipment (these panels are identical to **Fig. 2B**).

(**E**) Top view photograph of the LabDroid booth.

(**F**) A back view photograph of the LabDroid booth.

Components: (1) dual-arm humanoid; (2) $CO_2$ incubator; (3) micropipettes; (4) pipette tips; (5) tip sensor; (6) dustbin; (7) aspirator; (8) tube racks; (9) plate racks; (10) dry bath; and (11) microscope.

*3.2. Robotic workflows*

The program files for operating the LabDroid were created with ProtocolMaker (YASKAWA Electric Corp., Fukuoka, Japan), the GUI software used to operate the LabDroid. The workflow of the operations is shown in **Fig. S4**. Representative screenshots of ProtocolMaker are shown in **Figs. S5–9**.

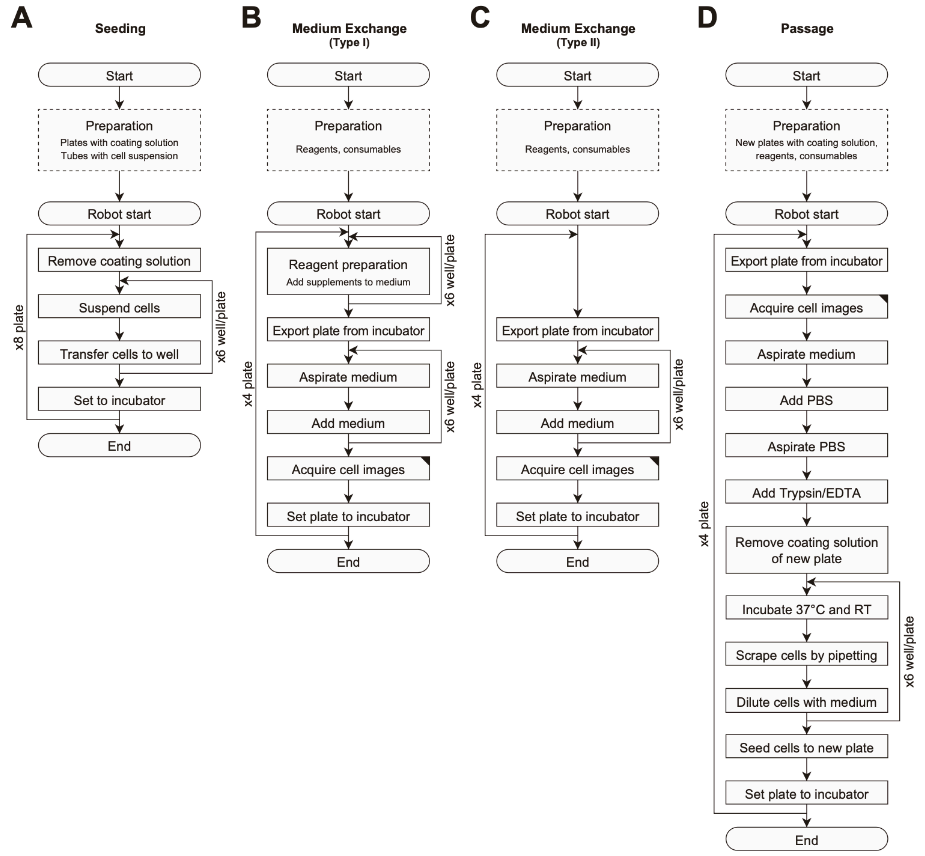


**Fig. S4: Workflows of the experimental operation**

(**A**) Seeding (DDay -7).

(**B**) Medium exchange type I for preconditioning (DDays -6 to -1) and the first part of RPE induction (DDays 1 to 19).

(**C**) Medium exchange type II for the second part of RPE induction (DDays 20 to 25), and RPE maintenance (DDays 26 to 32).

(**D**) Passage (DDay 0).

The dashed line rectangles represent the operations carried out by humans, and solid line rectangles represent the operations carried out by the robot.

A

B

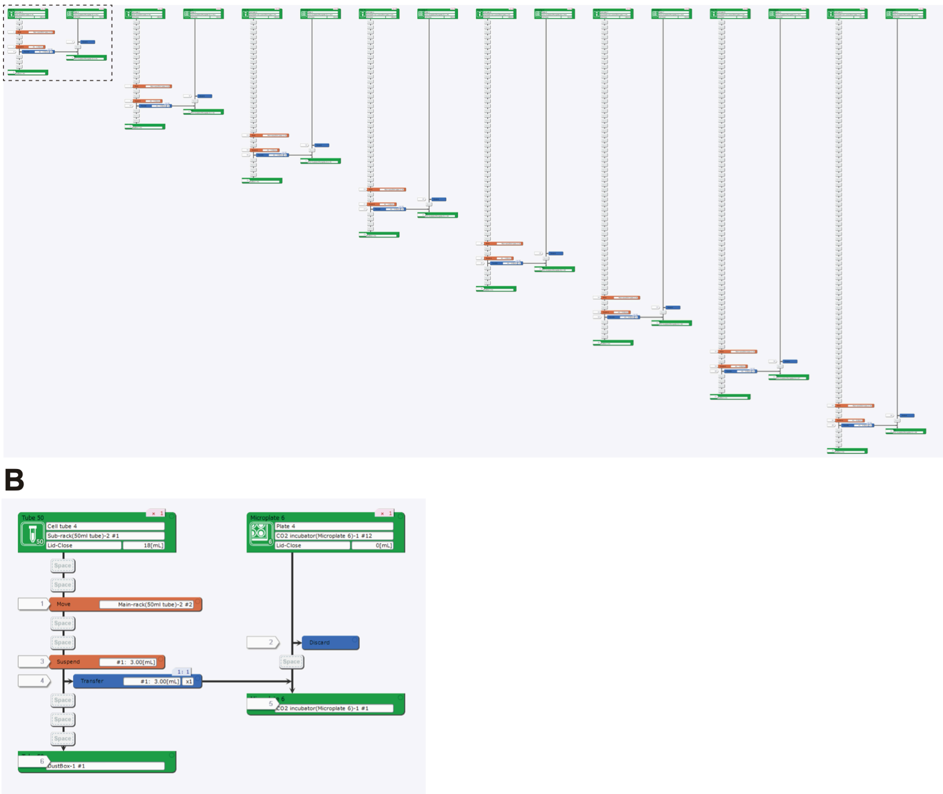


**Fig. S5: Representative LabDroid execution of a seeding experiment (round 3, DDay -7)**

(**A**) Entire image. High-resolution image is available at:
[https://www.dropbox.com/sh/1wy9pvrqml7a3ur/AABKKxfWXNZlBFqxNwvll_Qia?dl=0](https://www.dropbox.com/sh/1wy9pvrqml7a3ur/AABKKxfWXNZlBFqxNwvll_Qia?dl=0)

(**B**) Enlarged image of the dotted rectangle from panel **A**.

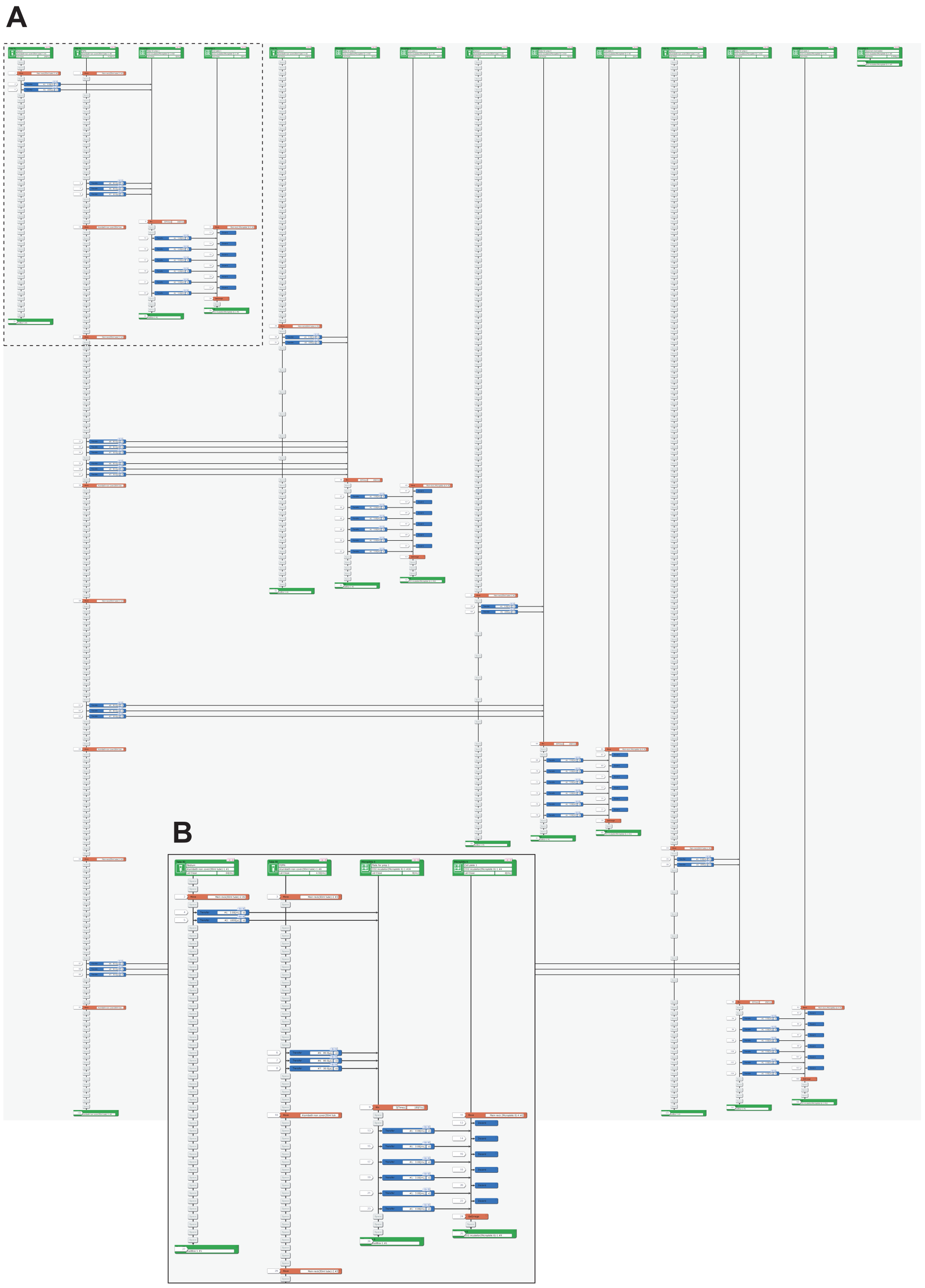

**Fig. S6: Representative LabDroid execution of a preconditioning experiment (round 3, DDay -6, 1st run)**

(**A**) Entire image. High-resolution image is available at:
https://www.dropbox.com/sh/1wy9pvrqml7a3ur/AABKKxfWXNZlBFqxNwvll_Qia?dl=0

(**B**) Enlarged image of the dotted rectangle from panel **A**.

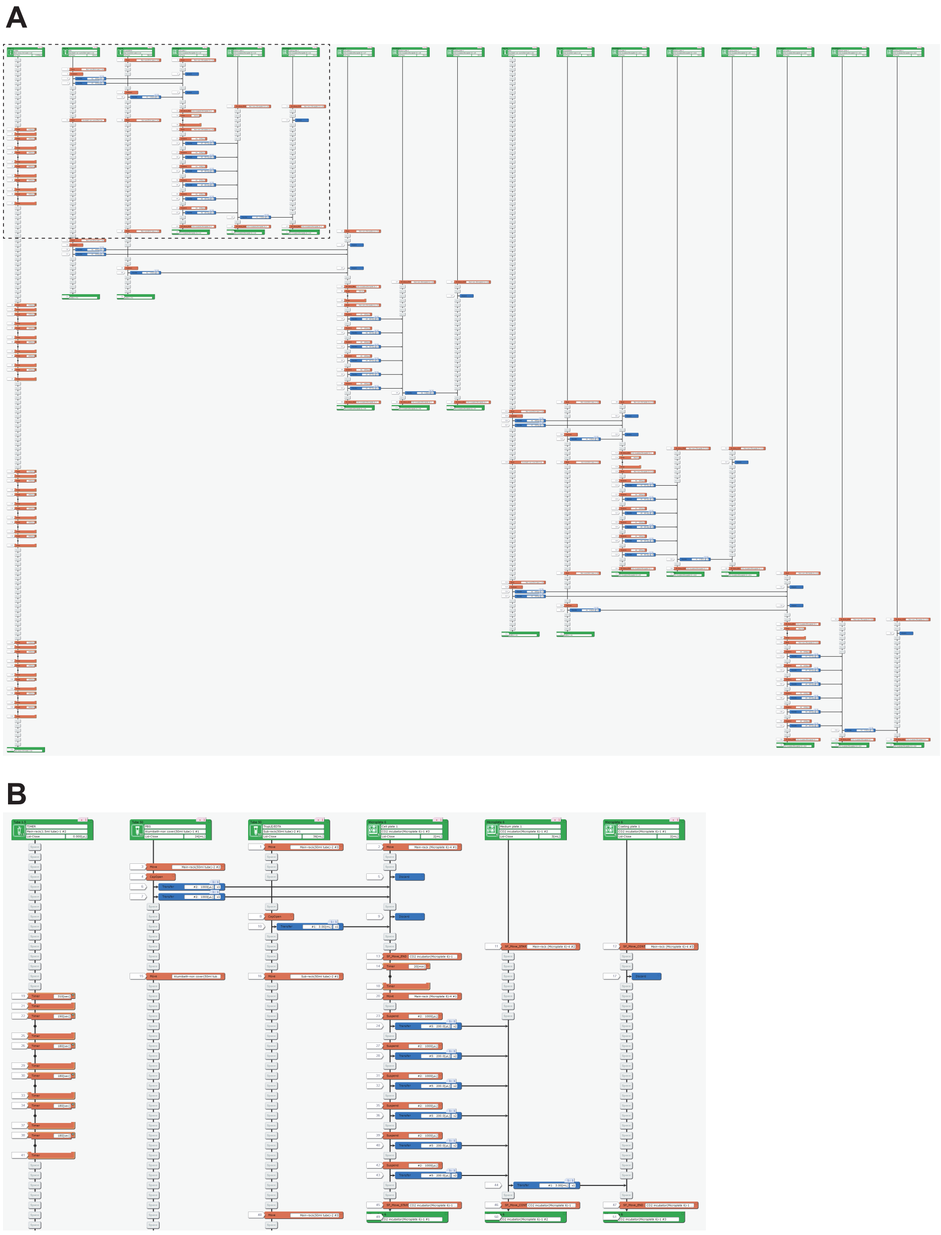

**Fig. S7: Representative LabDroid execution of a passage experiment (round 3, DDay 0, 1st run)**

(**A**) Entire image. High-resolution image is available at:

[https://www.dropbox.com/sh/1wy9pvrqml7a3ur/AABKKxfWXNZlBFqxNwvll_Qia?dl=0](https://www.dropbox.com/sh/1wy9pvrqml7a3ur/AABKKxfWXNZlBFqxNwvll_Qia?dl=0)

(**B**) Enlarged image of the dotted rectangle from panel **A**.

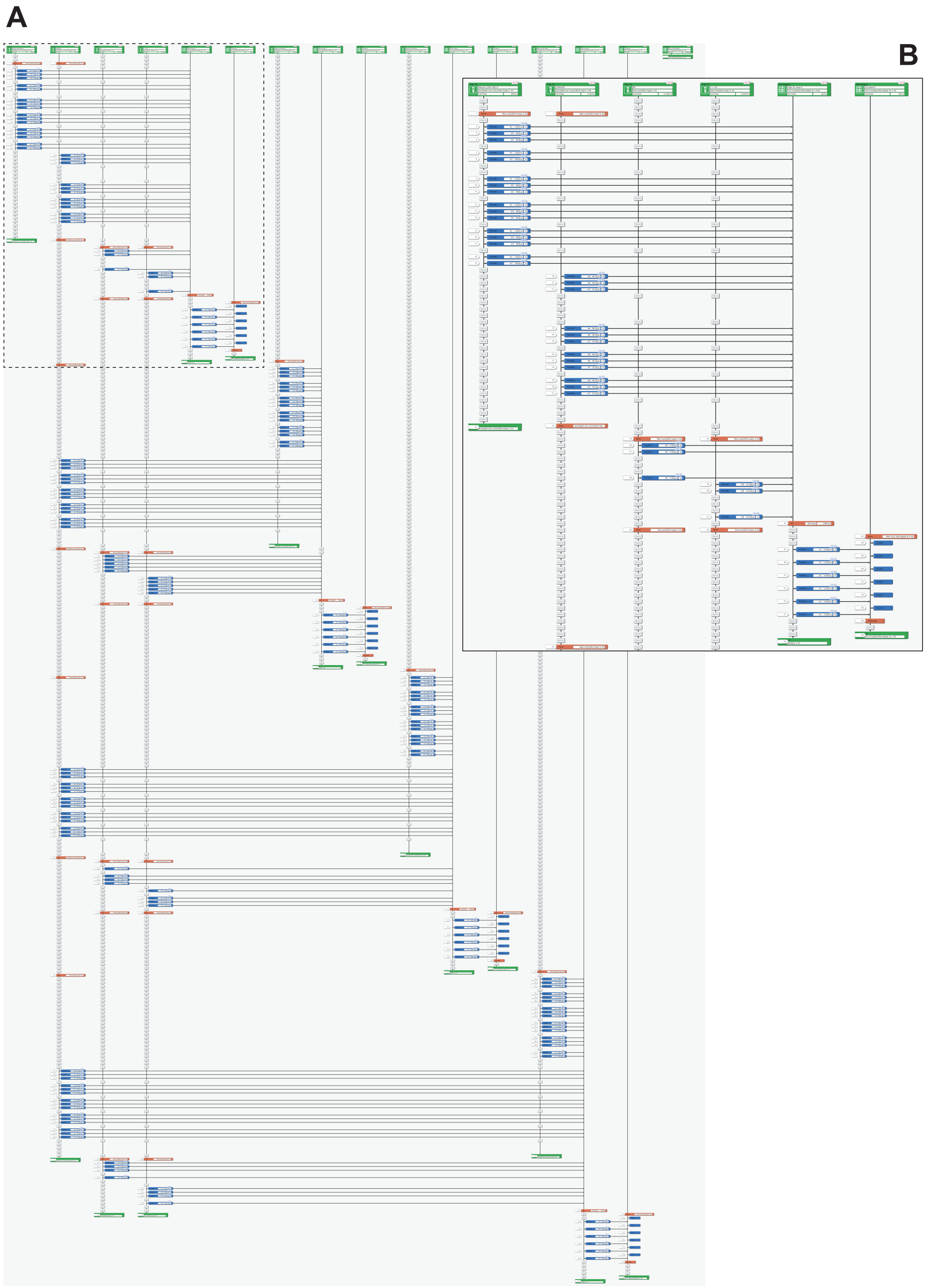

**Fig. S8: Representative LabDroid execution of an RPE differentiation experiment (round 3, DDay 10, 1st run)**

(**A**) Entire image. High-resolution image is available at:
https://www.dropbox.com/sh/1wy9pvrqml7a3ur/AABKKxfWXNZlBFqxNwvll_Qia?dl=0

(**B**) Enlarged image of the dotted rectangle from panel **A**.

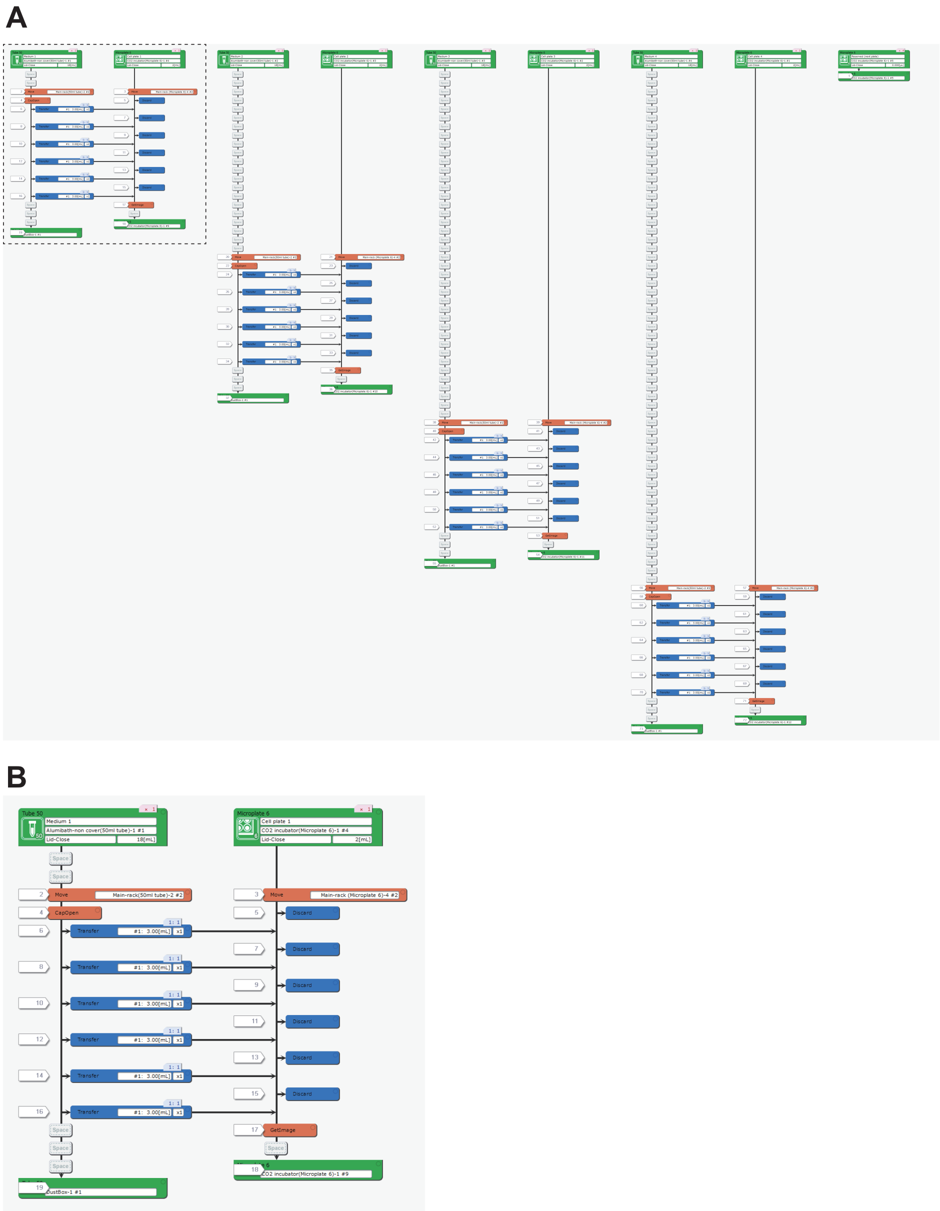


**Fig. S9: Representative LabDroid execution of an RPE maintenance experiment (round 3, DDay 32, 1st run)**

(**A**) Entire image. High-resolution image is available at:
https://www.dropbox.com/sh/1wy9pvrqml7a3ur/AABKKxfWXNZlBFqxNwvll_Qia?dl=0

(**B**) Enlarged image of the dotted rectangle from panel **A**.

*3.3. Micropipette combination for the transfer of a wide range of volumes with LabDroid*

Nine manual micropipettes were registered in the LabDroid used in this study. Because the movement to change the individual pipette volume is inefficiently implemented by the LabDroid, nine micropipettes of different volumes were prepared and combinations of them were used for the transfer of a wide range of volumes. In this study, the experiments were performed using 3000, 1000, 450, 300, 200, 80, 30, 10, and 5 µL pipettes, and the pipettes were combined up to three times. Because of this implementation constraint, approximations may have to be used for the *PC* and *KP* parameters; when the required quantities of each reagent are not feasible because of the above constraints, the closest feasible quantity is used. The feasible quantities and their pipette combinations are listed in **Table S1**.

### 4. Informational Component

*4.1. Compiling experimental design table to robot protocols*

The iPSC-RPE differentiation-inducing cultures performed in this study required a total of 73 robot-executed runs per round: one run for seeding, 12 runs for preconditioning, 2 runs for passage, and 58 runs for medium exchange after passage. The term "run" refers to a group of experiments in which a robot is set to work. For seeding, eight plates were used in one batch on one day; and on other days, two runs per day were conducted as one batch of four plates. Because 24 or 48 conditions were set for each of the 73 robot-executed runs, the manual job generation by humans was considered to be a cause of the frequent human error. In this study, we prepared a spreadsheet with the experimental design and a robot job file without parameters (base protocol), and built an environment to automatically generate 71 robot-executed runs (excluding passage) by inserting parameters according to the spreadsheet (**Fig. S2**).

*4.2. Parameterization of iPSC-RPE protocol*

The addition of FGFRi to the medium during preconditioning significantly increased the efficiency of induction of differentiation into RPE cells (*13*). The unit of *PC* is nM, and it takes values from 0 to 505, while *PP* is the number of days FGFRi is added, taking values from 1 to 6. For example, *PC* = 300 and *PP* = 2 indicates that 300 nM of FGFRi is added to DDays -2 and -1, and not from DDay -6 to -3.

*DS* was adopted because shear stress affects cell properties (*34*), and *DL* was adopted because the number of cells at passage is empirically known to contribute to the efficiency of differentiation induction. *DS* takes integer values from 10 to 100, indicating the speed at which the micropipette syringe is pressed (unit: mm/sec); *DL* takes two integer values, one long and one short, indicating the bottom surface area to be pipetted. We adopted *DP* at passage because of the constraints in the configuration of the LabDroid, which required different fixed values. *DP* takes integer values of 5, 8, 11, 14, 17, 20, and 23 to indicate the number of minutes that the trypsin solution was added and incubated at room temperature after incubation at 37 °C for 20 min.

The three chemical supplements are Rho-kinase inhibitor, TGF-β/Activin/Nodal signal inhibitor, and Wnt signal inhibitor, all of which are known to affect the culture and differentiation of iPS cells (*35*, *36*). KSR was added to the medium at a 20% concentration on DDay 0 and was progressively reduced to 10%. Under standard conditions, KSR is added in the following manner: 20% on DDays 0-4, 15% on DDays 5-7, and 10% from DDay 8, suggesting that this rate of reduction is important for the induction of RPE differentiation (*13*). The *KP* takes values from 1 to 19 and the concentration of KSR is decreased linearly daily until KSR becomes 10% on the DDay of the *KP* value. For example, when *KP* = 4, KSR will be 20% on DDay 0, 17.5% on DDay 1, 15.0% on DDay 2, 12.5% on DDay 3, and 10% on DDay 4. *3P* takes integer values from 1 to 19, and indicates the day in which the three chemical supplements are included in the differentiation-inducing medium from DDay 0 to the DDay of that *3P* value.

*4.3. Gaussian process overview*

In Bayesian optimization, the Gaussian process (GP) $p(f) = \mathcal{GP}(\mu, \sigma; k)$ is typically used as the surrogate model to approximate the I/O of the objective function $f$. The GP is characterized by three components: a mean function $\mu$, a variance function $\sigma$, and a kernel function (positive-value covariance function) $k$. In case of past experimental data $D = \{(\boldsymbol{x}_i, y_i)\}_{i=1}^{n}$ of $n$ observations, and Gaussian prior $\mathcal{GP}(0, \sigma; k)$, Gaussian posterior is then given in $\mathcal{GP}(\mu_n, \sigma_n; k)$, where

$$\mu_n(\boldsymbol{x}) = \boldsymbol{k}_n(\boldsymbol{x})^T \left(\boldsymbol{K}_n + \sigma^2 \boldsymbol{I}\right)^{-1} \boldsymbol{y}_n$$

$$\sigma_n^2(\boldsymbol{x}) = k(\boldsymbol{x}, \boldsymbol{x}) - \boldsymbol{k}_n(\boldsymbol{x})^T \left(\boldsymbol{K}_n + \sigma^2 \boldsymbol{I}_n\right)^{-1} \boldsymbol{k}_n(\boldsymbol{x})$$

$\mu_n(\boldsymbol{x})$ is the mean of Gaussian process posterior function, and $\sigma_n^2(\boldsymbol{x})$ is the variance of Gaussian process posterior function. In this work, we assume that the kernel function $k$ is stationary. $\sigma^2$ is the observation noise of the experiment.

**Fig. S10** shows an example of a Gaussian process prior to being updated to a Gaussian posterior using observations. The uncertainty of GP distribution decreases around the observed points and increases further away from observations. For a more comprehensive explanation of Gaussian process regression, see (*37*).

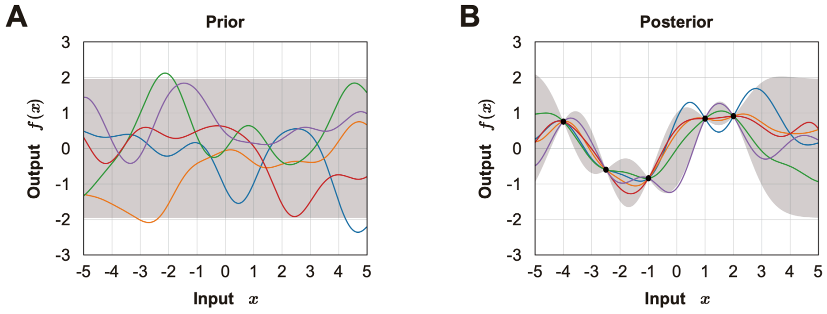


**Fig. S10: Demonstration of how Gaussian process regression updates a Bayesian posterior**

Test of a Bayesian posterior updated in a Gaussian process (GP).

(**A**) Sample paths from the zero-mean GP prior.

(**B**) Sample paths from the GP posterior after executing some experiments (indicated by black dots).

The gray shaded area represents the pointwise mean plus and minus twice the standard deviation for each input value.

*4.4. Bayesian optimization overview*

Bayesian optimization is a common black-box optimization method used to determine the input parameter $\boldsymbol{x}$, which maximizes the objective function $f$ with as few executions as possible using iterative experiment-observation loops.

$$\boldsymbol{x}_{\max} = \arg\max_{\boldsymbol{x} \in \mathcal{X}} f(\boldsymbol{x})$$

**Fig. S11** shows a toy example of how sequential Bayesian optimization generates the next experimental parameter. Bayesian optimization iteratively generates the following experimental parameters in three steps: construction of a surrogate model (e.g., Gaussian process regression) for past experimental results (left column), definition of an acquisition function $\alpha(\boldsymbol{x}; D)$ (e.g., expected Improvement), and optimization of the acquisition function to obtain the next experimental parameter (right column). For a more comprehensive explanation and application of Bayesian optimization, see (*38*, *39*).

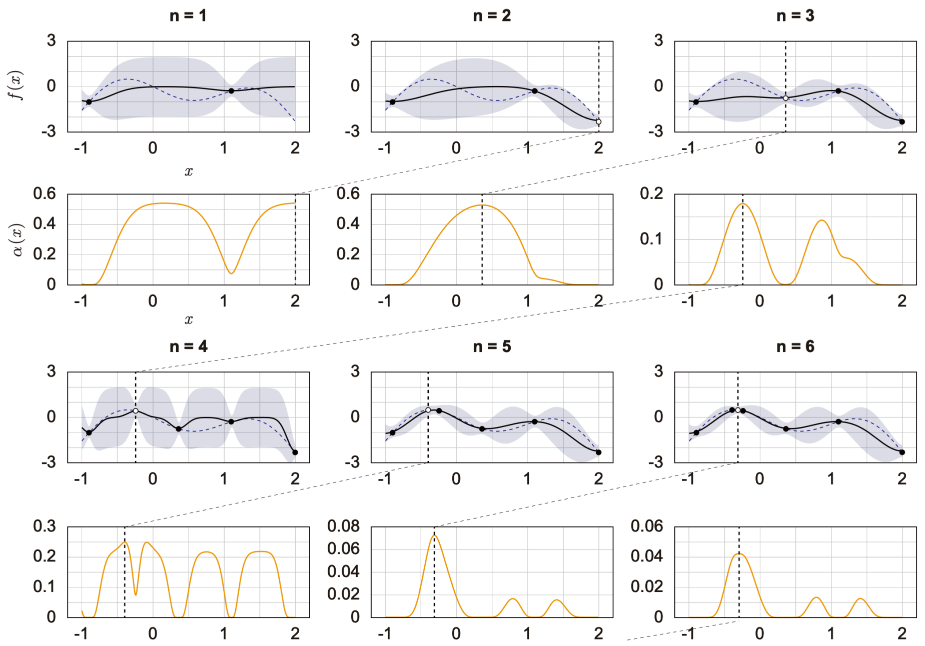


**Fig. S11: Demonstration of Bayesian optimization generation of experimental parameters**

Sequential Bayesian optimization was tested by determining the maximum of a one-dimensional toy objective function. This figure illustrates the Bayesian optimization procedure over several iterations. The objective function is indicated by a purple dashed curve on the left side of each plot. The past experimental results are indicated by dots, and the newest experimental results are indicated by white dots. The GP posterior mean function is indicated by a black line, and covariance intervals are indicated by purple shaded areas. The plots on the right side show the acquisition functions in the orange curves. The value of the acquisition function is high where the model predicts a high objective (exploitation) and where the prediction uncertainty is high (exploration). The black vertical dashed lines show the place of acquisition max, a factor of importance to be tested in a subsequent experiment.

#### 4.5. Bayesian optimization for the iPSC-RPE differentiation protocol

In the batch Bayesian optimization for the iPSC-RPE differentiation protocol, we generated the next experimental queries for each round in four steps (**Algorithm S1**). First, we constructed a Gaussian process posterior using a past experimental dataset. When no prior experimental results existed, we generated the next query from random uniform sampling. Next, using the Gaussian process posterior, we obtained an acquisition function. In this study, we used the expected improvement (*31*) as the acquisition function. Third, using the acquisition function, we generated a next experimental parameter set $\boldsymbol{X}_{\text{next}}$ using a policy function. In this study, we used batch contextual local penalization (BCLP) as the batch generation policy. Finally, after executing experiments on the parameter set $\boldsymbol{X}_{\text{next}}$, we computed the optimal context for the Detach trypsin Period $x_{DP}$ for the next round.

In Gaussian process regression, we assume the ARD-RBF kernel function $k$ as follows:

$$k_{\text{ARD-RBF}}\left(\boldsymbol{x},\boldsymbol{x}';\boldsymbol{\theta}\right)=\exp\left(-\frac{1}{2}\sum_{m=1}^{d}\left(\frac{\boldsymbol{x}-\boldsymbol{x}'}{l_m}\right)^2\right)$$

$\boldsymbol{\theta}=\{l_m\}_m^d$ represents d-dimensional length scale hyper-parameters that were optimized using the maximum-likelihood estimation in every GP fitting.

Based on the experimental results in **Fig. 2E**, the observation noise was assumed to follow a normal distribution with a variance of 0.039, at all points in the search space.

In this work, we used the expected improvement as an acquisition function. The expected improvement estimates how much improvement over the current best score $y_{max}$ is expected from each one of the input parameters $\boldsymbol{x}$ in the search space, as shown in the following form:

$$EI(\boldsymbol{x};D)=(y_{max}-\mu_n(\boldsymbol{x}))\,\Phi(Z)+\sigma_n(\boldsymbol{x})\phi(Z)\quad,\quad Z=\frac{y_{max}-\mu_n(\boldsymbol{x})}{\sigma_n(\boldsymbol{x})}$$

$D$ is the set of the past experimental results. $\boldsymbol{\Phi}$ is the standard normal cumulative distribution function, and $\phi$ is the standard normal probability density function.

In the policy function (see **Fig. 3E**), BCLP iteratively generated the parameter for each value of the contextual parameter *DP* in ascending order by maximizing and penalizing the acquisition function 48 times to obtain the next experimental parameters $\boldsymbol{X}_{\text{next}}$. Here we show the algorithm that generates $\boldsymbol{X}_{\text{next}}$ (**Fig. 3E, Algorithm S2**). Starting from plate1, well1, BCLP fixes the value of the DP corresponding to a given well number and then maximizes the acquisition function to generate an experimental parameter for the well. The penalization of the acquisition function is then performed using a hammer function to the point where the experimental parameter is generated. Similarly, for the next well, an experimental parameter is generated by maximizing the penalized acquisition function and is penalized using the hammer function. By repeating such maximization-penalization loops, BCLP generates the next experimental parameters for each subsequent well.

In **Algorithm S2**, the function $\varphi(\boldsymbol{x};\,\boldsymbol{x}_j)$ is a local penalizer of $\alpha(\boldsymbol{x};\,\mathcal{D}\,)$ at $\mathrm{x}_j$ such that:

$$\varphi\left(\boldsymbol{x};\boldsymbol{x}_j,\hat{L}\right)=\frac{1}{2}\operatorname{erfc}(-z)$$

where

$$z=\frac{1}{\sqrt{2\sigma_n^2\left(\mathbf{x}_j\right)}}\left(\hat{L}\left\|\mathbf{x}_j-\mathbf{x}\right\|-\hat{M}+\mu_n\left(\mathbf{x}_j\right)\right)$$

for $erfc$ the complementary error function, $\hat{M}=\max_i\{\mathbf{y}_i\}$ is the best score in the past experiments and $\hat{L}=\max_{\mathcal{X}}\|\mu_{\nabla}(\boldsymbol{x})\|$ is an approximated Lipschitz constant. Both $\hat{M}$ and $\hat{L}$ were calculated in each round. We used $g(z)=z$ when the acquisition function $\alpha(\boldsymbol{x};\,\mathcal{D})$ was positive and the $g(z)=softplus(z)$ elsewhere (*32*).

For the contextual parameter $x_{DP}$, contexts on a fixed trypsin processing time $c_{DP,t,i}$ are assigned for each well depending on the round number $t$ and the well number $i$, because of the implementation constraints of the protocol. After each round, the context on the $x_{DP}$ is reconfigured as a variable. In the initial state (round $t=1$), trypsin treatment time was assigned to the smallest well number (well 1) for eight minutes, and then it increased by three minutes for each additional well number.

$$\boldsymbol{c}_{DP,1}:=(c_{DP,1,1},c_{DP,1,2},c_{DP,1,3},c_{DP,1,4},c_{DP,1,5},c_{DP,1,6})=(8,11,14,17,20,23)$$

In the next round, the context on $x_{DP}$ was allowed to move back and forth $\Delta c=3$ in parallel while maintaining a three minute interval processing time between the different wells. Thus, the context in round 2 $\boldsymbol{c}_{DP,2}$ will be $\boldsymbol{c}_{DP}$ or $\boldsymbol{c}_{DP}^{-}$ or $\boldsymbol{c}_{DP}^{+}$ as follows:

$$\boldsymbol{c}_{DP}:=\boldsymbol{c}_{DP,1}=(8,11,14,17,20,23)$$
$$\boldsymbol{c}_{DP}^{-}:=\boldsymbol{c}_{DP,1}-\Delta c=(5,8,11,14,17,20)$$
$$\boldsymbol{c}_{DP}^{+}:=\boldsymbol{c}_{DP,1}+\Delta c=(11,14,17,20,23,26)$$

Using the flow shown in **Algorithm S3** in each round, we could choose whether the context in the next round $\boldsymbol{c}_{DP,t+1}$ will be $\boldsymbol{c}_{DP}$, $\boldsymbol{c}_{DP}^{-}$ or $\boldsymbol{c}_{DP}^{+}$, based on the past experimental data.

We performed a regression with GPR in each round and defined the acquisition function EI based on the regressions, deriving the integral value of the EI for $x_{DP}$ for each different candidate interval of $x_{DP}$ (either at 5 min, 8 min, or 11 min start), respectively. When any derived integrals were improved by 5% or more compared to the integrals in the interval used in the current round, the interval for the *DP* for the next round was run in the interval that showed the largest integral value among the candidate intervals.

**Algorithm S1: Batch Bayesian optimization for the iPSC-RPE differentiation protocol**

Input: The search space $\chi$, GP prior ($\mu_0$, $\sigma_0$, $k$ ), number of rounds $M$, number of Plates $P$, number of Wells $W$, Dataset $\mathcal{D}= \{\boldsymbol{x}_i,\ \mathrm{y}_i\ \}_{i=1}^{n}$

for $t$=1 to $M$ do

1. Construct GP posterior ($\mu_t$, $\sigma_t$, $k$ ) using $\mathcal{D}$.
2. Get the acquisition function $\alpha(\boldsymbol{x};\ \mathcal{D})$.
3. Generate a experiment parameter set $\boldsymbol{X}_{next}$ using the policy function.

Execute the experiments $f(\boldsymbol{X}_{next})$

Append the experiment results to past data $\mathcal{D} = \mathcal{D} \cup \{(\boldsymbol{X}_{next},\ f(\boldsymbol{X}_{next}))\}$.

4. Compute optimal context $\boldsymbol{c}_{DP}$ on Detatch trypsin Period in the next experiment.

end

**Algorithm S2: The policy function for the iPSC-RPE differentiation protocol**

Input: The acquisition function $\alpha(\boldsymbol{x};\ \mathcal{D})$, number of Plates $P$, number of Wells $W$

Output: The next experiment parameter set $\boldsymbol{X}_{next} = \{(\boldsymbol{x}_{t,p,w})\}_{(p,w)=1}^{(P,W)}$

1. Calculate utility functions from the acquisition function $\alpha(\boldsymbol{x},\ \mathcal{D})$

$\tilde{\alpha}_0(\boldsymbol{x};\ \mathcal{D}) \leftarrow g(\alpha(\boldsymbol{x};\ \mathcal{D}))$

$\tilde{\alpha}(\boldsymbol{x};\ \mathcal{D}) \leftarrow \tilde{\alpha}_0(\boldsymbol{x};\ \mathcal{D})$

2. Generate next experiment parameters $\boldsymbol{X}_{next} = \{(\boldsymbol{x}_{t,p,w})\}_{(p,w)=1}^{(P,W)}$ in Maximization-Penalization loop

for $p$=1 to $P$ do

for $w$=1 to $W$ do

1. Maximization-step: $\boldsymbol{x}_{t,p,w} \leftarrow \arg\max_{x\in\chi}\ \{\ \tilde{\alpha}\ (\boldsymbol{x};\ \mathcal{D})\ \}$
2. Penalization-step: $\tilde{\alpha}(\boldsymbol{x};\ \mathcal{D}) \leftarrow \tilde{\alpha}_0(\boldsymbol{x};\ \mathcal{D}) \prod_{(k,h)=1}^{(p,w)} \varphi(\boldsymbol{x}\ ;\ \boldsymbol{x}_{t,k,h},\ \hat{L})$

end

end

**Algorithm S3: Detachment trypsin period adjustment on the iPSC-RPE differentiation protocol**

Input: The acquisition function $\alpha(\boldsymbol{x}; \mathcal{D})$, current DP context $\boldsymbol{c}_{DP,t}$, context shift width $\Delta$c
Output: The next DP context $\boldsymbol{c}_{DP,\ t+1}$

1. Candidates of DP context ranges for the next round. (In this study, $\Delta$c= 3 min)

$\boldsymbol{c}_{DP} \leftarrow \mathrm{c}_{DP,t}$
$\boldsymbol{c}_{DP}^{-} \leftarrow \mathrm{c}_{DP,t}$ - $\Delta$c
$\boldsymbol{c}_{DP}^{+} \leftarrow \mathrm{c}_{DP,t} + \Delta\mathrm{c}$

2. Calculate values $V$, $V^{-}$, $V^{+}$ that accumulate $\alpha(\boldsymbol{x}, \mathcal{D})$ on each context ranges $\boldsymbol{c}_{DP}$, $\boldsymbol{c}_{DP}^{-}$, $\boldsymbol{c}_{DP}^{+}$

$V = \sum_i \int_{\chi} \alpha(\boldsymbol{x}; \mathcal{D}, x_{DP} = c_{DP,\ i})$

$V^{-} = \sum_i \int_{\chi} \alpha(\boldsymbol{x}; \mathcal{D}, x_{DP} = c_{DP,\ i}^{-})$

$V^{+} = \sum_i \int_{\chi} \alpha(\boldsymbol{x}; \mathcal{D}, x_{DP} = c_{DP,\ i}^{+})$

3. Calculate ratios $R^{-}$, $R^{+}$ between each values defined above.

$R^{-} = V^{-}/V$
$R^{+} = V^{+}/V$

4. Choose the next DP context $\boldsymbol{c}_{DP,t+1}$ in following rules.

if ( $\max R^{-}, R^{+} < 1.05$ ) then
| $\boldsymbol{c}_{DP,\ t+1} \leftarrow \mathrm{c}_{DP}$
end
else if ($R^{-} > R^{+}$) then
| $\boldsymbol{c}_{DP,\ t+1} \leftarrow \mathrm{c}_{DP}^{-}$
end
else if ($R^{-} \leq \mathrm{R}^{+}$) then
| $\boldsymbol{c}_{DP,\ t+1} \leftarrow \mathrm{c}_{DP}^{+}$
end

*4.6. Testing optimization in simulation*

Bayesian optimization was tested by optimizing the seven dimensional toy testing function shown in **Fig. S12** under different conditions. **Fig. S13A** shows a comparison between the performance of batch Bayesian optimization and that of random search when the scale of the observation noise on the testing function was changed. The testing function with different noise (SD = 0.000, 0.064, 0.400) was optimized using batch Bayesian optimization with batch contextual local penalization (BCLP) and uniform random sampling, using 8 plates × 6 wells = 48 queries per round. In each condition, we performed 18 independent experiments. When the observation noise was not present (SD = 0.000) or was sufficiently small (SD = 0.064), as in the proposed system shown in **Fig. 2E,** BCLP shows better convergence performance compared to the random search. When the scale of the observation noise was relatively large (SD = 0.400), the performance of BCLP was only as good as that of the random search case.

**Fig. S13B** compares the performances of BCLP when the batch size was changed. For the benchmark function (noise SD = 0.064) , optimization was performed using a different number of plates (Np) per round. In each condition, we performed n = 18 independent experiments and showed that the convergence performance improved as the number of plates (the batch size) increased.

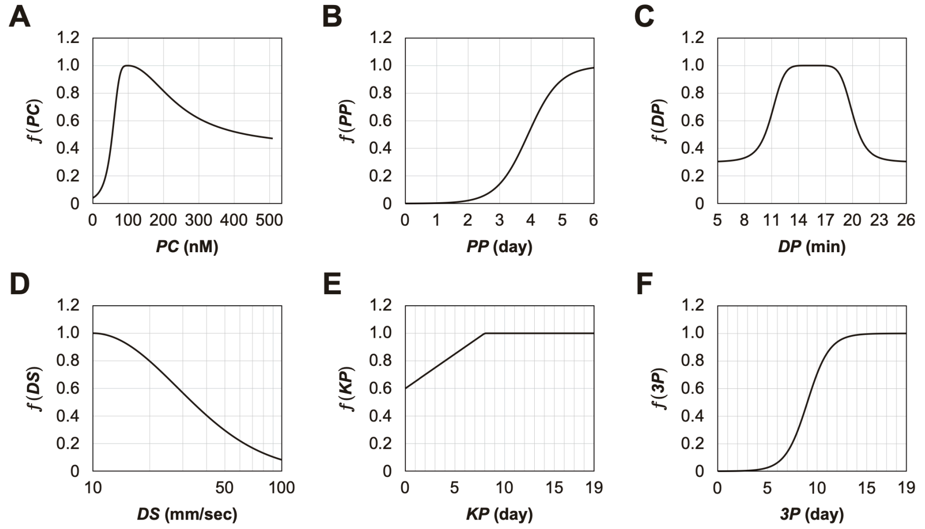


#### Fig. S12: Toy testing function based on domain knowledge

An approximate prediction of how much the value of each variable affects the score based on domain knowledge before performing a series of optimizations. The toy testing function was simply multiplied by the predicted function for each of the variables below, $f(x_{PC}, x_{PP}, x_{DS}, x_{DL}, x_{DP}, x_{KP}, x_{3P}) = f_{PC}(x_{PC})f_{PP}(x_{PP})f_{DS}(x_{DS})f_{DL}(x_{DL})f_{DP}(x_{DP})f_{KP}(x_{KP})f_{3P}(x_{3P})$.

(**A**) Prediction of preconditioning (FGFRi) concentration (PC) response. It increased from 0 nM, reached a maximum at 100 nM, decreased after 100 nM, and approached 0.5.

(**B**) Prediction of the preconditioning period (PP) response: On Day 1 it was 0, it increased monotonically and reached a maximum value on Day 6.

(**C**) Prediction of the detachment trypsin period (DP) response: an optimal value between 13 and 20 min was expected.

(**D**) Prediction of the detachment pipetting strength (DS) response: it achieved a maximum value at 10 mm/s and then decreased monotonically. Although an important parameter, the detachment pipetting length (DL) is always assumed to be 1 because of the difficulty of prediction.

(**E**) Prediction of the KSR period (KP) response: it was expected to increase monotonically until Day 8, and to be stationary thereafter.

(**F**) Prediction of the three-supplement period (3P) response: it was expected to take a non-zero value and increase monotonically from around Day 3 onwards.

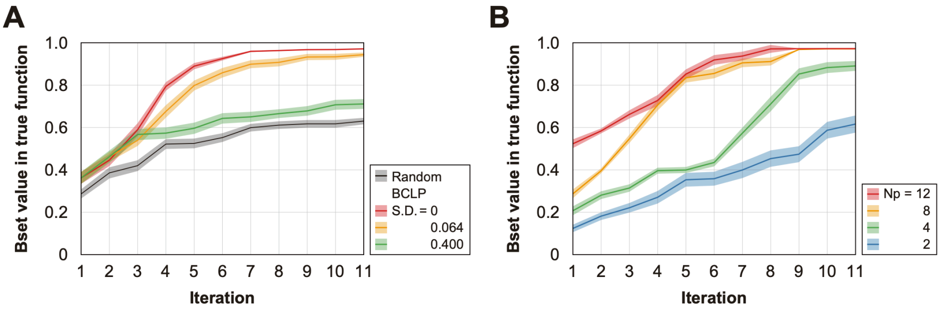


**Fig. S13: Preliminary testing of the Bayesian optimization under different conditions**

The vertical axis shows the value, and the horizontal axis shows the number of circles. The blue horizontal line represents the ground truth (optimal value). In each series, three experiments were conducted independently, and the score of the query with the highest evaluation score in the true function among the queries that had appeared in a certain round was plotted. The error bars represent the standard error (SEM) in each round.

(**A**) The red series shows the results of batch Bayesian optimization (BCLP) with no observation noise. The yellow series shows the results of BCLP with a Gaussian noise SD = 0.064. The green series shows the BCLP results with a high Gaussian noise SD = 0.4. The black series represents the results of random sampling. Compared to random sampling, Bayesian optimization improved the convergence performance and converged to the optimal solution when the observation noise was sufficiently small. Black, random; green, BCLP (SD = 0.4); orange, BCLP (SD = 0.064); red, BCLP (SD = 0). The shaded area represents the SEM image.

(**B**) Comparison of BCLP performance when the batch size was changed. For the benchmark function (noise SD = 0.064), optimization was performed using a different number of plates (Np) per round. The blue series represents Np = 2, the green series represents Np = 4, the orange series represents Np = 8, and the red series represents Np = 16. As the number of plates (the batch size) increased, BBO tended to converge to an optimal solution with fewer rounds. Blue, Np = 2; green, Np = 4; orange, Np = 8; red, Np = 16. The shaded area represents the SEM image.

### 5. Supplementary Results

*5.1. Image analysis for scoring*

Cells differentiating into RPE produce melanin, which causes them to turn brown. Therefore, the area ratio of the total number of pigmented cells on DDay 34 was used to estimate the differentiation induction efficiency and obtain evaluation scores, following the example of previous studies (*10*, *13*). Images were acquired using a digital camera (PSG7X MARKII, Canon Inc., Japan): ISO 500; focal length F=9.00, 50 mm; exposure time, 1/1250 sec. The camera was set in the same position throughout all experiments. The acquired images were automatically processed by filtering with Gaussian blur, subtracting the background, binarizing by thresholding with a constant value, and cropping with a constant pixel value. The colored cell area was then calculated (**Fig. S14**). All images from the robotic search experiment are shown in **Figs. S15–19**.

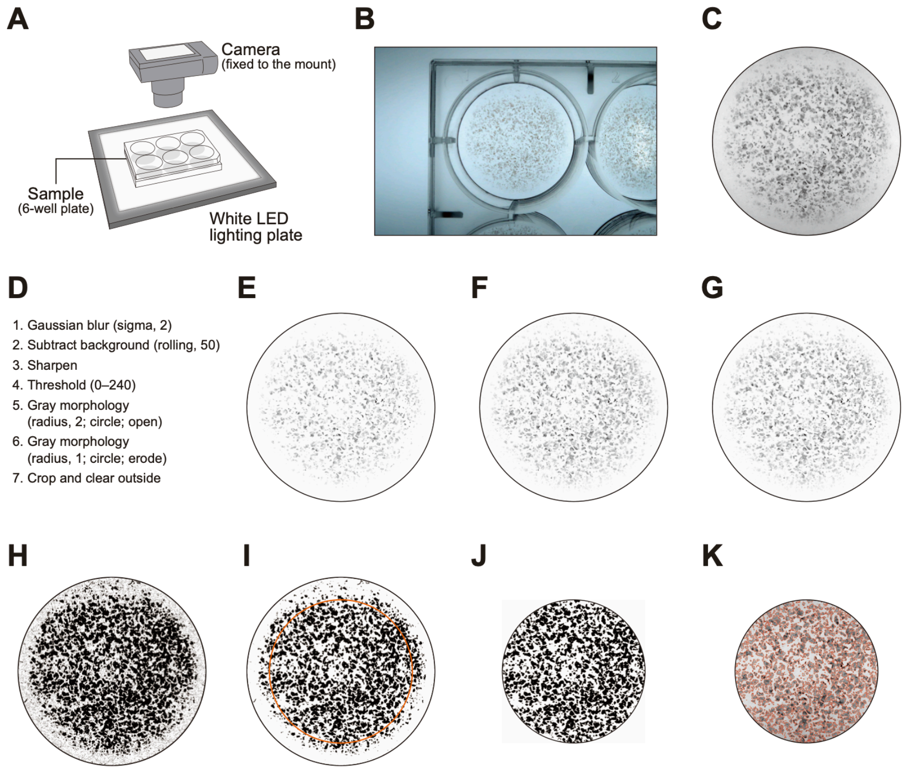


**Fig. S14: Image processing to calculate pigmentation scores**

(**A–C**) 6-well plates were placed on top of a white LED lighting plate and photographed with a camera fixed to the mount (**A**). The raw images acquired (**B**) were cropped in a circle to the size of the bottom of the well (**C**).

(**D**) Workflow of image processing performed using ImageJ/Fiji macro. The background was removed in steps 1–3, binarization in step 4, and noise in steps 5 and 6. Since it is empirically known that differentiation-inducing cells are less likely to grow near the sides of the wells, only the central portion was cropped (step 7) and the scores were subsequently calculated.

(**E–J**) Examples of processed images. The images shown are samples from well 1 of plate 1 in the baseline experiment. Images after Gaussian blur (**E**), background subtraction (**F**), sharpening (**G**), binarization by thresholding (**H**), mathematical morphology processing (**I**), crop, and clear outside (**J**) processing. The orange circle in panel **I** represents the crop area in step 7. The pigmentation score was calculated as the area of the black region in panel **J**.

(**K**) Merged image of the image before processing (corresponding to panel **C**) and the area determined to be pigmented (corresponding to panel **J**, red frame).

All images shown in this figure have been contrast-optimized to facilitate the visualization of the examples, but the actual values were used when processing the images for quantification.

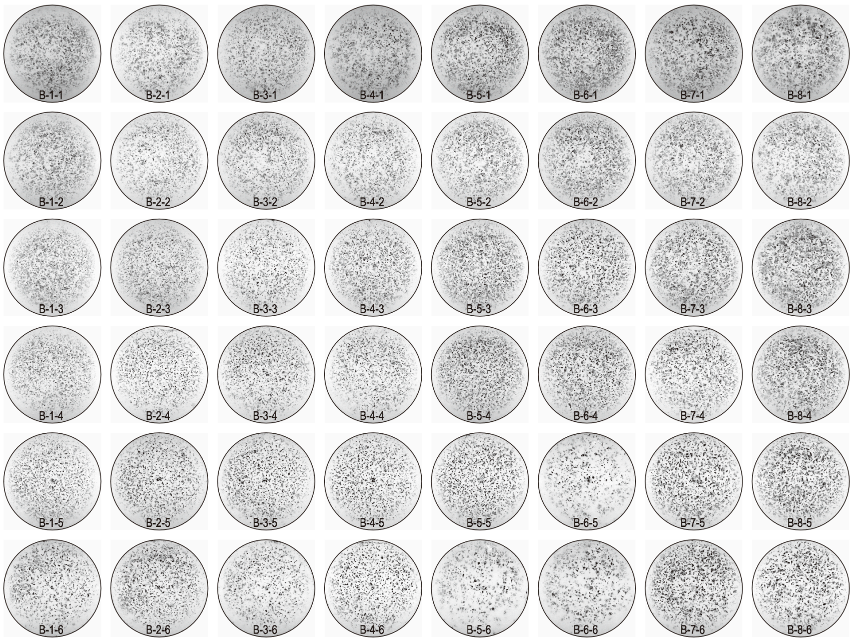

**Fig. S15: Acquired pigmented images of the baseline experiment**
Images acquired on Day 34 of the baseline experiment; images of the bottom of the well with cultured cells, cropped to the size of the well. These 8-bit images were adjusted to a minimum and maximum contrast value of 100 and 150, respectively. IDs on the bottom indicates 'B (baseline) - Plate No. - Well No.'.

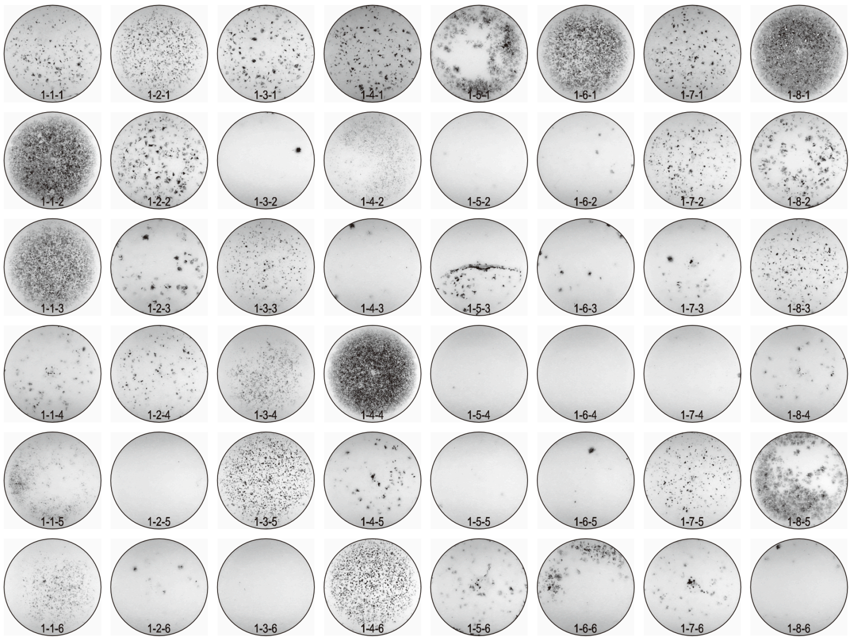

**Fig. S16: Acquired pigmented images of the round 1 experiment**
Images acquired on Day 34 of the round 1 experiment; images of the bottom of the well with cultured cells, cropped to the size of the well. These 8-bit images were adjusted to a minimum and maximum contrast value of 100 and 150, respectively. ID labeling on the bottom indicates '1 (round 1) - Plate No. - Well No.'.

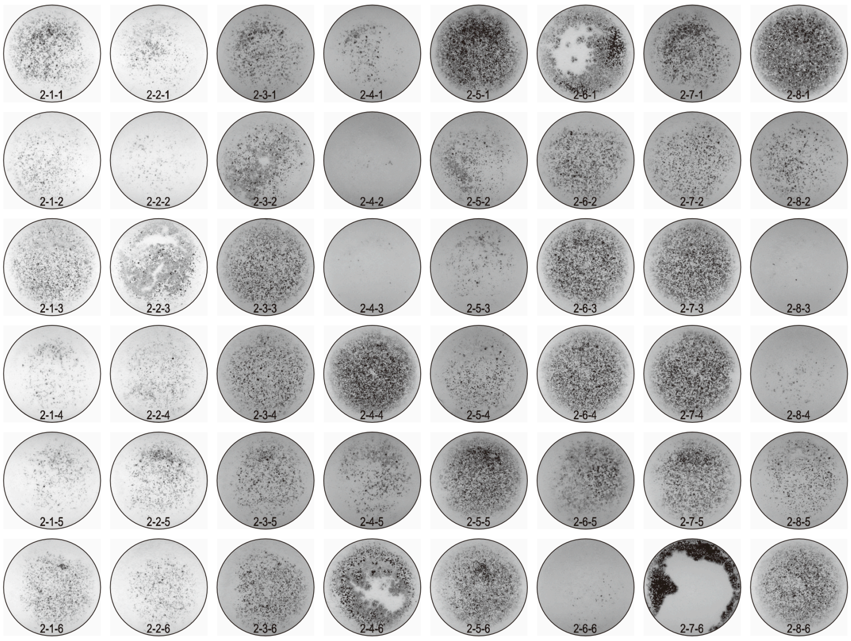


**Fig. S17: Acquired pigmented images of the round 2 experiment**
Images acquired on Day 34 of the round 2 experiment; images of the bottom of the well with cultured cells, cropped to the size of the well. These 8-bit images were adjusted to a minimum and maximum contrast value of 100 and 150, respectively. ID labeling on the bottom indicates '2 (round 2) - Plate No. - Well No.'.

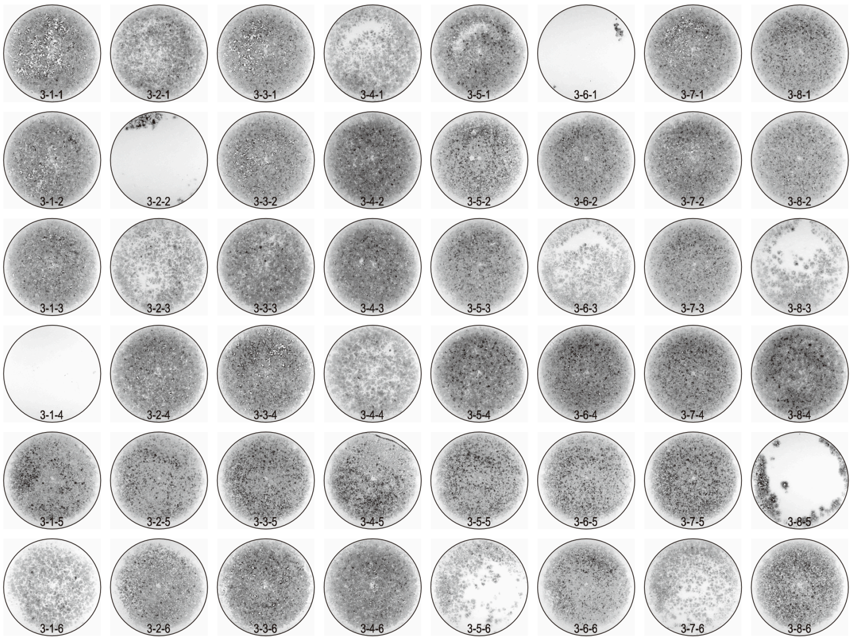


**Fig. S18: Acquired pigmented images of the round 3 experiment**
Images acquired on Day 34 of the round 3 experiment: images of the bottom of the well with cultured cells, cropped to the size of the well. These 8-bit images were adjusted to a minimum and maximum contrast value of 100 and 150, respectively. ID labeling on the bottom indicates '3 (round 3) - Plate No. - Well No.'.

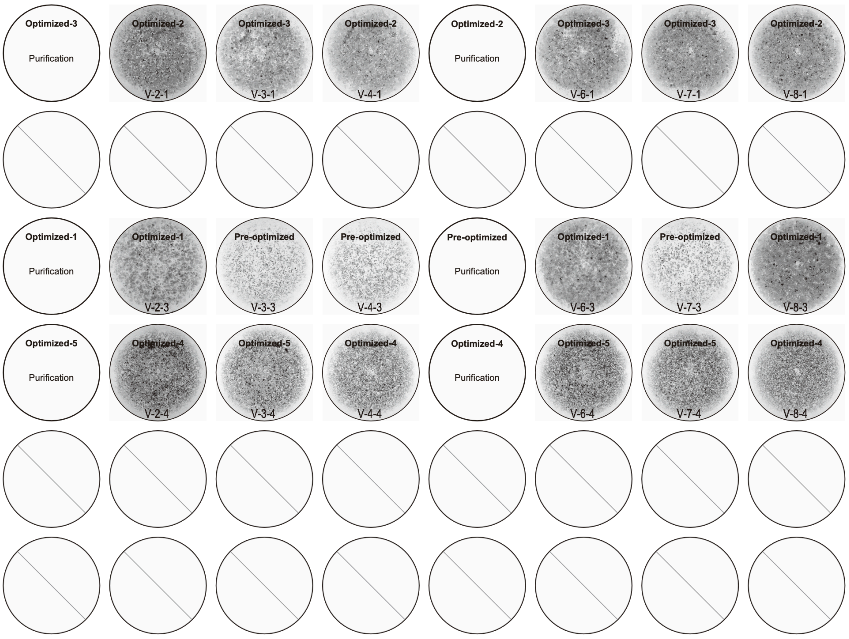


**Fig. S19: Acquired pigmented images of the validation experiment**

Images acquired on Day 34 of the validation experiment: images of the bottom of the well with cultured cells, cropped to the size of the well. These 8-bit images were adjusted to a minimum and maximum contrast value of 80 and 125, respectively. Sample names on the top correspond to **Fig. 5A**. ID labeling on the bottom indicates 'V (validation) - Plate No. - Well No.'. Wells 2, 5, and 6 were not subjected to the validation experiments. Plates 1 and 5 were used for cell biological analysis and were not evaluated using images.

*5.2. Score data visualization by PCP*

In this study, in order to optimize the culture conditions, a parallel experiment of 48 samples per 1 round was carried out for three rounds. **Fig. S20** is a visualization of the conditions and results of the series of experiments using a parallel coordinate plot (PCP). PCP is a common tool for visualizing and analyzing high-dimensional data. We can see that the experimental conditions tended to converge and the score tended to improve with each round.

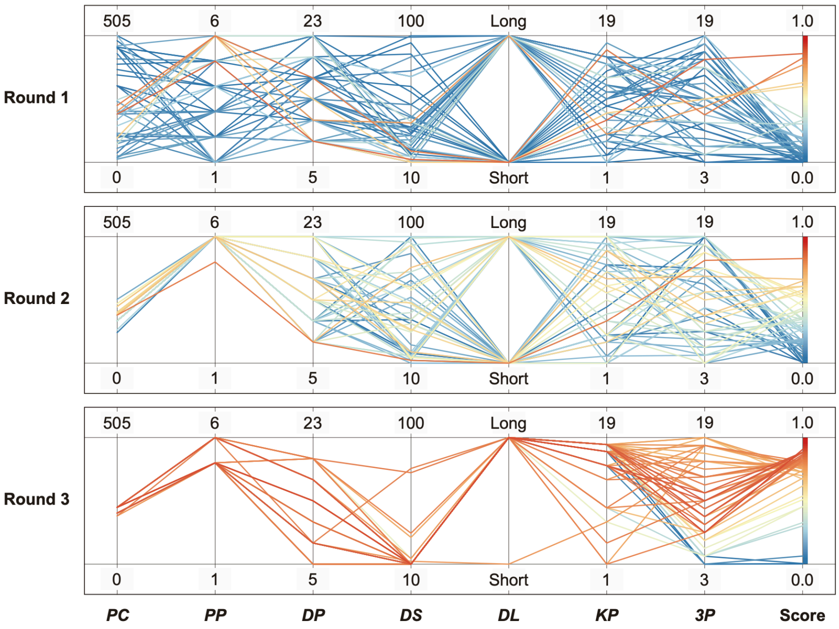


**Fig. S20: Parallel coordinate plot (PCP) of the robotic search experimental results**

The experiment results were visualized in rounds 1 to 3 using a parallel coordinate plot (PCP). The experimental results in 8-dimensional space (7-dimensional parameters + pigmented scores) are represented as a colored line with vertices on the parallel axes; the position of the vertex on the i-th axis corresponds to the i-th coordinate of the parameter. The color of the line represents the pigmented score: blue lines represent lower pigmented scores, and the closer to red, the higher the pigmented score.

### 6. Experimental Operations

*6.1. Daily robot experimental operation*

Daily robot experimental operations were conducted according to a daily check sheet to prevent human error and personalization of the operations. First, the temperature and $CO_2$ concentration displayed on the $CO_2$ incubator were checked and confirmed to be normal. Then, the proper working of the room dehumidifiers was checked. Next, the reagents to be used on that day were prepared and placed in the appropriate locations in the LabDroid booth. A LabDroid job was then executed to open and close the outer door of the $CO_2$ incubator, ensuring that the cell plate was in the proper location and position in the incubator. Labware, such as tips for micropipettes, was supplied if necessary, and the conditions of the reagents, labware, and cell plates were checked. Finally, an experimental job was executed. After execution, the reagents and labware were replenished, and a second experimental job was executed. After the execution of the final experimental job for the day, a tube connected to the aspirator was washed by the LabDroid with 70% ethanol. Then, the aspirator waste fluid tank was removed to dispose of the waste fluid, and the waste fluid tank was reinstalled. Then, a 70% ethanol aspiration job was executed to confirm that the waste tank had been properly reinstalled. Finally, cleaning of the dustbin in the LabDroid booth and disposal of the dehumidifier waste liquid were performed.

*6.2. 24/7 monitoring and recording*

The robot operation was monitored using two wide-angle cameras (24×7 monitoring), two magnifying cameras (recording only when the robot was running), and a live streaming camera (**Fig. S21**). Representative 24×7 monitoring camera logs are shown in **Movies S2–6**. Because the video footage of the LabDroid booth is continuously recorded, the behavior can be analyzed regardless of the time when an error occurs. The remote accessibility of the control PC allows the robot to be controlled without being in the laboratory, unless physical modifications are required.

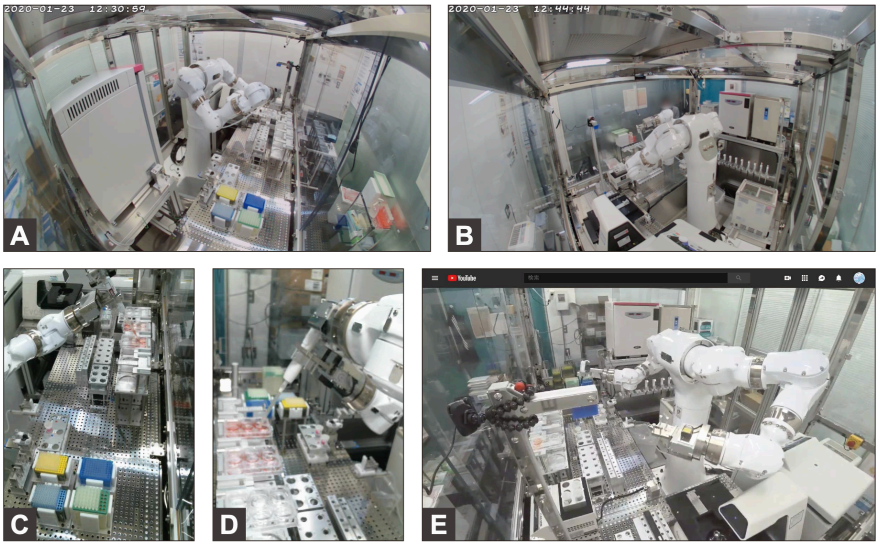


**Fig. S21: Video monitoring**

Example pictures of the monitoring cameras.

(**A–B**) Wide-angle cameras for 24×7 monitoring. Front camera (**A**) and back camera (**B**).

(**C–D**) Magnifying cameras that record only when the robot is running. Right camera (**C**) and left camera (**D**).

(**E**) Live streaming camera (available to research contributors only, no recordings).

*6.3. Execution logs and errors*

Robot experiments were conducted for 185 days, with a total robot operating time of 995 h, 39 min, and 21 s (**Table S5**). Each round consisted of 73 jobs, and five rounds were performed, including baseline and validation. The total number of jobs executed by the LabDroid was 365, of which 343 were successful on the first try; 22 required human intervention at least once. The job success rate was 93.973%. One job consisted of multiple commands. The total number of commands that the LabDroid was ordered to execute was 75039, of which 75011 were successful on the first try, 25 required human intervention at least once, and 3 were aborted. The command success rate was 99.963%. The reasons for the errors included micropipette tip loading error, 4 commands; micropipette tip ejection error, 7 commands; microscope and its control PC-derived errors, 13 commands (including 3 aborted commands); defective labware, 1 command; and human error, 3 commands. The errors occurred on the following dates: 2019 Feb (baseline), 11 commands; 2019 Apr (round 1), 11 (including three aborted commands); 2019 Jul (round 2), 4; 2020 Jan (round 3), 0; and 2020 Mar (validation), 2 (**Fig. S22**). The number of motions requiring the use of micropipettes that the LabDroid was ordered to execute was 39421; the rate of failure of either tip loading or ejection was 0.0279%.

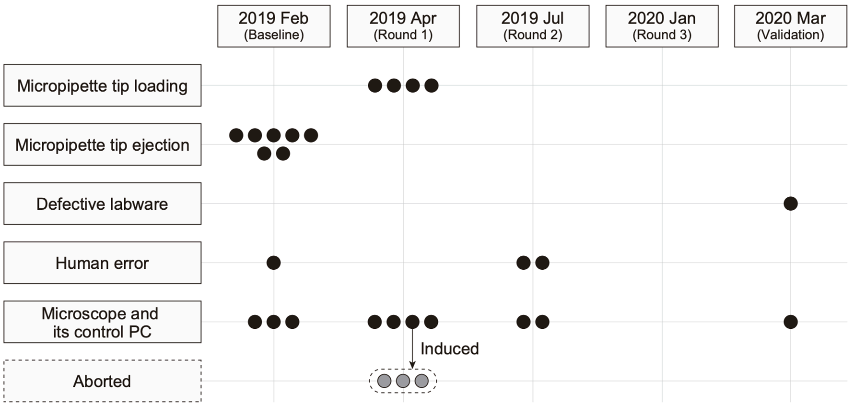


### Fig. S22: Errors in the robotic operations

Causes (vertical axis) and error timing (horizontal axis) of the errors that occurred during the robotic experiments. Each circle represents one error. The total number of commands the LabDroid was ordered to execute was 75039, of which 75011 were successful on the first try, 25 required human intervention at least once, and three were aborted. The three commands aborted in round 1 were caused by errors from the microscope and its control PC.

### 7. Supplementary Table Legends

#### Table S1: Pipetting volume and pipette combination

Related to **Fig. 2**. Given the limitations of the LabDroid, setting the micropipette to an arbitrary volume was difficult. Therefore, we pseudo-implemented a fine volume setting for the transfer of 0–1000 μL by combining nine micropipettes with pre-set volumes (3000, 1000, 450, 300, 200, 80, 30, 10, and 5 μL). The number of combinations was limited to three or fewer. The numbers in the table indicate the micropipettes and the number of times they had to be used to achieve the desired volumes. For example, 260 μL indicates that the 200 and 30 μL micropipettes had to be used once and twice, respectively.

#### Table S2: Executed parameters and scores of the baseline experiment

Related to **Fig. 2E**. Raw values of the parameter candidates and pigmentation scores in the baseline experiment. *KSR concentration was lowered in a systematic fashion, unlike the *KP* parameter. Specific values: DDays 1–3, 20% KSR; DDays 4–7, 15% KSR; from DDay 8, 10% KSR.

#### Table S3: Executed parameters and scores of the optimization experiments

Related to **Fig. 4**. Raw values of the parameter candidates and pigmentation scores in the experiments from rounds 1 to 3.

#### Table S4: Executed parameters and scores of the validation experiment

Related to **Fig. 4**. Raw values of the parameter candidates and pigmentation scores in the validation experiments. The sample names on the top correspond to **Fig. 5A**. Well numbers 2, 5, and 6 were not subjected to the validation experiments. *Plate numbers 1 and 5 were used for cell biological analysis and were not validated using images.

#### Table S5: Robot log

List of job names, start times, end times, time required, number of commands, and errors (if any) for all experiments performed by the LabDroid in this study.

#### Table S6: ELISA scores

Related to **Fig. 5D, E**. Raw values of ELISA scores from the validation experiments.

### 8. Supplementary Movie Legends

#### Movie S1: Representative LabDroid movements

Example of LabDroid movements extracted from actual cell culture operation: handling of a 50-mL tube, aspirator, micropipette, microscope, and $CO_2$ incubator. The speed factor was 100%.

#### Movie S2: Seeding operation

Video recording an example of a seeding operation (round 3, DDay -7). The speed factor was 6000%.

#### Movie S3: Preconditioning operation

Video recording an example of a preconditioning operation (round 3, DDay -6, 1st run). The speed factor was 6000%.

#### Movie S4: Passaging operation

Video recording an example of a passage operation (round 3, DDay 0, 1st run). The speed factor was 6000%.

#### Movie S5: RPE differentiation operation

Video recording an example of an RPE differentiation operation (round 3, DDay 10, 1st run). The speed factor was 6000%.

#### Movie S6: RPE maintenance operation

Video recording an example of an RPE maintenance operation (round 3, DDay 32, 1st run). The speed factor was 6000%.

Movies S1 to S6 are available at:
https://www.dropbox.com/sh/1wy9pvrqml7a3ur/AABKKxfWXNZlBFqxNwvll_Qia?dl=0

Table S1

| Pre-set Volume | Pipette combinations | | | | | | | | |
|---|---|---|---|---|---|---|---|---|---|
| | 3000 µL | 1000 µL | 450 µL | 300 µL | 200 µL | 80 µL | 30 µL | 10 µL | 5 µL |
| 0 | 0 | 0 | 0 | 0 | 0 | 0 | 0 | 0 | 0 |
| 5 | 0 | 0 | 0 | 0 | 0 | 0 | 0 | 0 | 1 |
| 10 | 0 | 0 | 0 | 0 | 0 | 0 | 0 | 1 | 0 |
| 15 | 0 | 0 | 0 | 0 | 0 | 0 | 0 | 1 | 1 |
| 20 | 0 | 0 | 0 | 0 | 0 | 0 | 0 | 2 | 0 |
| 25 | 0 | 0 | 0 | 0 | 0 | 0 | 0 | 2 | 1 |
| 30 | 0 | 0 | 0 | 0 | 0 | 0 | 1 | 0 | 0 |
| 35 | 0 | 0 | 0 | 0 | 0 | 0 | 1 | 0 | 1 |
| 40 | 0 | 0 | 0 | 0 | 0 | 0 | 1 | 1 | 0 |
| 45 | 0 | 0 | 0 | 0 | 0 | 0 | 1 | 1 | 1 |
| 50 | 0 | 0 | 0 | 0 | 0 | 0 | 1 | 2 | 0 |
| 60 | 0 | 0 | 0 | 0 | 0 | 0 | 2 | 0 | 0 |
| 65 | 0 | 0 | 0 | 0 | 0 | 0 | 2 | 0 | 1 |
| 70 | 0 | 0 | 0 | 0 | 0 | 0 | 2 | 1 | 0 |
| 80 | 0 | 0 | 0 | 0 | 0 | 1 | 0 | 0 | 0 |
| 85 | 0 | 0 | 0 | 0 | 0 | 1 | 0 | 0 | 1 |
| 90 | 0 | 0 | 0 | 0 | 0 | 1 | 0 | 1 | 0 |
| 95 | 0 | 0 | 0 | 0 | 0 | 1 | 0 | 1 | 1 |
| 100 | 0 | 0 | 0 | 0 | 0 | 1 | 0 | 2 | 0 |
| 110 | 0 | 0 | 0 | 0 | 0 | 1 | 1 | 0 | 0 |
| 115 | 0 | 0 | 0 | 0 | 0 | 1 | 1 | 0 | 1 |
| 120 | 0 | 0 | 0 | 0 | 0 | 1 | 1 | 1 | 0 |
| 140 | 0 | 0 | 0 | 0 | 0 | 1 | 2 | 0 | 0 |
| 160 | 0 | 0 | 0 | 0 | 0 | 2 | 0 | 0 | 0 |
| 165 | 0 | 0 | 0 | 0 | 0 | 2 | 0 | 0 | 1 |
| 170 | 0 | 0 | 0 | 0 | 0 | 2 | 0 | 1 | 0 |
| 190 | 0 | 0 | 0 | 0 | 0 | 2 | 1 | 0 | 0 |
| 200 | 0 | 0 | 0 | 0 | 1 | 0 | 0 | 0 | 0 |
| 205 | 0 | 0 | 0 | 0 | 1 | 0 | 0 | 0 | 1 |
| 210 | 0 | 0 | 0 | 0 | 1 | 0 | 0 | 1 | 0 |
| 215 | 0 | 0 | 0 | 0 | 1 | 0 | 0 | 1 | 1 |
| 220 | 0 | 0 | 0 | 0 | 1 | 0 | 0 | 2 | 0 |
| 230 | 0 | 0 | 0 | 0 | 1 | 0 | 1 | 0 | 0 |
| 235 | 0 | 0 | 0 | 0 | 1 | 0 | 1 | 0 | 1 |
| 240 | 0 | 0 | 0 | 0 | 1 | 0 | 1 | 1 | 0 |
| 260 | 0 | 0 | 0 | 0 | 1 | 0 | 2 | 0 | 0 |
| 280 | 0 | 0 | 0 | 0 | 1 | 1 | 0 | 0 | 0 |
| 285 | 0 | 0 | 0 | 0 | 1 | 1 | 0 | 0 | 1 |

| | | | | | | | | | |
|---|---|---|---|---|---|---|---|---|---|
| 290 | 0 | 0 | 0 | 0 | 1 | 1 | 0 | 1 | 0 |
| 300 | 0 | 0 | 0 | 1 | 0 | 0 | 0 | 0 | 0 |
| 305 | 0 | 0 | 0 | 1 | 0 | 0 | 0 | 0 | 1 |
| 310 | 0 | 0 | 0 | 1 | 0 | 0 | 0 | 1 | 0 |
| 315 | 0 | 0 | 0 | 1 | 0 | 0 | 0 | 1 | 1 |
| 320 | 0 | 0 | 0 | 1 | 0 | 0 | 0 | 2 | 0 |
| 330 | 0 | 0 | 0 | 1 | 0 | 0 | 1 | 0 | 0 |
| 335 | 0 | 0 | 0 | 1 | 0 | 0 | 1 | 0 | 1 |
| 340 | 0 | 0 | 0 | 1 | 0 | 0 | 1 | 1 | 0 |
| 360 | 0 | 0 | 0 | 1 | 0 | 0 | 2 | 0 | 0 |
| 380 | 0 | 0 | 0 | 1 | 0 | 1 | 0 | 0 | 0 |
| 385 | 0 | 0 | 0 | 1 | 0 | 1 | 0 | 0 | 1 |
| 390 | 0 | 0 | 0 | 1 | 0 | 1 | 0 | 1 | 0 |
| 400 | 0 | 0 | 0 | 0 | 2 | 0 | 0 | 0 | 0 |
| 405 | 0 | 0 | 0 | 0 | 2 | 0 | 0 | 0 | 1 |
| 410 | 0 | 0 | 0 | 1 | 0 | 1 | 1 | 0 | 0 |
| 430 | 0 | 0 | 0 | 0 | 2 | 0 | 1 | 0 | 0 |
| 450 | 0 | 0 | 1 | 0 | 0 | 0 | 0 | 0 | 0 |
| 455 | 0 | 0 | 1 | 0 | 0 | 0 | 0 | 0 | 1 |
| 460 | 0 | 0 | 1 | 0 | 0 | 0 | 0 | 1 | 0 |
| 465 | 0 | 0 | 1 | 0 | 0 | 0 | 0 | 1 | 1 |
| 470 | 0 | 0 | 1 | 0 | 0 | 0 | 0 | 2 | 0 |
| 480 | 0 | 0 | 1 | 0 | 0 | 0 | 1 | 0 | 0 |
| 485 | 0 | 0 | 1 | 0 | 0 | 0 | 1 | 0 | 1 |
| 490 | 0 | 0 | 1 | 0 | 0 | 0 | 1 | 1 | 0 |
| 500 | 0 | 0 | 0 | 1 | 1 | 0 | 0 | 0 | 0 |
| 505 | 0 | 0 | 0 | 1 | 1 | 0 | 0 | 0 | 1 |
| 510 | 0 | 0 | 1 | 0 | 0 | 0 | 2 | 0 | 0 |
| 530 | 0 | 0 | 1 | 0 | 0 | 1 | 0 | 0 | 0 |
| 535 | 0 | 0 | 1 | 0 | 0 | 1 | 0 | 0 | 1 |
| 540 | 0 | 0 | 1 | 0 | 0 | 1 | 0 | 1 | 0 |
| 560 | 0 | 0 | 1 | 0 | 0 | 1 | 1 | 0 | 0 |
| 580 | 0 | 0 | 0 | 1 | 1 | 1 | 0 | 0 | 0 |
| 600 | 0 | 0 | 0 | 2 | 0 | 0 | 0 | 0 | 0 |
| 605 | 0 | 0 | 0 | 2 | 0 | 0 | 0 | 0 | 1 |
| 610 | 0 | 0 | 1 | 0 | 0 | 2 | 0 | 0 | 0 |
| 630 | 0 | 0 | 0 | 2 | 0 | 0 | 1 | 0 | 0 |
| 650 | 0 | 0 | 1 | 0 | 1 | 0 | 0 | 0 | 0 |
| 655 | 0 | 0 | 1 | 0 | 1 | 0 | 0 | 0 | 1 |
| 660 | 0 | 0 | 1 | 0 | 1 | 0 | 0 | 1 | 0 |
| 680 | 0 | 0 | 1 | 0 | 1 | 0 | 1 | 0 | 0 |

| | | | | | | | | | |
|---|---|---|---|---|---|---|---|---|---|
| 700 | 0 | 0 | 0 | 1 | 2 | 0 | 0 | 0 | 0 |
| 730 | 0 | 0 | 1 | 0 | 1 | 1 | 0 | 0 | 0 |
| 750 | 0 | 0 | 1 | 1 | 0 | 0 | 0 | 0 | 0 |
| 755 | 0 | 0 | 1 | 1 | 0 | 0 | 0 | 0 | 1 |
| 760 | 0 | 0 | 1 | 1 | 0 | 0 | 0 | 1 | 0 |
| 780 | 0 | 0 | 1 | 1 | 0 | 0 | 1 | 0 | 0 |
| 800 | 0 | 0 | 0 | 2 | 1 | 0 | 0 | 0 | 0 |
| 830 | 0 | 0 | 1 | 1 | 0 | 1 | 0 | 0 | 0 |
| 850 | 0 | 0 | 1 | 0 | 2 | 0 | 0 | 0 | 0 |
| 900 | 0 | 0 | 2 | 0 | 0 | 0 | 0 | 0 | 0 |
| 905 | 0 | 0 | 2 | 0 | 0 | 0 | 0 | 0 | 1 |
| 910 | 0 | 0 | 2 | 0 | 0 | 0 | 0 | 1 | 0 |
| 930 | 0 | 0 | 2 | 0 | 0 | 0 | 1 | 0 | 0 |
| 950 | 0 | 0 | 1 | 1 | 1 | 0 | 0 | 0 | 0 |
| 980 | 0 | 0 | 2 | 0 | 0 | 1 | 0 | 0 | 0 |
| 1000 | 0 | 1 | 0 | 0 | 0 | 0 | 0 | 0 | 0 |

**Table S2**

| | Plate No. | Well No. | Parameters | | | | | | | Score |
|---|---|---|---|---|---|---|---|---|---|---|
| | | | *PC* | *PP* | *DP* | *DS* | *DL* | *KP* | *3P* | |
| Baseline | 1 | 1 | 100 | 6 | 8 | 10 | S | * | 11 | 0.46 |
| Baseline | 2 | 1 | 100 | 6 | 8 | 10 | S | * | 11 | 0.45 |
| Baseline | 3 | 1 | 100 | 6 | 8 | 10 | S | * | 11 | 0.41 |
| Baseline | 4 | 1 | 100 | 6 | 8 | 10 | S | * | 11 | 0.36 |
| Baseline | 5 | 1 | 100 | 6 | 8 | 10 | S | * | 11 | 0.46 |
| Baseline | 6 | 1 | 100 | 6 | 8 | 10 | S | * | 11 | 0.45 |
| Baseline | 7 | 1 | 100 | 6 | 8 | 10 | S | * | 11 | 0.46 |
| Baseline | 8 | 1 | 100 | 6 | 8 | 10 | S | * | 11 | 0.44 |
| Baseline | 1 | 2 | 100 | 6 | 11 | 10 | S | * | 11 | 0.44 |
| Baseline | 2 | 2 | 100 | 6 | 11 | 10 | S | * | 11 | 0.36 |
| Baseline | 3 | 2 | 100 | 6 | 11 | 10 | S | * | 11 | 0.39 |
| Baseline | 4 | 2 | 100 | 6 | 11 | 10 | S | * | 11 | 0.35 |
| Baseline | 5 | 2 | 100 | 6 | 11 | 10 | S | * | 11 | 0.44 |
| Baseline | 6 | 2 | 100 | 6 | 11 | 10 | S | * | 11 | 0.42 |
| Baseline | 7 | 2 | 100 | 6 | 11 | 10 | S | * | 11 | 0.41 |
| Baseline | 8 | 2 | 100 | 6 | 11 | 10 | S | * | 11 | 0.44 |
| Baseline | 1 | 3 | 100 | 6 | 14 | 10 | S | * | 11 | 0.52 |
| Baseline | 2 | 3 | 100 | 6 | 14 | 10 | S | * | 11 | 0.43 |
| Baseline | 3 | 3 | 100 | 6 | 14 | 10 | S | * | 11 | 0.34 |
| Baseline | 4 | 3 | 100 | 6 | 14 | 10 | S | * | 11 | 0.44 |
| Baseline | 5 | 3 | 100 | 6 | 14 | 10 | S | * | 11 | 0.48 |
| Baseline | 6 | 3 | 100 | 6 | 14 | 10 | S | * | 11 | 0.36 |
| Baseline | 7 | 3 | 100 | 6 | 14 | 10 | S | * | 11 | 0.49 |
| Baseline | 8 | 3 | 100 | 6 | 14 | 10 | S | * | 11 | 0.44 |
| Baseline | 1 | 4 | 100 | 6 | 17 | 10 | S | * | 11 | 0.44 |
| Baseline | 2 | 4 | 100 | 6 | 17 | 10 | S | * | 11 | 0.36 |
| Baseline | 3 | 4 | 100 | 6 | 17 | 10 | S | * | 11 | 0.28 |
| Baseline | 4 | 4 | 100 | 6 | 17 | 10 | S | * | 11 | 0.30 |
| Baseline | 5 | 4 | 100 | 6 | 17 | 10 | S | * | 11 | 0.41 |
| Baseline | 6 | 4 | 100 | 6 | 17 | 10 | S | * | 11 | 0.41 |
| Baseline | 7 | 4 | 100 | 6 | 17 | 10 | S | * | 11 | 0.44 |
| Baseline | 8 | 4 | 100 | 6 | 17 | 10 | S | * | 11 | 0.52 |
| Baseline | 1 | 5 | 100 | 6 | 20 | 10 | S | * | 11 | 0.43 |
| Baseline | 2 | 5 | 100 | 6 | 20 | 10 | S | * | 11 | 0.35 |
| Baseline | 3 | 5 | 100 | 6 | 20 | 10 | S | * | 11 | 0.32 |
| Baseline | 4 | 5 | 100 | 6 | 20 | 10 | S | * | 11 | 0.31 |
| Baseline | 5 | 5 | 100 | 6 | 20 | 10 | S | * | 11 | 0.34 |
| Baseline | 6 | 5 | 100 | 6 | 20 | 10 | S | * | 11 | 0.25 |
| Baseline | 7 | 5 | 100 | 6 | 20 | 10 | S | * | 11 | 0.38 |
| Baseline | 8 | 5 | 100 | 6 | 20 | 10 | S | * | 11 | 0.46 |
| Baseline | 1 | 6 | 100 | 6 | 23 | 10 | S | * | 11 | 0.32 |

| | | | | | | | | | | |
|---|---|---|---|---|---|---|---|---|---|---|
| Baseline | 2 | 6 | 100 | 6 | 23 | 10 | S | * | 11 | 0.38 |
| Baseline | 3 | 6 | 100 | 6 | 23 | 10 | S | * | 11 | 0.30 |
| Baseline | 4 | 6 | 100 | 6 | 23 | 10 | S | * | 11 | 0.28 |
| Baseline | 5 | 6 | 100 | 6 | 23 | 10 | S | * | 11 | 0.23 |
| Baseline | 6 | 6 | 100 | 6 | 23 | 10 | S | * | 11 | 0.34 |
| Baseline | 7 | 6 | 100 | 6 | 23 | 10 | S | * | 11 | 0.38 |
| Baseline | 8 | 6 | 100 | 6 | 23 | 10 | S | * | 11 | 0.39 |

**Table S3**

| Round No. | Plate No. | Well No. | Parameters | | | | | | | Score |
|---|---|---|---|---|---|---|---|---|---|---|
| | | | *PC* | *PP* | *DP* | *DS* | *DL* | *KP* | *3P* | |
| 1 | 8 | 1 | 192.3 | 5 | 8 | 12 | S | 7 | 16 | 0.86 |
| 1 | 4 | 4 | 226.7 | 6 | 17 | 18 | S | 17 | 9 | 0.82 |
| 1 | 1 | 2 | 203.8 | 6 | 11 | 38 | L | 5 | 10 | 0.77 |
| 1 | 1 | 3 | 98 | 6 | 14 | 10 | S | 8 | 11 | 0.63 |
| 1 | 6 | 1 | 98 | 6 | 8 | 10 | S | 8 | 11 | 0.60 |
| 1 | 4 | 6 | 73.9 | 6 | 23 | 14 | L | 13 | 15 | 0.35 |
| 1 | 8 | 5 | 192.3 | 5 | 20 | 39 | S | 6 | 4 | 0.33 |
| 1 | 5 | 1 | 249.4 | 5 | 8 | 23 | L | 8 | 8 | 0.30 |
| 1 | 2 | 1 | 145.6 | 4 | 8 | 14 | S | 15 | 15 | 0.24 |
| 1 | 3 | 5 | 192.3 | 4 | 20 | 71 | L | 1 | 19 | 0.24 |
| 1 | 2 | 2 | 505.6 | 1 | 11 | 11 | S | 17 | 9 | 0.20 |
| 1 | 3 | 4 | 169.1 | 6 | 17 | 20 | S | 15 | 11 | 0.19 |
| 1 | 1 | 6 | 192.3 | 6 | 23 | 24 | L | 1 | 11 | 0.18 |
| 1 | 8 | 2 | 243.8 | 1 | 11 | 14 | S | 4 | 5 | 0.17 |
| 1 | 4 | 2 | 464.9 | 6 | 11 | 34 | L | 13 | 18 | 0.14 |
| 1 | 3 | 1 | 203.8 | 1 | 8 | 26 | L | 12 | 5 | 0.12 |
| 1 | 4 | 1 | 86 | 2 | 8 | 10 | L | 7 | 8 | 0.12 |
| 1 | 1 | 1 | 24.9 | 2 | 8 | 42 | L | 15 | 5 | 0.11 |
| 1 | 7 | 2 | 12.5 | 2 | 11 | 38 | S | 18 | 12 | 0.11 |
| 1 | 8 | 3 | 73.9 | 3 | 14 | 85 | L | 15 | 10 | 0.11 |
| 1 | 7 | 1 | 116 | 2 | 8 | 29 | S | 11 | 13 | 0.10 |
| 1 | 3 | 3 | 18.7 | 4 | 14 | 33 | L | 10 | 16 | 0.09 |
| 1 | 7 | 5 | 55.6 | 3 | 20 | 43 | S | 5 | 17 | 0.08 |
| 1 | 6 | 6 | 412.8 | 5 | 23 | 14 | S | 4 | 17 | 0.06 |
| 1 | 2 | 4 | 18.7 | 3 | 17 | 42 | S | 12 | 10 | 0.06 |
| 1 | 5 | 3 | 110 | 1 | 14 | 51 | S | 5 | 6 | 0.06 |
| 1 | 2 | 3 | 203.8 | 3 | 14 | 22 | L | 8 | 4 | 0.06 |
| 1 | 4 | 5 | 192.3 | 3 | 20 | 71 | L | 16 | 11 | 0.05 |
| 1 | 5 | 6 | 37.2 | 5 | 23 | 15 | L | 16 | 3 | 0.04 |
| 1 | 7 | 6 | 98 | 3 | 23 | 99 | L | 12 | 11 | 0.04 |
| 1 | 1 | 4 | 433.8 | 1 | 17 | 75 | L | 11 | 11 | 0.04 |
| 1 | 7 | 3 | 104 | 2 | 14 | 30 | L | 15 | 7 | 0.02 |
| 1 | 8 | 4 | 243.8 | 1 | 17 | 100 | L | 16 | 12 | 0.02 |
| 1 | 6 | 3 | 305.2 | 1 | 14 | 35 | L | 3 | 4 | 0.01 |
| 1 | 2 | 6 | 305.2 | 3 | 23 | 31 | L | 2 | 15 | 0.01 |
| 1 | 6 | 5 | 348.8 | 4 | 20 | 87 | S | 14 | 19 | 0.01 |
| 1 | 6 | 2 | 370.4 | 4 | 11 | 29 | S | 10 | 18 | 0.01 |
| 1 | 3 | 2 | 459.7 | 1 | 11 | 19 | S | 4 | 3 | 0.01 |
| 1 | 4 | 3 | 454.6 | 4 | 14 | 19 | L | 10 | 14 | 0.00 |
| 1 | 5 | 4 | 305.2 | 2 | 17 | 33 | S | 2 | 8 | 0.00 |
| 1 | 3 | 6 | 459.7 | 5 | 23 | 60 | S | 1 | 4 | 0.00 |

| | | | | | | | | | | |
|---|---|---|---|---|---|---|---|---|---|---|
| 1 | 5 | 2 | 305.2 | 3 | 11 | 35 | S | 10 | 17 | 0.00 |
| 1 | 5 | 5 | 459.7 | 2 | 20 | 14 | L | 16 | 16 | 0.00 |
| 1 | 7 | 4 | 505.6 | 4 | 17 | 95 | S | 13 | 17 | 0.00 |
| 1 | 8 | 6 | 391.7 | 5 | 23 | 16 | S | 9 | 14 | 0.00 |
| 1 | 2 | 5 | 485.3 | 3 | 20 | 51 | L | 9 | 7 | 0.00 |
| 1 | 6 | 4 | 459.7 | 5 | 17 | 64 | S | 5 | 3 | 0.00 |
| 2 | 8 | 1 | 192.3 | 5 | 8 | 12 | S | 7 | 16 | 0.83 |
| 2 | 5 | 1 | 192.3 | 6 | 8 | 78 | L | 13 | 14 | 0.66 |
| 2 | 4 | 4 | 226.7 | 6 | 17 | 18 | S | 17 | 9 | 0.65 |
| 2 | 6 | 3 | 198.1 | 6 | 14 | 53 | S | 4 | 12 | 0.61 |
| 2 | 6 | 4 | 203.8 | 6 | 17 | 34 | S | 17 | 11 | 0.61 |
| 2 | 7 | 4 | 198.1 | 6 | 17 | 33 | L | 12 | 8 | 0.58 |
| 2 | 7 | 3 | 198.1 | 6 | 14 | 94 | S | 14 | 9 | 0.55 |
| 2 | 5 | 5 | 203.8 | 6 | 20 | 64 | L | 11 | 17 | 0.55 |
| 2 | 5 | 6 | 238.1 | 6 | 23 | 13 | L | 5 | 17 | 0.53 |
| 2 | 8 | 6 | 226.7 | 6 | 23 | 14 | S | 12 | 15 | 0.52 |
| 2 | 4 | 6 | 226.7 | 6 | 23 | 62 | L | 19 | 3 | 0.50 |
| 2 | 7 | 5 | 203.8 | 6 | 20 | 52 | S | 1 | 18 | 0.45 |
| 2 | 6 | 1 | 192.3 | 6 | 8 | 38 | S | 10 | 5 | 0.44 |
| 2 | 2 | 3 | 192.3 | 6 | 14 | 41 | L | 5 | 3 | 0.42 |
| 2 | 3 | 3 | 192.3 | 6 | 14 | 32 | L | 19 | 7 | 0.41 |
| 2 | 1 | 3 | 192.3 | 6 | 14 | 13 | L | 11 | 6 | 0.36 |
| 2 | 6 | 2 | 169.1 | 6 | 11 | 46 | L | 17 | 18 | 0.35 |
| 2 | 3 | 4 | 203.8 | 6 | 17 | 75 | L | 7 | 6 | 0.35 |
| 2 | 1 | 1 | 169.1 | 6 | 8 | 15 | S | 8 | 12 | 0.34 |
| 2 | 7 | 1 | 169.1 | 6 | 8 | 99 | L | 19 | 19 | 0.34 |
| 2 | 3 | 6 | 203.8 | 6 | 23 | 96 | S | 7 | 7 | 0.32 |
| 2 | 3 | 2 | 169.1 | 6 | 11 | 94 | L | 16 | 4 | 0.31 |
| 2 | 1 | 6 | 203.8 | 6 | 23 | 100 | L | 19 | 14 | 0.28 |
| 2 | 6 | 5 | 133.8 | 6 | 20 | 54 | S | 18 | 3 | 0.27 |
| 2 | 2 | 5 | 192.3 | 6 | 20 | 18 | S | 5 | 15 | 0.26 |
| 2 | 3 | 1 | 169.1 | 6 | 8 | 15 | L | 18 | 19 | 0.26 |
| 2 | 7 | 2 | 192.3 | 6 | 11 | 66 | L | 1 | 10 | 0.26 |
| 2 | 3 | 5 | 192.3 | 6 | 20 | 18 | L | 14 | 15 | 0.24 |
| 2 | 5 | 4 | 198.1 | 6 | 17 | 49 | S | 2 | 4 | 0.19 |
| 2 | 8 | 5 | 192.3 | 6 | 20 | 16 | L | 5 | 4 | 0.19 |
| 2 | 5 | 2 | 169.1 | 6 | 11 | 10 | S | 4 | 5 | 0.18 |
| 2 | 8 | 2 | 169.1 | 6 | 11 | 12 | L | 18 | 12 | 0.18 |
| 2 | 2 | 4 | 192.3 | 6 | 17 | 15 | S | 14 | 5 | 0.17 |
| 2 | 2 | 6 | 203.8 | 6 | 23 | 65 | S | 19 | 18 | 0.17 |
| 2 | 4 | 5 | 192.3 | 6 | 20 | 13 | L | 15 | 8 | 0.16 |
| 2 | 2 | 1 | 169.1 | 6 | 8 | 57 | S | 5 | 6 | 0.16 |
| 2 | 1 | 2 | 169.1 | 6 | 11 | 74 | S | 19 | 7 | 0.15 |
| 2 | 7 | 6 | 255 | 6 | 23 | 23 | S | 18 | 3 | 0.15 |

| | | | | | | | | | | |
|---|---|---|---|---|---|---|---|---|---|---|
| 2 | 1 | 5 | 192.3 | 6 | 20 | 41 | L | 2 | 14 | 0.12 |
| 2 | 1 | 4 | 192.3 | 6 | 17 | 88 | S | 5 | 18 | 0.11 |
| 2 | 4 | 1 | 169.1 | 6 | 8 | 69 | S | 15 | 13 | 0.09 |
| 2 | 5 | 3 | 169.1 | 6 | 14 | 15 | L | 4 | 9 | 0.07 |
| 2 | 2 | 2 | 169.1 | 6 | 11 | 100 | S | 19 | 19 | 0.04 |
| 2 | 8 | 4 | 122 | 6 | 17 | 38 | S | 9 | 19 | 0.03 |
| 2 | 6 | 6 | 169.1 | 6 | 23 | 12 | S | 5 | 7 | 0.01 |
| 2 | 4 | 2 | 169.1 | 6 | 11 | 41 | S | 1 | 19 | 0.00 |
| 2 | 4 | 3 | 169.1 | 6 | 14 | 17 | S | 15 | 15 | 0.00 |
| 2 | 8 | 3 | 169.1 | 6 | 14 | 17 | S | 1 | 10 | 0.00 |
| 3 | 8 | 4 | 226.7 | 5 | 14 | 10 | L | 17 | 7 | 0.91 |
| 3 | 7 | 2 | 203.8 | 5 | 8 | 10 | L | 18 | 11 | 0.89 |
| 3 | 4 | 2 | 203.8 | 5 | 8 | 10 | L | 18 | 9 | 0.88 |
| 3 | 3 | 5 | 226.7 | 6 | 17 | 10 | L | 15 | 12 | 0.88 |
| 3 | 7 | 5 | 226.7 | 6 | 17 | 10 | L | 15 | 11 | 0.88 |
| 3 | 5 | 4 | 226.7 | 5 | 14 | 10 | L | 17 | 9 | 0.87 |
| 3 | 1 | 5 | 226.7 | 6 | 17 | 10 | L | 15 | 8 | 0.86 |
| 3 | 4 | 3 | 203.8 | 5 | 11 | 10 | L | 18 | 10 | 0.84 |
| 3 | 5 | 1 | 203.8 | 5 | 5 | 10 | L | 18 | 10 | 0.83 |
| 3 | 6 | 2 | 203.8 | 5 | 8 | 10 | L | 18 | 14 | 0.83 |
| 3 | 8 | 6 | 226.7 | 6 | 20 | 75 | L | 4 | 11 | 0.83 |
| 3 | 7 | 1 | 203.8 | 5 | 5 | 10 | L | 18 | 13 | 0.83 |
| 3 | 4 | 5 | 226.7 | 6 | 17 | 10 | L | 15 | 7 | 0.82 |
| 3 | 3 | 3 | 203.8 | 5 | 11 | 10 | L | 18 | 8 | 0.82 |
| 3 | 5 | 2 | 192.3 | 6 | 8 | 78 | L | 13 | 14 | 0.82 |
| 3 | 6 | 4 | 226.7 | 5 | 14 | 10 | L | 17 | 14 | 0.81 |
| 3 | 3 | 6 | 226.7 | 5 | 20 | 10 | L | 9 | 18 | 0.81 |
| 3 | 2 | 6 | 226.7 | 6 | 20 | 32 | L | 1 | 7 | 0.81 |
| 3 | 4 | 6 | 226.7 | 5 | 20 | 10 | L | 9 | 11 | 0.81 |
| 3 | 2 | 4 | 226.7 | 5 | 14 | 10 | L | 17 | 12 | 0.80 |
| 3 | 1 | 2 | 203.8 | 5 | 8 | 10 | L | 18 | 12 | 0.79 |
| 3 | 1 | 1 | 203.8 | 5 | 5 | 10 | L | 18 | 15 | 0.79 |
| 3 | 3 | 4 | 226.7 | 5 | 14 | 10 | L | 17 | 18 | 0.79 |
| 3 | 3 | 2 | 203.8 | 5 | 8 | 10 | L | 18 | 18 | 0.79 |
| 3 | 1 | 3 | 203.8 | 5 | 11 | 10 | L | 18 | 14 | 0.79 |
| 3 | 5 | 5 | 226.7 | 6 | 17 | 10 | L | 15 | 16 | 0.78 |
| 3 | 6 | 5 | 226.7 | 6 | 17 | 14 | L | 13 | 18 | 0.78 |
| 3 | 8 | 2 | 192.3 | 5 | 8 | 12 | S | 7 | 16 | 0.77 |
| 3 | 3 | 1 | 203.8 | 5 | 5 | 10 | L | 18 | 19 | 0.76 |
| 3 | 7 | 4 | 226.7 | 5 | 14 | 10 | L | 17 | 16 | 0.76 |
| 3 | 6 | 6 | 226.7 | 6 | 20 | 29 | L | 1 | 18 | 0.75 |
| 3 | 2 | 1 | 203.8 | 5 | 5 | 10 | L | 18 | 9 | 0.74 |
| 3 | 2 | 5 | 226.7 | 6 | 17 | 10 | L | 15 | 19 | 0.73 |
| 3 | 8 | 1 | 203.8 | 5 | 5 | 10 | L | 18 | 17 | 0.73 |

| | | | | | | | | | | |
|---|---|---|---|---|---|---|---|---|---|---|
| 3 | 5 | 3 | 203.8 | 5 | 11 | 10 | L | 18 | 16 | 0.70 |
| 3 | 1 | 6 | 226.7 | 5 | 20 | 10 | L | 9 | 7 | 0.68 |
| 3 | 7 | 3 | 203.8 | 5 | 11 | 10 | L | 18 | 19 | 0.67 |
| 3 | 4 | 4 | 226.7 | 5 | 14 | 10 | L | 17 | 6 | 0.62 |
| 3 | 2 | 3 | 203.8 | 5 | 11 | 10 | L | 18 | 6 | 0.54 |
| 3 | 4 | 1 | 203.8 | 5 | 5 | 10 | L | 18 | 6 | 0.51 |
| 3 | 6 | 3 | 203.8 | 5 | 11 | 10 | L | 18 | 5 | 0.45 |
| 3 | 7 | 6 | 226.7 | 5 | 20 | 14 | L | 7 | 4 | 0.45 |
| 3 | 8 | 3 | 203.8 | 5 | 11 | 10 | L | 18 | 4 | 0.33 |
| 3 | 5 | 6 | 226.7 | 5 | 20 | 10 | L | 9 | 4 | 0.30 |
| 3 | 8 | 5 | 226.7 | 6 | 17 | 10 | L | 15 | 3 | 0.06 |
| 3 | 2 | 2 | 203.8 | 5 | 8 | 10 | L | 18 | 4 | 0.01 |
| 3 | 6 | 1 | 203.8 | 5 | 5 | 10 | L | 18 | 3 | 0.00 |
| 3 | 1 | 4 | 226.7 | 5 | 14 | 10 | L | 17 | 3 | 0.00 |

**Table S4**

| | Plate No. | Well No. | Sample name | Parameters | | | | | | | Score |
|---|---|---|---|---|---|---|---|---|---|---|---|
| | | | | *PC* | *PP* | *DP* | *DS* | *DL* | *KP* | *3P* | |
| Validation | 1 | 1 | Optimized-3 | 203.8 | 5 | 8 | 10 | L | 18 | 9 | * |
| Validation | 1 | 2 | - | - | - | - | - | - | - | - | - |
| Validation | 1 | 3 | Optimized-1 | 226.7 | 5 | 14 | 10 | L | 17 | 7 | * |
| Validation | 1 | 4 | Optimized-5 | 226.7 | 6 | 17 | 10 | L | 15 | 11 | * |
| Validation | 1 | 5 | - | - | - | - | - | - | - | - | - |
| Validation | 1 | 6 | - | - | - | - | - | - | - | - | - |
| Validation | 2 | 1 | Optimized-2 | 203.8 | 5 | 8 | 10 | L | 18 | 11 | 0.78 |
| Validation | 2 | 2 | - | - | - | - | - | - | - | - | - |
| Validation | 2 | 3 | Optimized-1 | 226.7 | 5 | 14 | 10 | L | 17 | 7 | 0.61 |
| Validation | 2 | 4 | Optimized-4 | 226.7 | 6 | 17 | 10 | L | 15 | 12 | 0.82 |
| Validation | 2 | 5 | - | - | - | - | - | - | - | - | - |
| Validation | 2 | 6 | - | - | - | - | - | - | - | - | - |
| Validation | 3 | 1 | Optimized-3 | 203.8 | 5 | 8 | 10 | L | 18 | 9 | 0.72 |
| Validation | 3 | 2 | - | - | - | - | - | - | - | - | - |
| Validation | 3 | 3 | Pre-optimized | 98 | 6 | 14 | 10 | S | 8 | 11 | 0.41 |
| Validation | 3 | 4 | Optimized-5 | 226.7 | 6 | 17 | 10 | L | 15 | 11 | 0.82 |
| Validation | 3 | 5 | - | - | - | - | - | - | - | - | - |
| Validation | 3 | 6 | - | - | - | - | - | - | - | - | - |
| Validation | 4 | 1 | Optimized-2 | 203.8 | 5 | 8 | 10 | L | 18 | 11 | 0.68 |
| Validation | 4 | 2 | - | - | - | - | - | - | - | - | - |
| Validation | 4 | 3 | Pre-optimized | 98 | 6 | 14 | 10 | S | 8 | 11 | 0.48 |
| Validation | 4 | 4 | Optimized-4 | 226.7 | 6 | 17 | 10 | L | 15 | 12 | 0.77 |
| Validation | 4 | 5 | - | - | - | - | - | - | - | - | - |
| Validation | 4 | 6 | - | - | - | - | - | - | - | - | - |
| Validation | 5 | 1 | Optimized-2 | 203.8 | 5 | 8 | 10 | L | 18 | 11 | * |
| Validation | 5 | 2 | - | - | - | - | - | - | - | - | - |
| Validation | 5 | 3 | Pre-optimized | 98 | 6 | 14 | 10 | S | 8 | 11 | * |
| Validation | 5 | 4 | Optimized-4 | 226.7 | 6 | 17 | 10 | L | 15 | 12 | * |
| Validation | 5 | 5 | - | - | - | - | - | - | - | - | - |
| Validation | 5 | 6 | - | - | - | - | - | - | - | - | - |
| Validation | 6 | 1 | Optimized-3 | 203.8 | 5 | 8 | 10 | L | 18 | 9 | 0.75 |
| Validation | 6 | 2 | - | - | - | - | - | - | - | - | - |
| Validation | 6 | 3 | Optimized-1 | 226.7 | 5 | 14 | 10 | L | 17 | 7 | 0.70 |
| Validation | 6 | 4 | Optimized-5 | 226.7 | 6 | 17 | 10 | L | 15 | 11 | 0.84 |
| Validation | 6 | 5 | - | - | - | - | - | - | - | - | - |
| Validation | 6 | 6 | - | - | - | - | - | - | - | - | - |
| Validation | 7 | 1 | Optimized-3 | 203.8 | 5 | 8 | 10 | L | 18 | 9 | 0.80 |
| Validation | 7 | 2 | - | - | - | - | - | - | - | - | - |
| Validation | 7 | 3 | Pre-optimized | 98 | 6 | 14 | 10 | S | 8 | 11 | 0.41 |
| Validation | 7 | 4 | Optimized-5 | 226.7 | 6 | 17 | 10 | L | 15 | 11 | 0.78 |
| Validation | 7 | 5 | - | - | - | - | - | - | - | - | - |
| Validation | 7 | 6 | - | - | - | - | - | - | - | - | - |
| Validation | 8 | 1 | Optimized-2 | 203.8 | 5 | 8 | 10 | L | 18 | 11 | 0.70 |
| Validation | 8 | 2 | - | - | - | - | - | - | - | - | - |
| Validation | 8 | 3 | Optimized-1 | 226.7 | 5 | 14 | 10 | L | 17 | 7 | 0.81 |
| Validation | 8 | 4 | Optimized-4 | 226.7 | 6 | 17 | 10 | L | 15 | 12 | 0.77 |
| Validation | 8 | 5 | - | - | - | - | - | - | - | - | - |
| Validation | 8 | 6 | - | - | - | - | - | - | - | - | - |

**Table S5**

| Experiment | DDay | Run | Category | Plate | Start | End | Duration | Number of commands | | | | Error detail |
|---|---|---|---|---|---|---|---|---|---|---|---|---|
| | | | | | | | | Total | Complete | Error | Abort | |
| Baseline | -7 | 1 | Seeding | 4 to 1, 8 to 5 | 2019/02/07 12:22:29 | 2019/02/07 14:10:28 | 1:47:59 | 89 | 89 | 0 | 0 | - |
| Baseline | -6 | 1 | Preconditioning | 1 to 4 | 2019/02/08 15:17:04 | 2019/02/08 17:48:33 | 2:31:29 | 104 | 104 | 0 | 0 | - |
| Baseline | -6 | 2 | Preconditioning | 5 to 8 | 2019/02/08 18:02:44 | 2019/02/08 20:33:09 | 2:30:25 | 104 | 104 | 0 | 0 | - |
| Baseline | -5 | 1 | Preconditioning | 8 to 5 | 2019/02/09 09:00:28 | 2019/02/09 11:30:37 | 2:30:09 | 104 | 104 | 0 | 0 | - |
| Baseline | -5 | 2 | Preconditioning | 4 to 1 | 2019/02/09 11:40:26 | 2019/02/09 14:10:44 | 2:30:18 | 104 | 104 | 0 | 0 | - |
| Baseline | -4 | 1 | Preconditioning | 1 to 4 | 2019/02/10 10:42:05 | 2019/02/10 13:35:55 | 2:53:50 | 104 | 103 | 1 | 0 | Microscope error |
| Baseline | -4 | 2 | Preconditioning | 5 to 8 | 2019/02/10 13:48:43 | 2019/02/10 16:19:28 | 2:30:45 | 104 | 104 | 0 | 0 | - |
| Baseline | -3 | 1 | Preconditioning | 8 to 5 | 2019/02/11 11:33:11 | 2019/02/11 14:04:12 | 2:31:01 | 104 | 104 | 0 | 0 | - |
| Baseline | -3 | 2 | Preconditioning | 4 to 1 | 2019/02/11 15:50:22 | 2019/02/11 18:21:34 | 2:31:12 | 104 | 104 | 0 | 0 | - |
| Baseline | -2 | 1 | Preconditioning | 1 to 4 | 2019/02/12 10:40:20 | 2019/02/12 13:10:57 | 2:30:37 | 104 | 104 | 0 | 0 | - |
| Baseline | -2 | 2 | Preconditioning | 5 to 8 | 2019/02/12 13:23:49 | 2019/02/12 15:54:23 | 2:30:34 | 104 | 104 | 0 | 0 | - |
| Baseline | -1 | 1 | Preconditioning | 8 to 5 | 2019/02/13 11:06:08 | 2019/02/13 13:36:06 | 2:29:58 | 104 | 104 | 0 | 0 | - |
| Baseline | -1 | 2 | Preconditioning | 4 to 1 | 2019/02/13 13:50:46 | 2019/02/13 16:20:19 | 2:29:33 | 104 | 104 | 0 | 0 | - |
| Baseline | 0 | 1 | Passage | 1 to 4 | 2019/02/14 10:15:55 | 2019/02/14 15:58:04 | 5:42:09 | 151 | 151 | 0 | 0 | - |
| Baseline | 0 | 2 | Passage | 5 to 8 | 2019/02/14 16:44:16 | 2019/02/14 22:27:46 | 5:43:30 | 151 | 151 | 0 | 0 | - |
| Baseline | 1 | 1 | RPE differentiation | 1 to 4 | 2019/02/15 12:29:03 | 2019/02/15 14:59:32 | 2:30:29 | 104 | 104 | 0 | 0 | - |
| Baseline | 1 | 2 | RPE differentiation | 5 to 8 | 2019/02/15 15:16:55 | 2019/02/15 17:47:38 | 2:30:43 | 104 | 104 | 0 | 0 | - |
| Baseline | 2 | 1 | RPE differentiation | 8 to 5 | 2019/02/16 10:57:01 | 2019/02/16 13:30:10 | 2:33:09 | 104 | 104 | 0 | 0 | - |
| Baseline | 2 | 2 | RPE differentiation | 4 to 1 | 2019/02/16 13:42:48 | 2019/02/16 16:12:18 | 2:29:30 | 104 | 104 | 0 | 0 | - |
| Baseline | 3 | 1 | RPE differentiation | 1 to 4 | 2019/02/17 10:43:56 | 2019/02/17 13:13:53 | 2:29:57 | 104 | 104 | 0 | 0 | - |
| Baseline | 3 | 2 | RPE differentiation | 5 to 8 | 2019/02/17 13:25:31 | 2019/02/17 15:58:21 | 2:32:50 | 104 | 104 | 0 | 0 | - |
| Baseline | 4 | 1 | RPE differentiation | 8 to 5 | 2019/02/18 10:23:48 | 2019/02/18 12:56:37 | 2:32:49 | 104 | 104 | 0 | 0 | - |
| Baseline | 4 | 2 | RPE differentiation | 4 to 1 | 2019/02/18 13:19:30 | 2019/02/18 15:49:05 | 2:29:35 | 104 | 104 | 0 | 0 | - |
| Baseline | 5 | 1 | RPE differentiation | 1 to 4 | 2019/02/19 10:52:50 | 2019/02/19 13:23:47 | 2:30:57 | 104 | 104 | 0 | 0 | - |
| Baseline | 5 | 2 | RPE differentiation | 5 to 8 | 2019/02/19 13:39:38 | 2019/02/19 16:11:43 | 2:32:05 | 104 | 104 | 0 | 0 | - |
| Baseline | 6 | 1 | RPE differentiation | 8 to 5 | 2019/02/20 10:47:01 | 2019/02/20 13:19:42 | 2:32:41 | 104 | 104 | 0 | 0 | - |
| Baseline | 6 | 2 | RPE differentiation | 4 to 1 | 2019/02/20 13:30:50 | 2019/02/20 16:00:53 | 2:30:03 | 104 | 104 | 0 | 0 | - |
| Baseline | 7 | 1 | RPE differentiation | 1 to 4 | 2019/02/21 11:51:43 | 2019/02/21 14:22:25 | 2:30:42 | 104 | 104 | 0 | 0 | - |
| Baseline | 7 | 2 | RPE differentiation | 5 to 8 | 2019/02/21 14:38:16 | 2019/02/21 17:18:05 | 2:39:49 | 104 | 104 | 0 | 0 | - |
| Baseline | 8 | 1 | RPE differentiation | 8 to 5 | 2019/02/22 10:51:39 | 2019/02/22 13:22:00 | 2:30:21 | 104 | 104 | 0 | 0 | - |
| Baseline | 8 | 2 | RPE differentiation | 4 to 1 | 2019/02/22 13:32:56 | 2019/02/22 16:02:55 | 2:29:59 | 104 | 104 | 0 | 0 | - |
| Baseline | 9 | 1 | RPE differentiation | 1 to 4 | 2019/02/23 11:51:10 | 2019/02/23 14:21:48 | 2:30:38 | 104 | 104 | 0 | 0 | - |
| Baseline | 9 | 2 | RPE differentiation | 5 to 8 | 2019/02/23 14:32:11 | 2019/02/23 17:05:01 | 2:32:50 | 104 | 104 | 0 | 0 | - |
| Baseline | 10 | 1 | RPE differentiation | 8 to 5 | 2019/02/24 11:14:25 | 2019/02/24 13:47:13 | 2:32:48 | 104 | 104 | 0 | 0 | - |
| Baseline | 10 | 2 | RPE differentiation | 4 to 1 | 2019/02/24 13:58:19 | 2019/02/24 16:28:38 | 2:30:19 | 104 | 104 | 0 | 0 | - |
| Baseline | 11 | 1 | RPE differentiation | 1 to 4 | 2019/02/25 10:25:02 | 2019/02/25 12:55:03 | 2:30:01 | 104 | 104 | 0 | 0 | - |
| Baseline | 11 | 2 | RPE differentiation | 5 to 8 | 2019/02/25 13:06:49 | 2019/02/25 15:38:24 | 2:31:35 | 104 | 104 | 0 | 0 | - |
| Baseline | 12 | 1 | RPE differentiation | 8 to 5 | 2019/02/26 10:27:07 | 2019/02/26 12:59:19 | 2:32:12 | 104 | 104 | 0 | 0 | - |
| Baseline | 12 | 2 | RPE differentiation | 4 to 1 | 2019/02/26 13:11:17 | 2019/02/26 15:42:55 | 2:31:38 | 104 | 104 | 0 | 0 | - |
| Baseline | 13 | 1 | RPE differentiation | 1 to 4 | 2019/02/27 10:38:13 | 2019/02/27 13:32:25 | 2:54:12 | 104 | 103 | 1 | 0 | Microscope error |
| Baseline | 13 | 2 | RPE differentiation | 5 to 8 | 2019/02/27 13:46:37 | 2019/02/27 16:23:20 | 2:36:43 | 104 | 103 | 1 | 0 | Micropipet tip ejection error |
| Baseline | 14 | 1 | RPE differentiation | 8 to 5 | 2019/02/28 10:33:47 | 2019/02/28 13:07:34 | 2:33:47 | 104 | 102 | 2 | 0 | Micropipet tip ejection error (x2) |
| Baseline | 14 | 2 | RPE differentiation | 4 to 1 | 2019/02/28 13:21:42 | 2019/02/28 15:57:03 | 2:35:21 | 104 | 101 | 3 | 0 | Micropipet tip ejection error (x3) |
| Baseline | 15 | 1 | RPE differentiation | 1 to 4 | 2019/03/01 11:11:44 | 2019/03/01 13:42:16 | 2:30:32 | 104 | 104 | 0 | 0 | - |
| Baseline | 15 | 2 | RPE differentiation | 5 to 8 | 2019/03/01 13:53:04 | 2019/03/01 16:25:28 | 2:32:24 | 104 | 104 | 0 | 0 | - |
| Baseline | 16 | 1 | RPE differentiation | 8 to 5 | 2019/03/02 11:57:01 | 2019/03/02 14:27:19 | 2:30:18 | 104 | 104 | 0 | 0 | - |
| Baseline | 16 | 2 | RPE differentiation | 4 to 1 | 2019/03/02 14:38:14 | 2019/03/02 17:07:53 | 2:29:39 | 104 | 104 | 0 | 0 | - |
| Baseline | 17 | 1 | RPE differentiation | 1 to 4 | 2019/03/03 10:37:33 | 2019/03/03 13:26:30 | 2:48:57 | 104 | 103 | 1 | 0 | Human error |
| Baseline | 17 | 2 | RPE differentiation | 5 to 8 | 2019/03/03 13:49:29 | 2019/03/03 16:22:08 | 2:32:39 | 104 | 104 | 0 | 0 | - |
| Baseline | 18 | 1 | RPE differentiation | 8 to 5 | 2019/03/04 09:00:02 | 2019/03/04 11:31:58 | 2:31:56 | 104 | 104 | 0 | 0 | - |
| Baseline | 18 | 2 | RPE differentiation | 4 to 1 | 2019/03/04 12:12:20 | 2019/03/04 14:45:17 | 2:32:57 | 104 | 104 | 0 | 0 | - |
| Baseline | 19 | 1 | RPE differentiation | 1 to 4 | 2019/03/05 10:40:30 | 2019/03/05 13:10:05 | 2:29:35 | 104 | 104 | 0 | 0 | - |
| Baseline | 19 | 2 | RPE differentiation | 5 to 8 | 2019/03/05 13:23:54 | 2019/03/05 15:55:27 | 2:31:33 | 104 | 104 | 0 | 0 | - |
| Baseline | 20 | 1 | RPE differentiation | 8 to 5 | 2019/03/06 10:25:10 | 2019/03/06 12:13:36 | 1:48:26 | 77 | 77 | 0 | 0 | - |
| Baseline | 20 | 2 | RPE differentiation | 4 to 1 | 2019/03/06 12:20:49 | 2019/03/06 15:16:26 | 2:55:37 | 77 | 76 | 1 | 0 | HDD for microscope PC error |
| Baseline | 21 | 1 | RPE differentiation | 1 to 4 | 2019/03/07 11:17:57 | 2019/03/07 13:04:43 | 1:46:46 | 77 | 77 | 0 | 0 | - |
| Baseline | 21 | 2 | RPE differentiation | 5 to 8 | 2019/03/07 13:07:17 | 2019/03/07 14:55:11 | 1:47:54 | 77 | 77 | 0 | 0 | - |
| Baseline | 22 | 1 | RPE differentiation | 8 to 5 | 2019/03/08 10:22:27 | 2019/03/08 12:10:50 | 1:48:23 | 77 | 77 | 0 | 0 | - |
| Baseline | 22 | 2 | RPE differentiation | 4 to 1 | 2019/03/08 12:15:10 | 2019/03/08 14:02:38 | 1:47:28 | 77 | 77 | 0 | 0 | - |
| Baseline | 23 | 1 | RPE differentiation | 1 to 4 | 2019/03/09 10:25:05 | 2019/03/09 12:12:25 | 1:47:20 | 77 | 77 | 0 | 0 | - |
| Baseline | 23 | 2 | RPE differentiation | 5 to 8 | 2019/03/09 12:15:54 | 2019/03/09 14:04:59 | 1:49:05 | 77 | 77 | 0 | 0 | - |
| Baseline | 24 | 1 | RPE differentiation | 8 to 5 | 2019/03/10 10:38:53 | 2019/03/10 12:27:04 | 1:48:11 | 77 | 77 | 0 | 0 | - |
| Baseline | 24 | 2 | RPE differentiation | 4 to 1 | 2019/03/10 12:32:18 | 2019/03/10 14:19:17 | 1:46:59 | 77 | 77 | 0 | 0 | - |
| Baseline | 25 | 1 | RPE differentiation | 1 to 4 | 2019/03/11 10:16:04 | 2019/03/11 12:04:50 | 1:48:46 | 77 | 77 | 0 | 0 | - |
| Baseline | 25 | 2 | RPE differentiation | 5 to 8 | 2019/03/11 12:07:34 | 2019/03/11 13:55:20 | 1:47:46 | 77 | 77 | 0 | 0 | - |
| Baseline | 26 | 1 | RPE maintenance | 8 to 5 | 2019/03/12 10:52:50 | 2019/03/12 12:40:53 | 1:48:03 | 77 | 77 | 0 | 0 | - |
| Baseline | 26 | 2 | RPE maintenance | 4 to 1 | 2019/03/12 12:43:34 | 2019/03/12 14:32:14 | 1:48:40 | 77 | 77 | 0 | 0 | - |
| Baseline | 28 | 1 | RPE maintenance | 1 to 4 | 2019/03/14 10:28:18 | 2019/03/14 12:17:38 | 1:49:20 | 77 | 77 | 0 | 0 | - |
| Baseline | 28 | 2 | RPE maintenance | 5 to 8 | 2019/03/14 12:20:12 | 2019/03/14 14:11:09 | 1:50:57 | 77 | 77 | 0 | 0 | - |
| Baseline | 30 | 1 | RPE maintenance | 8 to 5 | 2019/03/16 10:23:16 | 2019/03/16 12:12:15 | 1:48:59 | 77 | 77 | 0 | 0 | - |
| Baseline | 30 | 2 | RPE maintenance | 4 to 1 | 2019/03/16 12:14:59 | 2019/03/16 14:06:20 | 1:51:21 | 77 | 76 | 1 | 0 | Micropipet tip ejection error |
| Baseline | 32 | 1 | RPE maintenance | 1 to 4 | 2019/03/18 11:30:01 | 2019/03/18 13:16:22 | 1:46:21 | 77 | 77 | 0 | 0 | - |
| Baseline | 32 | 2 | RPE maintenance | 5 to 8 | 2019/03/18 13:18:52 | 2019/03/18 15:06:29 | 1:47:37 | 77 | 77 | 0 | 0 | - |
| Round 1 | -7 | 1 | Seeding | 4 to 1, 8 to 5 | 2019/04/18 12:39:42 | 2019/04/18 14:27:46 | 1:48:04 | 89 | 89 | 0 | 0 | - |
| Round 1 | -6 | 1 | Preconditioning | 1 to 4 | 2019/04/19 13:05:56 | 2019/04/19 15:36:58 | 2:31:02 | 193 | 193 | 0 | 0 | - |
| Round 1 | -6 | 2 | Preconditioning | 5 to 8 | 2019/04/19 15:47:38 | 2019/04/19 18:06:44 | 2:19:06 | 193 | 193 | 0 | 0 | - |
| Round 1 | -5 | 1 | Preconditioning | 8 to 5 | 2019/04/20 11:18:42 | 2019/04/20 13:48:48 | 2:30:06 | 193 | 193 | 0 | 0 | - |
| Round 1 | -5 | 2 | Preconditioning | 4 to 1 | 2019/04/20 14:16:38 | 2019/04/20 16:59:46 | 2:43:08 | 193 | 192 | 1 | 0 | Micropipet tip loading error |
| Round 1 | -4 | 1 | Preconditioning | 1 to 4 | 2019/04/21 10:43:17 | 2019/04/21 13:23:02 | 2:39:45 | 193 | 193 | 0 | 0 | - |
| Round 1 | -4 | 2 | Preconditioning | 5 to 8 | 2019/04/21 13:35:04 | 2019/04/21 16:09:11 | 2:34:07 | 193 | 193 | 0 | 0 | - |
| Round 1 | -3 | 1 | Preconditioning | 8 to 5 | 2019/04/22 10:40:37 | 2019/04/22 13:19:21 | 2:38:44 | 193 | 193 | 0 | 0 | - |
| Round 1 | -3 | 2 | Preconditioning | 4 to 1 | 2019/04/22 13:32:35 | 2019/04/22 16:18:44 | 2:46:09 | 193 | 193 | 0 | 0 | - |
| Round 1 | -2 | 1 | Preconditioning | 1 to 4 | 2019/04/23 10:53:46 | 2019/04/23 13:43:26 | 2:49:40 | 193 | 193 | 0 | 0 | - |
| Round 1 | -2 | 2 | Preconditioning | 5 to 8 | 2019/04/23 13:56:40 | 2019/04/23 16:42:29 | 2:45:49 | 193 | 193 | 0 | 0 | - |
| Round 1 | -1 | 1 | Preconditioning | 8 to 5 | 2019/04/24 10:55:04 | 2019/04/24 13:46:14 | 2:51:10 | 193 | 193 | 0 | 0 | - |
| Round 1 | -1 | 2 | Preconditioning | 4 to 1 | 2019/04/24 13:59:33 | 2019/04/24 16:46:11 | 2:46:38 | 193 | 193 | 0 | 0 | - |
| Round 1 | 0 | 1 | Passage | 1 to 4 | 2019/04/25 10:00:39 | 2019/04/25 15:44:39 | 5:44:00 | 151 | 151 | 0 | 0 | - |
| Round 1 | 0 | 2 | Passage | 5 to 8 | 2019/04/25 16:31:28 | 2019/04/25 22:14:27 | 5:42:59 | 151 | 151 | 0 | 0 | - |
| Round 1 | 1 | 1 | RPE differentiation | 1 to 4 | 2019/04/26 11:03:13 | 2019/04/26 14:53:11 | 3:49:58 | 335 | 335 | 0 | 0 | - |
| Round 1 | 1 | 2 | RPE differentiation | 5 to 8 | 2019/04/26 15:14:08 | 2019/04/26 19:09:21 | 3:55:13 | 335 | 335 | 0 | 0 | - |
| Round 1 | 2 | 1 | RPE differentiation | 8 to 5 | 2019/04/27 10:45:19 | 2019/04/27 14:38:28 | 3:53:09 | 335 | 335 | 0 | 0 | - |
| Round 1 | 2 | 2 | RPE differentiation | 4 to 1 | 2019/04/27 14:56:29 | 2019/04/27 18:44:53 | 3:48:24 | 335 | 335 | 0 | 0 | - |
| Round 1 | 3 | 1 | RPE differentiation | 1 to 4 | 2019/04/28 11:06:14 | 2019/04/28 14:51:31 | 3:45:17 | 335 | 335 | 0 | 0 | - |
| Round 1 | 3 | 2 | RPE differentiation | 5 to 8 | 2019/04/28 15:11:37 | 2019/04/28 19:00:29 | 3:48:52 | 335 | 335 | 0 | 0 | - |
| Round 1 | 4 | 1 | RPE differentiation | 8 to 5 | 2019/04/29 10:35:44 | 2019/04/29 14:19:20 | 3:43:36 | 335 | 335 | 0 | 0 | - |
| Round 1 | 4 | 2 | RPE differentiation | 4 to 1 | 2019/04/29 14:34:45 | 2019/04/29 18:21:40 | 3:46:55 | 335 | 334 | 1 | 0 | Micropipet tip loading error |
| Round 1 | 5 | 1 | RPE differentiation | 1 to 4 | 2019/04/30 10:23:46 | 2019/04/30 14:00:16 | 3:36:30 | 335 | 335 | 0 | 0 | - |
| Round 1 | 5 | 2 | RPE differentiation | 5 to 8 | 2019/04/30 14:22:26 | 2019/04/30 17:58:38 | 3:36:12 | 335 | 334 | 1 | 0 | Micropipet tip loading error |
| Round 1 | 6 | 1 | RPE differentiation | 8 to 5 | 2019/05/01 10:20:57 | 2019/05/01 13:51:48 | 3:30:51 | 335 | 335 | 0 | 0 | - |
| Round 1 | 6 | 2 | RPE differentiation | 4 to 1 | 2019/05/01 14:07:19 | 2019/05/01 17:44:17 | 3:36:58 | 335 | 335 | 0 | 0 | - |
| Round 1 | 7 | 1 | RPE differentiation | 1 to 4 | 2019/05/02 11:49:40 | 2019/05/02 15:23:40 | 3:34:00 | 335 | 335 | 0 | 0 | - |
| Round 1 | 7 | 2 | RPE differentiation | 5 to 8 | 2019/05/02 15:38:03 | 2019/05/02 19:46:03 | 4:08:00 | 335 | 334 | 1 | 0 | Microscope error |
| Round 1 | 8 | 1 | RPE differentiation | 8 to 5 | 2019/05/03 11:10:58 | 2019/05/03 14:36:35 | 3:25:37 | 335 | 335 | 0 | 0 | - |
| Round 1 | 8 | 2 | RPE differentiation | 4 to 1 | 2019/05/03 14:56:27 | 2019/05/03 18:24:53 | 3:28:26 | 335 | 335 | 0 | 0 | - |
| Round 1 | 9 | 1 | RPE differentiation | 1 to 4 | 2019/05/04 12:27:22 | 2019/05/04 15:52:27 | 3:25:05 | 335 | 335 | 0 | 0 | - |
| Round 1 | 9 | 2 | RPE differentiation | 5 to 8 | 2019/05/04 16:06:13 | 2019/05/04 19:26:36 | 3:20:23 | 335 | 334 | 1 | 0 | Micropipet tip loading error |
| Round 1 | 10 | 1 | RPE differentiation | 8 to 5 | 2019/05/05 11:22:46 | 2019/05/05 14:39:21 | 3:16:35 | 335 | 335 | 0 | 0 | - |
| Round 1 | 10 | 2 | RPE differentiation | 4 to 1 | 2019/05/05 14:53:39 | 2019/05/05 18:13:57 | 3:20:18 | 335 | 335 | 0 | 0 | - |
| Round 1 | 11 | 1 | RPE differentiation | 1 to 4 | 2019/05/06 11:03:43 | 2019/05/06 14:18:02 | 3:14:19 | 335 | 335 | 0 | 0 | - |
| Round 1 | 11 | 2 | RPE differentiation | 5 to 8 | 2019/05/06 14:34:48 | 2019/05/06 17:48:26 | 3:13:38 | 335 | 335 | 0 | 0 | - |
| Round 1 | 12 | 1 | RPE differentiation | 8 to 5 | 2019/05/07 10:46:31 | 2019/05/07 13:54:26 | 3:07:55 | 335 | 335 | 0 | 0 | - |
| Round 1 | 12 | 2 | RPE differentiation | 4 to 1 | 2019/05/07 14:16:57 | 2019/05/07 17:22:22 | 3:05:25 | 335 | 335 | 0 | 0 | - |
| Round 1 | 13 | 1 | RPE differentiation | 1 to 4 | 2019/05/08 11:09:22 | 2019/05/08 14:10:08 | 3:00:46 | 335 | 335 | 0 | 0 | - |
| Round 1 | 13 | 2 | RPE differentiation | 5 to 8 | 2019/05/08 14:23:51 | 2019/05/08 17:28:12 | 3:04:21 | 335 | 335 | 0 | 0 | - |
| Round 1 | 14 | 1 | RPE differentiation | 8 to 5 | 2019/05/09 11:00:10 | 2019/05/09 13:58:53 | 2:58:43 | 335 | 335 | 0 | 0 | - |
| Round 1 | 14 | 2 | RPE differentiation | 4 to 1 | 2019/05/09 14:12:53 | 2019/05/09 17:10:04 | 2:57:11 | 335 | 335 | 0 | 0 | - |
| Round 1 | 15 | 1 | RPE differentiation | 1 to 4 | 2019/05/10 11:28:45 | 2019/05/10 14:23:09 | 2:54:24 | 335 | 335 | 0 | 0 | - |
| Round 1 | 15 | 2 | RPE differentiation | 5 to 8 | 2019/05/10 14:40:10 | 2019/05/10 17:35:10 | 2:55:00 | 335 | 335 | 0 | 0 | - |

| | | | | | | | | | | | | |
|---|---|---|---|---|---|---|---|---|---|---|---|---|
| Round 1 | 16 | 1 | RPE differentiation | 8 to 5 | 2019/05/11 11:09:12 | 2019/05/11 14:02:03 | 2:52:51 | 335 | 335 | 0 | 0 | - |
| Round 1 | 16 | 2 | RPE differentiation | 4 to 1 | 2019/05/11 14:14:49 | 2019/05/11 17:04:09 | 2:49:20 | 335 | 335 | 0 | 0 | - |
| Round 1 | 17 | 1 | RPE differentiation | 1 to 4 | 2019/05/12 11:17:24 | 2019/05/12 14:00:06 | 2:42:42 | 335 | 335 | 0 | 0 | - |
| Round 1 | 17 | 2 | RPE differentiation | 5 to 8 | 2019/05/12 14:14:23 | 2019/05/12 17:02:58 | 2:48:35 | 335 | 335 | 0 | 0 | - |
| Round 1 | 18 | 1 | RPE differentiation | 8 to 5 | 2019/05/13 11:14:38 | 2019/05/13 13:54:07 | 2:39:29 | 335 | 335 | 0 | 0 | - |
| Round 1 | 18 | 2 | RPE differentiation | 4 to 1 | 2019/05/13 14:08:24 | 2019/05/13 16:51:46 | 2:43:22 | 335 | 335 | 0 | 0 | - |
| Round 1 | 19 | 1 | RPE differentiation | 1 to 4 | 2019/05/14 10:47:52 | 2019/05/14 13:28:10 | 2:40:18 | 335 | 335 | 0 | 0 | - |
| Round 1 | 19 | 2 | RPE differentiation | 5 to 8 | 2019/05/14 13:39:32 | 2019/05/14 16:17:52 | 2:38:20 | 335 | 335 | 0 | 0 | - |
| Round 1 | 20 | 1 | RPE differentiation | 8 to 5 | 2019/05/15 10:41:33 | 2019/05/15 12:49:04 | 2:07:31 | 77 | 76 | 1 | 0 | Microscope error |
| Round 1 | 20 | 2 | RPE differentiation | 4 to 1 | 2019/05/15 12:52:38 | 2019/05/15 14:40:33 | 1:47:55 | 77 | 77 | 0 | 0 | - |
| Round 1 | 21 | 1 | RPE differentiation | 1 to 4 | 2019/05/16 10:49:00 | 2019/05/16 12:37:02 | 1:48:02 | 77 | 77 | 0 | 0 | - |
| Round 1 | 21 | 2 | RPE differentiation | 5 to 8 | 2019/05/16 12:39:44 | 2019/05/16 14:27:38 | 1:47:54 | 77 | 77 | 0 | 0 | - |
| Round 1 | 22 | 1 | RPE differentiation | 8 to 5 | 2019/05/17 10:36:09 | 2019/05/17 12:24:33 | 1:48:24 | 77 | 77 | 0 | 0 | - |
| Round 1 | 22 | 2 | RPE differentiation | 4 to 1 | 2019/05/17 12:32:36 | 2019/05/17 14:21:25 | 1:48:49 | 77 | 77 | 0 | 0 | - |
| Round 1 | 23 | 1 | RPE differentiation | 1 to 4 | 2019/05/18 11:45:26 | 2019/05/18 13:33:01 | 1:47:35 | 77 | 77 | 0 | 0 | - |
| Round 1 | 23 | 2 | RPE differentiation | 5 to 8 | 2019/05/18 13:57:57 | 2019/05/18 15:46:24 | 1:48:27 | 77 | 77 | 0 | 0 | - |
| Round 1 | 24 | 1 | RPE differentiation | 8 to 5 | 2019/05/19 12:10:03 | 2019/05/19 13:57:52 | 1:47:49 | 77 | 77 | 0 | 0 | - |
| Round 1 | 24 | 2 | RPE differentiation | 4 to 1 | 2019/05/19 14:09:03 | 2019/05/19 15:57:58 | 1:48:55 | 77 | 77 | 0 | 0 | - |
| Round 1 | 25 | 1 | RPE differentiation | 1 to 4 | 2019/05/20 10:23:49 | 2019/05/20 13:22:28 | 2:58:39 | 77 | 73 | 1 | 3 | HDD for microscope PC error |
| Round 1 | 25 | 2 | RPE differentiation | 5 to 8 | 2019/05/20 14:33:42 | 2019/05/20 16:06:50 | 1:33:08 | 77 | 76 | 1 | 0 | HDD for microscope PC error |
| Round 1 | 26 | 1 | RPE maintenance | 8 to 5 | 2019/05/21 10:43:02 | 2019/05/21 12:30:52 | 1:47:50 | 77 | 77 | 0 | 0 | - |
| Round 1 | 26 | 2 | RPE maintenance | 4 to 1 | 2019/05/21 12:35:08 | 2019/05/21 14:23:15 | 1:48:07 | 77 | 77 | 0 | 0 | - |
| Round 1 | 28 | 1 | RPE maintenance | 1 to 4 | 2019/05/23 10:27:32 | 2019/05/23 12:15:19 | 1:47:47 | 77 | 77 | 0 | 0 | - |
| Round 1 | 28 | 2 | RPE maintenance | 5 to 8 | 2019/05/23 12:20:06 | 2019/05/23 14:02:00 | 1:41:54 | 77 | 77 | 0 | 0 | - |
| Round 1 | 30 | 1 | RPE maintenance | 8 to 5 | 2019/05/25 11:19:43 | 2019/05/25 13:07:38 | 1:47:55 | 77 | 77 | 0 | 0 | - |
| Round 1 | 30 | 2 | RPE maintenance | 4 to 1 | 2019/05/25 13:20:54 | 2019/05/25 15:08:39 | 1:47:45 | 77 | 77 | 0 | 0 | - |
| Round 1 | 32 | 1 | RPE maintenance | 1 to 4 | 2019/05/27 11:20:59 | 2019/05/27 13:08:24 | 1:47:25 | 77 | 77 | 0 | 0 | - |
| Round 1 | 32 | 2 | RPE maintenance | 5 to 8 | 2019/05/27 13:11:12 | 2019/05/27 14:58:17 | 1:47:05 | 77 | 77 | 0 | 0 | - |
| Round 2 | -7 | 1 | Seeding | 4 to 1, 8 to 5 | 2019/07/05 12:13:58 | 2019/07/05 14:02:26 | 1:48:28 | 89 | 89 | 0 | 0 | - |
| Round 2 | -6 | 1 | Preconditioning | 1 to 4 | 2019/07/06 12:37:47 | 2019/07/06 15:29:57 | 2:52:10 | 193 | 193 | 0 | 0 | - |
| Round 2 | -6 | 2 | Preconditioning | 5 to 8 | 2019/07/06 15:42:20 | 2019/07/06 18:34:12 | 2:51:52 | 193 | 193 | 0 | 0 | - |
| Round 2 | -5 | 1 | Preconditioning | 8 to 5 | 2019/07/07 11:59:55 | 2019/07/07 14:52:32 | 2:52:37 | 193 | 193 | 0 | 0 | - |
| Round 2 | -5 | 2 | Preconditioning | 4 to 1 | 2019/07/07 15:06:29 | 2019/07/07 18:06:35 | 3:00:06 | 193 | 193 | 0 | 0 | - |
| Round 2 | -4 | 1 | Preconditioning | 1 to 4 | 2019/07/08 10:50:01 | 2019/07/08 14:02:48 | 3:12:47 | 193 | 193 | 0 | 0 | - |
| Round 2 | -4 | 2 | Preconditioning | 5 to 8 | 2019/07/08 14:16:30 | 2019/07/08 17:10:29 | 2:53:59 | 193 | 193 | 0 | 0 | - |
| Round 2 | -3 | 1 | Preconditioning | 8 to 5 | 2019/07/09 10:51:04 | 2019/07/09 13:47:44 | 2:56:40 | 193 | 193 | 0 | 0 | - |
| Round 2 | -3 | 2 | Preconditioning | 4 to 1 | 2019/07/09 14:01:09 | 2019/07/09 17:47:02 | 3:45:53 | 193 | 192 | 1 | 0 | Microscope error |
| Round 2 | -2 | 1 | Preconditioning | 1 to 4 | 2019/07/10 10:45:28 | 2019/07/10 14:22:52 | 3:37:24 | 193 | 192 | 1 | 0 | Microscope error |
| Round 2 | -2 | 2 | Preconditioning | 5 to 8 | 2019/07/10 14:35:27 | 2019/07/10 17:34:50 | 2:59:23 | 193 | 193 | 0 | 0 | - |
| Round 2 | -1 | 1 | Preconditioning | 8 to 5 | 2019/07/11 11:11:05 | 2019/07/11 14:05:03 | 2:53:58 | 193 | 193 | 0 | 0 | - |
| Round 2 | -1 | 2 | Preconditioning | 4 to 1 | 2019/07/11 14:19:25 | 2019/07/11 17:50:07 | 3:30:42 | 193 | 193 | 0 | 0 | - |
| Round 2 | 0 | 1 | Passage | 1 to 4 | 2019/07/12 09:52:09 | 2019/07/12 15:32:04 | 5:39:55 | 151 | 151 | 0 | 0 | - |
| Round 2 | 0 | 2 | Passage | 5 to 8 | 2019/07/12 16:21:53 | 2019/07/12 22:01:45 | 5:39:52 | 151 | 151 | 0 | 0 | - |
| Round 2 | 1 | 1 | RPE differentiation | 1 to 4 | 2019/07/13 11:39:30 | 2019/07/13 15:34:40 | 3:55:10 | 335 | 335 | 0 | 0 | - |
| Round 2 | 1 | 2 | RPE differentiation | 5 to 8 | 2019/07/13 15:51:55 | 2019/07/13 19:39:46 | 3:47:51 | 335 | 335 | 0 | 0 | - |
| Round 2 | 2 | 1 | RPE differentiation | 8 to 5 | 2019/07/14 11:42:19 | 2019/07/14 15:29:58 | 3:47:39 | 335 | 335 | 0 | 0 | - |
| Round 2 | 2 | 2 | RPE differentiation | 4 to 1 | 2019/07/14 15:46:01 | 2019/07/14 19:39:59 | 3:53:58 | 335 | 335 | 0 | 0 | - |
| Round 2 | 3 | 1 | RPE differentiation | 1 to 4 | 2019/07/15 11:58:25 | 2019/07/15 15:48:11 | 3:49:46 | 335 | 335 | 0 | 0 | - |
| Round 2 | 3 | 2 | RPE differentiation | 5 to 8 | 2019/07/15 16:23:15 | 2019/07/15 20:07:39 | 3:44:24 | 335 | 335 | 0 | 0 | - |
| Round 2 | 4 | 1 | RPE differentiation | 8 to 5 | 2019/07/16 11:48:14 | 2019/07/16 15:25:54 | 3:37:40 | 335 | 335 | 0 | 0 | - |
| Round 2 | 4 | 2 | RPE differentiation | 4 to 1 | 2019/07/16 15:44:54 | 2019/07/16 19:38:34 | 3:53:40 | 335 | 335 | 0 | 0 | - |
| Round 2 | 5 | 1 | RPE differentiation | 1 to 4 | 2019/07/17 12:02:00 | 2019/07/17 15:41:33 | 3:39:33 | 335 | 335 | 0 | 0 | - |
| Round 2 | 5 | 2 | RPE differentiation | 5 to 8 | 2019/07/17 15:59:59 | 2019/07/17 19:33:51 | 3:33:52 | 335 | 335 | 0 | 0 | - |
| Round 2 | 6 | 1 | RPE differentiation | 8 to 5 | 2019/07/18 11:14:42 | 2019/07/18 14:42:16 | 3:27:34 | 335 | 335 | 0 | 0 | - |
| Round 2 | 6 | 2 | RPE differentiation | 4 to 1 | 2019/07/18 15:11:44 | 2019/07/18 18:51:06 | 3:39:22 | 335 | 335 | 0 | 0 | - |
| Round 2 | 7 | 1 | RPE differentiation | 1 to 4 | 2019/07/19 11:13:50 | 2019/07/19 14:56:27 | 3:42:37 | 335 | 335 | 0 | 0 | - |
| Round 2 | 7 | 2 | RPE differentiation | 5 to 8 | 2019/07/19 15:16:34 | 2019/07/19 18:43:52 | 3:27:18 | 335 | 335 | 0 | 0 | - |
| Round 2 | 8 | 1 | RPE differentiation | 8 to 5 | 2019/07/20 11:29:35 | 2019/07/20 14:57:45 | 3:28:10 | 335 | 335 | 0 | 0 | - |
| Round 2 | 8 | 2 | RPE differentiation | 4 to 1 | 2019/07/20 15:50:29 | 2019/07/20 19:23:04 | 3:32:35 | 335 | 335 | 0 | 0 | - |
| Round 2 | 9 | 1 | RPE differentiation | 1 to 4 | 2019/07/21 11:29:51 | 2019/07/21 14:59:10 | 3:29:19 | 335 | 335 | 0 | 0 | - |
| Round 2 | 9 | 2 | RPE differentiation | 5 to 8 | 2019/07/21 15:20:13 | 2019/07/21 18:41:38 | 3:21:25 | 335 | 335 | 0 | 0 | - |
| Round 2 | 10 | 1 | RPE differentiation | 8 to 5 | 2019/07/22 11:23:57 | 2019/07/22 14:42:57 | 3:19:00 | 335 | 335 | 0 | 0 | - |
| Round 2 | 10 | 2 | RPE differentiation | 4 to 1 | 2019/07/22 14:57:36 | 2019/07/22 18:29:12 | 3:31:36 | 335 | 335 | 0 | 0 | - |
| Round 2 | 11 | 1 | RPE differentiation | 1 to 4 | 2019/07/23 11:06:03 | 2019/07/23 14:40:34 | 3:34:31 | 335 | 334 | 1 | 0 | Human error |
| Round 2 | 11 | 2 | RPE differentiation | 5 to 8 | 2019/07/23 15:02:19 | 2019/07/23 18:13:29 | 3:11:10 | 335 | 335 | 0 | 0 | - |
| Round 2 | 12 | 1 | RPE differentiation | 8 to 5 | 2019/07/24 11:08:29 | 2019/07/24 14:21:30 | 3:13:01 | 335 | 335 | 0 | 0 | - |
| Round 2 | 12 | 2 | RPE differentiation | 4 to 1 | 2019/07/24 14:38:04 | 2019/07/24 17:56:27 | 3:18:23 | 335 | 335 | 0 | 0 | - |
| Round 2 | 13 | 1 | RPE differentiation | 1 to 4 | 2019/07/25 11:50:41 | 2019/07/25 15:07:52 | 3:17:11 | 335 | 335 | 0 | 0 | - |
| Round 2 | 13 | 2 | RPE differentiation | 5 to 8 | 2019/07/25 15:32:25 | 2019/07/25 18:39:43 | 3:07:18 | 335 | 335 | 0 | 0 | - |
| Round 2 | 14 | 1 | RPE differentiation | 8 to 5 | 2019/07/26 11:04:57 | 2019/07/26 14:11:04 | 3:06:07 | 335 | 335 | 0 | 0 | - |
| Round 2 | 14 | 2 | RPE differentiation | 4 to 1 | 2019/07/26 14:26:17 | 2019/07/26 17:38:10 | 3:11:53 | 335 | 335 | 0 | 0 | - |
| Round 2 | 15 | 1 | RPE differentiation | 1 to 4 | 2019/07/27 11:13:59 | 2019/07/27 14:24:03 | 3:10:04 | 335 | 335 | 0 | 0 | - |
| Round 2 | 15 | 2 | RPE differentiation | 5 to 8 | 2019/07/27 14:44:33 | 2019/07/27 17:49:16 | 3:04:43 | 335 | 335 | 0 | 0 | - |
| Round 2 | 16 | 1 | RPE differentiation | 8 to 5 | 2019/07/28 11:06:32 | 2019/07/28 14:19:05 | 3:12:33 | 335 | 334 | 1 | 0 | Human error |
| Round 2 | 16 | 2 | RPE differentiation | 4 to 1 | 2019/07/28 14:33:43 | 2019/07/28 17:37:40 | 3:03:57 | 335 | 335 | 0 | 0 | - |
| Round 2 | 17 | 1 | RPE differentiation | 1 to 4 | 2019/07/29 13:00:38 | 2019/07/29 16:02:53 | 3:02:15 | 335 | 335 | 0 | 0 | - |
| Round 2 | 17 | 2 | RPE differentiation | 5 to 8 | 2019/07/29 16:17:44 | 2019/07/29 19:12:41 | 2:54:57 | 335 | 335 | 0 | 0 | - |
| Round 2 | 18 | 1 | RPE differentiation | 8 to 5 | 2019/07/30 11:15:06 | 2019/07/30 14:00:02 | 2:44:56 | 335 | 335 | 0 | 0 | - |
| Round 2 | 18 | 2 | RPE differentiation | 4 to 1 | 2019/07/30 14:14:24 | 2019/07/30 17:12:07 | 2:57:43 | 335 | 335 | 0 | 0 | - |
| Round 2 | 19 | 1 | RPE differentiation | 1 to 4 | 2019/07/31 11:08:58 | 2019/07/31 13:51:09 | 2:42:11 | 335 | 335 | 0 | 0 | - |
| Round 2 | 19 | 2 | RPE differentiation | 5 to 8 | 2019/07/31 14:03:03 | 2019/07/31 16:44:12 | 2:41:09 | 335 | 335 | 0 | 0 | - |
| Round 2 | 20 | 1 | RPE differentiation | 8 to 5 | 2019/08/01 11:15:57 | 2019/08/01 13:03:06 | 1:47:09 | 77 | 77 | 0 | 0 | - |
| Round 2 | 20 | 2 | RPE differentiation | 4 to 1 | 2019/08/01 13:06:24 | 2019/08/01 14:54:13 | 1:47:49 | 77 | 77 | 0 | 0 | - |
| Round 2 | 21 | 1 | RPE differentiation | 1 to 4 | 2019/08/02 11:14:00 | 2019/08/02 13:02:07 | 1:48:07 | 77 | 77 | 0 | 0 | - |
| Round 2 | 21 | 2 | RPE differentiation | 5 to 8 | 2019/08/02 13:04:57 | 2019/08/02 14:53:15 | 1:48:18 | 77 | 77 | 0 | 0 | - |
| Round 2 | 22 | 1 | RPE differentiation | 8 to 5 | 2019/08/03 12:14:11 | 2019/08/03 14:02:17 | 1:48:06 | 77 | 77 | 0 | 0 | - |
| Round 2 | 22 | 2 | RPE differentiation | 4 to 1 | 2019/08/03 14:27:08 | 2019/08/03 16:15:12 | 1:48:04 | 77 | 77 | 0 | 0 | - |
| Round 2 | 23 | 1 | RPE differentiation | 1 to 4 | 2019/08/04 11:19:55 | 2019/08/04 13:08:08 | 1:48:13 | 77 | 77 | 0 | 0 | - |
| Round 2 | 23 | 2 | RPE differentiation | 5 to 8 | 2019/08/04 13:11:15 | 2019/08/04 14:59:14 | 1:47:59 | 77 | 77 | 0 | 0 | - |
| Round 2 | 24 | 1 | RPE differentiation | 8 to 5 | 2019/08/05 11:15:04 | 2019/08/05 13:03:23 | 1:48:19 | 77 | 77 | 0 | 0 | - |
| Round 2 | 24 | 2 | RPE differentiation | 4 to 1 | 2019/08/05 13:07:05 | 2019/08/05 14:55:22 | 1:48:17 | 77 | 77 | 0 | 0 | - |
| Round 2 | 25 | 1 | RPE differentiation | 1 to 4 | 2019/08/06 11:10:12 | 2019/08/06 12:58:22 | 1:48:10 | 77 | 77 | 0 | 0 | - |
| Round 2 | 25 | 2 | RPE differentiation | 5 to 8 | 2019/08/06 13:02:11 | 2019/08/06 14:50:27 | 1:48:16 | 77 | 77 | 0 | 0 | - |
| Round 2 | 26 | 1 | RPE maintenance | 8 to 5 | 2019/08/07 11:17:07 | 2019/08/07 13:05:18 | 1:48:11 | 77 | 77 | 0 | 0 | - |
| Round 2 | 26 | 2 | RPE maintenance | 4 to 1 | 2019/08/07 13:08:59 | 2019/08/07 14:56:37 | 1:47:38 | 77 | 77 | 0 | 0 | - |
| Round 2 | 28 | 1 | RPE maintenance | 1 to 4 | 2019/08/09 11:07:57 | 2019/08/09 12:56:19 | 1:48:22 | 77 | 77 | 0 | 0 | - |
| Round 2 | 28 | 2 | RPE maintenance | 5 to 8 | 2019/08/09 13:00:08 | 2019/08/09 14:48:23 | 1:48:15 | 77 | 77 | 0 | 0 | - |
| Round 2 | 30 | 1 | RPE maintenance | 8 to 5 | 2019/08/11 11:36:18 | 2019/08/11 13:24:17 | 1:47:59 | 77 | 77 | 0 | 0 | - |
| Round 2 | 30 | 2 | RPE maintenance | 4 to 1 | 2019/08/11 13:28:48 | 2019/08/11 15:16:20 | 1:47:32 | 77 | 77 | 0 | 0 | - |
| Round 2 | 32 | 1 | RPE maintenance | 1 to 4 | 2019/08/13 11:01:08 | 2019/08/13 12:48:29 | 1:47:21 | 77 | 77 | 0 | 0 | - |
| Round 2 | 32 | 2 | RPE maintenance | 5 to 8 | 2019/08/13 12:51:17 | 2019/08/13 14:39:31 | 1:48:14 | 77 | 77 | 0 | 0 | - |
| Round 3 | -7 | 1 | Seeding | 4 to 1, 8 to 5 | 2020/01/23 12:14:11 | 2020/01/23 14:02:42 | 1:48:31 | 89 | 89 | 0 | 0 | - |
| Round 3 | -6 | 1 | Preconditioning | 1 to 4 | 2020/01/24 12:17:19 | 2020/01/24 14:35:29 | 2:18:10 | 193 | 193 | 0 | 0 | - |
| Round 3 | -6 | 2 | Preconditioning | 5 to 8 | 2020/01/24 14:51:09 | 2020/01/24 17:11:29 | 2:20:20 | 193 | 193 | 0 | 0 | - |
| Round 3 | -5 | 1 | Preconditioning | 8 to 5 | 2020/01/25 10:30:07 | 2020/01/25 13:17:00 | 2:46:53 | 193 | 193 | 0 | 0 | - |
| Round 3 | -5 | 2 | Preconditioning | 4 to 1 | 2020/01/25 13:30:34 | 2020/01/25 16:19:06 | 2:48:32 | 193 | 193 | 0 | 0 | - |
| Round 3 | -4 | 1 | Preconditioning | 1 to 4 | 2020/01/26 12:26:20 | 2020/01/26 15:15:04 | 2:48:44 | 193 | 193 | 0 | 0 | - |
| Round 3 | -4 | 2 | Preconditioning | 5 to 8 | 2020/01/26 15:42:40 | 2020/01/26 18:29:27 | 2:46:47 | 193 | 193 | 0 | 0 | - |
| Round 3 | -3 | 1 | Preconditioning | 8 to 5 | 2020/01/27 11:20:00 | 2020/01/27 14:06:37 | 2:46:37 | 193 | 193 | 0 | 0 | - |
| Round 3 | -3 | 2 | Preconditioning | 4 to 1 | 2020/01/27 14:36:26 | 2020/01/27 17:24:48 | 2:48:22 | 193 | 193 | 0 | 0 | - |
| Round 3 | -2 | 1 | Preconditioning | 1 to 4 | 2020/01/28 10:47:01 | 2020/01/28 13:35:08 | 2:48:07 | 193 | 193 | 0 | 0 | - |
| Round 3 | -2 | 2 | Preconditioning | 5 to 8 | 2020/01/28 13:52:28 | 2020/01/28 16:39:04 | 2:46:36 | 193 | 193 | 0 | 0 | - |
| Round 3 | -1 | 1 | Preconditioning | 8 to 5 | 2020/01/29 10:34:00 | 2020/01/29 13:20:50 | 2:46:50 | 193 | 193 | 0 | 0 | - |
| Round 3 | -1 | 2 | Preconditioning | 4 to 1 | 2020/01/29 13:38:20 | 2020/01/29 16:26:34 | 2:48:14 | 193 | 193 | 0 | 0 | - |
| Round 3 | 0 | 1 | Passage | 1 to 4 | 2020/01/30 10:12:02 | 2020/01/30 15:03:27 | 4:51:25 | 151 | 151 | 0 | 0 | - |
| Round 3 | 0 | 2 | Passage | 5 to 8 | 2020/01/30 16:08:40 | 2020/01/30 20:59:38 | 4:50:58 | 151 | 151 | 0 | 0 | - |
| Round 3 | 1 | 1 | RPE differentiation | 1 to 4 | 2020/01/31 11:30:25 | 2020/01/31 15:16:44 | 3:46:19 | 335 | 335 | 0 | 0 | - |
| Round 3 | 1 | 2 | RPE differentiation | 5 to 8 | 2020/01/31 15:45:29 | 2020/01/31 19:30:05 | 3:44:36 | 335 | 335 | 0 | 0 | - |
| Round 3 | 2 | 1 | RPE differentiation | 8 to 5 | 2020/02/01 10:09:19 | 2020/02/01 13:52:04 | 3:42:45 | 335 | 335 | 0 | 0 | - |
| Round 3 | 2 | 2 | RPE differentiation | 4 to 1 | 2020/02/01 14:11:49 | 2020/02/01 17:55:03 | 3:43:14 | 335 | 335 | 0 | 0 | - |
| Round 3 | 3 | 1 | RPE differentiation | 1 to 4 | 2020/02/02 11:38:24 | 2020/02/02 15:20:01 | 3:41:37 | 335 | 335 | 0 | 0 | - |
| Round 3 | 3 | 2 | RPE differentiation | 5 to 8 | 2020/02/02 16:01:22 | 2020/02/02 19:46:28 | 3:45:06 | 335 | 335 | 0 | 0 | - |
| Round 3 | 4 | 1 | RPE differentiation | 8 to 5 | 2020/02/03 12:10:35 | 2020/02/03 15:53:17 | 3:42:42 | 335 | 335 | 0 | 0 | - |
| Round 3 | 4 | 2 | RPE differentiation | 4 to 1 | 2020/02/03 16:12:03 | 2020/02/03 20:01:30 | 3:49:27 | 335 | 335 | 0 | 0 | - |
| Round 3 | 5 | 1 | RPE differentiation | 1 to 4 | 2020/02/04 10:48:49 | 2020/02/04 14:32:21 | 3:43:32 | 335 | 335 | 0 | 0 | - |
| Round 3 | 5 | 2 | RPE differentiation | 5 to 8 | 2020/02/04 14:51:07 | 2020/02/04 18:26:28 | 3:35:21 | 335 | 335 | 0 | 0 | - |
| Round 3 | 6 | 1 | RPE differentiation | 8 to 5 | 2020/02/05 10:47:33 | 2020/02/05 14:15:17 | 3:27:44 | 335 | 335 | 0 | 0 | - |
| Round 3 | 6 | 2 | RPE differentiation | 4 to 1 | 2020/02/05 15:02:00 | 2020/02/05 18:34:28 | 3:32:28 | 335 | 335 | 0 | 0 | - |

| | | | | | | | | | | | | |
|---|---|---|---|---|---|---|---|---|---|---|---|---|
| Round 3 | 7 | 1 | RPE differentiation | 1 to 4 | 2020/02/06 10:57:14 | 2020/02/06 14:34:23 | 3:37:09 | 335 | 335 | 0 | 0 | - |
| Round 3 | 7 | 2 | RPE differentiation | 5 to 8 | 2020/02/06 14:50:40 | 2020/02/06 18:19:29 | 3:28:49 | 335 | 335 | 0 | 0 | - |
| Round 3 | 8 | 1 | RPE differentiation | 8 to 5 | 2020/02/07 10:49:46 | 2020/02/07 14:21:17 | 3:31:31 | 335 | 335 | 0 | 0 | - |
| Round 3 | 8 | 2 | RPE differentiation | 4 to 1 | 2020/02/07 14:41:38 | 2020/02/07 18:19:31 | 3:37:53 | 335 | 335 | 0 | 0 | - |
| Round 3 | 9 | 1 | RPE differentiation | 1 to 4 | 2020/02/08 12:17:03 | 2020/02/08 15:47:23 | 3:30:20 | 335 | 335 | 0 | 0 | - |
| Round 3 | 9 | 2 | RPE differentiation | 5 to 8 | 2020/02/08 16:07:54 | 2020/02/08 19:39:30 | 3:31:36 | 335 | 335 | 0 | 0 | - |
| Round 3 | 10 | 1 | RPE differentiation | 8 to 5 | 2020/02/09 12:12:10 | 2020/02/09 15:42:21 | 3:30:11 | 335 | 335 | 0 | 0 | - |
| Round 3 | 10 | 2 | RPE differentiation | 4 to 1 | 2020/02/09 16:19:02 | 2020/02/09 19:44:19 | 3:25:17 | 335 | 335 | 0 | 0 | - |
| Round 3 | 11 | 1 | RPE differentiation | 1 to 4 | 2020/02/10 11:04:03 | 2020/02/10 14:20:52 | 3:16:49 | 335 | 335 | 0 | 0 | - |
| Round 3 | 11 | 2 | RPE differentiation | 5 to 8 | 2020/02/10 14:38:01 | 2020/02/10 17:59:34 | 3:21:33 | 335 | 335 | 0 | 0 | - |
| Round 3 | 12 | 1 | RPE differentiation | 8 to 5 | 2020/02/11 10:29:51 | 2020/02/11 13:49:54 | 3:20:03 | 335 | 335 | 0 | 0 | - |
| Round 3 | 12 | 2 | RPE differentiation | 4 to 1 | 2020/02/11 14:06:17 | 2020/02/11 17:26:52 | 3:20:35 | 335 | 335 | 0 | 0 | - |
| Round 3 | 13 | 1 | RPE differentiation | 1 to 4 | 2020/02/12 11:13:48 | 2020/02/12 14:30:45 | 3:16:57 | 335 | 335 | 0 | 0 | - |
| Round 3 | 13 | 2 | RPE differentiation | 5 to 8 | 2020/02/12 14:56:06 | 2020/02/12 18:14:33 | 3:18:27 | 335 | 335 | 0 | 0 | - |
| Round 3 | 14 | 1 | RPE differentiation | 8 to 5 | 2020/02/13 10:48:58 | 2020/02/13 14:01:34 | 3:12:36 | 335 | 335 | 0 | 0 | - |
| Round 3 | 14 | 2 | RPE differentiation | 4 to 1 | 2020/02/13 14:19:19 | 2020/02/13 17:31:45 | 3:12:26 | 335 | 335 | 0 | 0 | - |
| Round 3 | 15 | 1 | RPE differentiation | 1 to 4 | 2020/02/14 10:56:44 | 2020/02/14 14:10:38 | 3:13:54 | 335 | 335 | 0 | 0 | - |
| Round 3 | 15 | 2 | RPE differentiation | 5 to 8 | 2020/02/14 14:38:41 | 2020/02/14 17:50:15 | 3:11:34 | 335 | 335 | 0 | 0 | - |
| Round 3 | 16 | 1 | RPE differentiation | 8 to 5 | 2020/02/15 12:00:36 | 2020/02/15 15:06:18 | 3:05:42 | 335 | 335 | 0 | 0 | - |
| Round 3 | 16 | 2 | RPE differentiation | 4 to 1 | 2020/02/15 15:27:23 | 2020/02/15 18:31:39 | 3:04:16 | 335 | 335 | 0 | 0 | - |
| Round 3 | 17 | 1 | RPE differentiation | 1 to 4 | 2020/02/16 11:21:51 | 2020/02/16 14:20:40 | 2:58:49 | 335 | 335 | 0 | 0 | - |
| Round 3 | 17 | 2 | RPE differentiation | 5 to 8 | 2020/02/16 15:26:58 | 2020/02/16 18:21:43 | 2:54:45 | 335 | 335 | 0 | 0 | - |
| Round 3 | 18 | 1 | RPE differentiation | 8 to 5 | 2020/02/17 10:36:42 | 2020/02/17 13:09:04 | 2:32:22 | 335 | 335 | 0 | 0 | - |
| Round 3 | 18 | 2 | RPE differentiation | 4 to 1 | 2020/02/17 13:22:34 | 2020/02/17 15:57:49 | 2:35:15 | 335 | 335 | 0 | 0 | - |
| Round 3 | 19 | 1 | RPE differentiation | 1 to 4 | 2020/02/18 11:04:49 | 2020/02/18 13:36:47 | 2:31:58 | 335 | 335 | 0 | 0 | - |
| Round 3 | 19 | 2 | RPE differentiation | 5 to 8 | 2020/02/18 14:01:25 | 2020/02/18 16:31:50 | 2:30:25 | 335 | 335 | 0 | 0 | - |
| Round 3 | 20 | 1 | RPE differentiation | 8 to 5 | 2020/02/19 10:48:18 | 2020/02/19 12:25:46 | 1:37:28 | 77 | 77 | 0 | 0 | - |
| Round 3 | 20 | 2 | RPE differentiation | 4 to 1 | 2020/02/19 12:32:42 | 2020/02/19 14:12:55 | 1:40:13 | 77 | 77 | 0 | 0 | - |
| Round 3 | 21 | 1 | RPE differentiation | 1 to 4 | 2020/02/20 11:00:04 | 2020/02/20 12:39:46 | 1:39:42 | 77 | 77 | 0 | 0 | - |
| Round 3 | 21 | 2 | RPE differentiation | 5 to 8 | 2020/02/20 12:57:08 | 2020/02/20 14:35:49 | 1:38:41 | 77 | 77 | 0 | 0 | - |
| Round 3 | 22 | 1 | RPE differentiation | 8 to 5 | 2020/02/21 11:15:01 | 2020/02/21 12:53:42 | 1:38:41 | 77 | 77 | 0 | 0 | - |
| Round 3 | 22 | 2 | RPE differentiation | 4 to 1 | 2020/02/21 13:19:54 | 2020/02/21 14:58:56 | 1:39:02 | 77 | 77 | 0 | 0 | - |
| Round 3 | 23 | 1 | RPE differentiation | 1 to 4 | 2020/02/22 11:14:10 | 2020/02/22 12:53:50 | 1:39:40 | 77 | 77 | 0 | 0 | - |
| Round 3 | 23 | 2 | RPE differentiation | 5 to 8 | 2020/02/22 13:02:20 | 2020/02/22 14:39:52 | 1:37:32 | 77 | 77 | 0 | 0 | - |
| Round 3 | 24 | 1 | RPE differentiation | 8 to 5 | 2020/02/23 11:54:56 | 2020/02/23 13:32:46 | 1:37:50 | 77 | 77 | 0 | 0 | - |
| Round 3 | 24 | 2 | RPE differentiation | 4 to 1 | 2020/02/23 13:56:51 | 2020/02/23 15:36:55 | 1:40:04 | 77 | 77 | 0 | 0 | - |
| Round 3 | 25 | 1 | RPE differentiation | 1 to 4 | 2020/02/24 11:47:49 | 2020/02/24 13:28:01 | 1:40:12 | 77 | 77 | 0 | 0 | - |
| Round 3 | 25 | 2 | RPE differentiation | 5 to 8 | 2020/02/24 13:48:53 | 2020/02/24 15:22:06 | 1:33:13 | 77 | 77 | 0 | 0 | - |
| Round 3 | 26 | 1 | RPE maintenance | 8 to 5 | 2020/02/25 10:51:04 | 2020/02/25 12:28:52 | 1:37:48 | 77 | 77 | 0 | 0 | - |
| Round 3 | 26 | 2 | RPE maintenance | 4 to 1 | 2020/02/25 12:50:19 | 2020/02/25 14:29:00 | 1:38:41 | 77 | 77 | 0 | 0 | - |
| Round 3 | 28 | 1 | RPE maintenance | 1 to 4 | 2020/02/27 10:47:29 | 2020/02/27 12:27:10 | 1:39:41 | 77 | 77 | 0 | 0 | - |
| Round 3 | 28 | 2 | RPE maintenance | 5 to 8 | 2020/02/27 13:20:27 | 2020/02/27 14:58:11 | 1:37:44 | 77 | 77 | 0 | 0 | - |
| Round 3 | 30 | 1 | RPE maintenance | 8 to 5 | 2020/02/29 11:09:03 | 2020/02/29 12:47:54 | 1:38:51 | 77 | 77 | 0 | 0 | - |
| Round 3 | 30 | 2 | RPE maintenance | 4 to 1 | 2020/02/29 12:52:16 | 2020/02/29 14:31:08 | 1:38:52 | 77 | 77 | 0 | 0 | - |
| Round 3 | 32 | 1 | RPE maintenance | 1 to 4 | 2020/03/02 10:55:35 | 2020/03/02 12:35:13 | 1:39:38 | 77 | 77 | 0 | 0 | - |
| Round 3 | 32 | 2 | RPE maintenance | 5 to 8 | 2020/03/02 12:40:00 | 2020/03/02 14:18:08 | 1:38:08 | 77 | 77 | 0 | 0 | - |
| Validation | -7 | 1 | Seeding | 4 to 1, 8 to 5 | 2020/03/19 12:08:21 | 2020/03/19 13:56:49 | 1:48:28 | 89 | 89 | 0 | 0 | - |
| Validation | -6 | 1 | Preconditioning | 1 to 4 | 2020/03/20 12:21:02 | 2020/03/20 14:40:31 | 2:19:29 | 193 | 193 | 0 | 0 | - |
| Validation | -6 | 2 | Preconditioning | 5 to 8 | 2020/03/20 14:52:24 | 2020/03/20 17:11:03 | 2:18:39 | 193 | 193 | 0 | 0 | - |
| Validation | -5 | 1 | Preconditioning | 8 to 5 | 2020/03/21 11:21:43 | 2020/03/21 13:49:58 | 2:28:15 | 193 | 193 | 0 | 0 | - |
| Validation | -5 | 2 | Preconditioning | 4 to 1 | 2020/03/21 14:39:40 | 2020/03/21 17:08:07 | 2:28:27 | 193 | 193 | 0 | 0 | - |
| Validation | -4 | 1 | Preconditioning | 1 to 4 | 2020/03/22 11:41:00 | 2020/03/22 14:09:42 | 2:28:42 | 193 | 193 | 0 | 0 | - |
| Validation | -4 | 2 | Preconditioning | 5 to 8 | 2020/03/22 14:24:09 | 2020/03/22 16:52:43 | 2:28:34 | 193 | 193 | 0 | 0 | - |
| Validation | -3 | 1 | Preconditioning | 8 to 5 | 2020/03/23 11:02:28 | 2020/03/23 13:30:41 | 2:28:13 | 193 | 193 | 0 | 0 | - |
| Validation | -3 | 2 | Preconditioning | 4 to 1 | 2020/03/23 13:45:00 | 2020/03/23 16:13:54 | 2:28:54 | 193 | 193 | 0 | 0 | - |
| Validation | -2 | 1 | Preconditioning | 1 to 4 | 2020/03/24 10:54:41 | 2020/03/24 13:22:54 | 2:28:13 | 193 | 193 | 0 | 0 | - |
| Validation | -2 | 2 | Preconditioning | 5 to 8 | 2020/03/24 13:35:32 | 2020/03/24 16:03:37 | 2:28:05 | 193 | 193 | 0 | 0 | - |
| Validation | -1 | 1 | Preconditioning | 8 to 5 | 2020/03/25 10:51:01 | 2020/03/25 13:19:33 | 2:28:32 | 193 | 193 | 0 | 0 | - |
| Validation | -1 | 2 | Preconditioning | 4 to 1 | 2020/03/25 13:32:00 | 2020/03/25 16:01:31 | 2:29:31 | 193 | 193 | 0 | 0 | - |
| Validation | 0 | 1 | Passage | 1 to 4 | 2020/03/26 09:50:33 | 2020/03/26 14:54:29 | 5:03:56 | 151 | 151 | 0 | 0 | - |
| Validation | 0 | 2 | Passage | 5 to 8 | 2020/03/26 15:58:43 | 2020/03/26 21:02:35 | 5:03:52 | 151 | 151 | 0 | 0 | - |
| Validation | 1 | 1 | RPE differentiation | 1 to 4 | 2020/03/27 11:03:34 | 2020/03/27 14:32:45 | 3:29:11 | 335 | 335 | 0 | 0 | - |
| Validation | 1 | 2 | RPE differentiation | 5 to 8 | 2020/03/27 14:45:31 | 2020/03/27 18:13:14 | 3:27:43 | 335 | 335 | 0 | 0 | - |
| Validation | 2 | 1 | RPE differentiation | 8 to 5 | 2020/03/28 11:09:54 | 2020/03/28 14:36:49 | 3:26:55 | 335 | 335 | 0 | 0 | - |
| Validation | 2 | 2 | RPE differentiation | 4 to 1 | 2020/03/28 15:09:07 | 2020/03/28 18:36:40 | 3:27:33 | 335 | 335 | 0 | 0 | - |
| Validation | 3 | 1 | RPE differentiation | 1 to 4 | 2020/03/29 11:18:54 | 2020/03/29 14:44:47 | 3:25:53 | 335 | 335 | 0 | 0 | - |
| Validation | 3 | 2 | RPE differentiation | 5 to 8 | 2020/03/29 15:10:41 | 2020/03/29 18:36:30 | 3:25:49 | 335 | 335 | 0 | 0 | - |
| Validation | 4 | 1 | RPE differentiation | 8 to 5 | 2020/03/30 10:58:35 | 2020/03/30 14:16:32 | 3:17:57 | 335 | 335 | 0 | 0 | - |
| Validation | 4 | 2 | RPE differentiation | 4 to 1 | 2020/03/30 14:30:35 | 2020/03/30 17:49:07 | 3:18:32 | 335 | 335 | 0 | 0 | - |
| Validation | 5 | 1 | RPE differentiation | 1 to 4 | 2020/03/31 10:42:54 | 2020/03/31 13:54:05 | 3:11:11 | 335 | 335 | 0 | 0 | - |
| Validation | 5 | 2 | RPE differentiation | 5 to 8 | 2020/03/31 14:09:58 | 2020/03/31 17:20:02 | 3:10:04 | 335 | 335 | 0 | 0 | - |
| Validation | 6 | 1 | RPE differentiation | 8 to 5 | 2020/04/01 10:42:30 | 2020/04/01 13:51:42 | 3:09:12 | 335 | 335 | 0 | 0 | - |
| Validation | 6 | 2 | RPE differentiation | 4 to 1 | 2020/04/01 14:03:21 | 2020/04/01 17:13:11 | 3:09:50 | 335 | 335 | 0 | 0 | - |
| Validation | 7 | 1 | RPE differentiation | 1 to 4 | 2020/04/02 10:45:16 | 2020/04/02 13:58:42 | 3:13:26 | 335 | 335 | 0 | 0 | - |
| Validation | 7 | 2 | RPE differentiation | 5 to 8 | 2020/04/02 14:13:40 | 2020/04/02 17:26:39 | 3:12:59 | 335 | 335 | 0 | 0 | - |
| Validation | 8 | 1 | RPE differentiation | 8 to 5 | 2020/04/03 10:57:42 | 2020/04/03 14:07:54 | 3:10:12 | 335 | 335 | 0 | 0 | - |
| Validation | 8 | 2 | RPE differentiation | 4 to 1 | 2020/04/03 14:22:42 | 2020/04/03 17:32:11 | 3:09:29 | 335 | 335 | 0 | 0 | - |
| Validation | 9 | 1 | RPE differentiation | 1 to 4 | 2020/04/04 11:39:21 | 2020/04/04 14:50:54 | 3:11:33 | 335 | 335 | 0 | 0 | - |
| Validation | 9 | 2 | RPE differentiation | 5 to 8 | 2020/04/04 15:11:22 | 2020/04/04 18:22:00 | 3:10:38 | 335 | 335 | 0 | 0 | - |
| Validation | 10 | 1 | RPE differentiation | 8 to 5 | 2020/04/05 10:53:44 | 2020/04/05 14:01:48 | 3:08:04 | 335 | 335 | 0 | 0 | - |
| Validation | 10 | 2 | RPE differentiation | 4 to 1 | 2020/04/05 14:19:51 | 2020/04/05 17:27:56 | 3:08:05 | 335 | 335 | 0 | 0 | - |
| Validation | 11 | 1 | RPE differentiation | 1 to 4 | 2020/04/06 10:44:01 | 2020/04/06 13:50:03 | 3:06:02 | 335 | 335 | 0 | 0 | - |
| Validation | 11 | 2 | RPE differentiation | 5 to 8 | 2020/04/06 14:03:00 | 2020/04/06 17:08:10 | 3:05:10 | 335 | 335 | 0 | 0 | - |
| Validation | 12 | 1 | RPE differentiation | 8 to 5 | 2020/04/07 10:35:01 | 2020/04/07 13:32:56 | 2:57:55 | 335 | 335 | 0 | 0 | - |
| Validation | 12 | 2 | RPE differentiation | 4 to 1 | 2020/04/07 14:04:51 | 2020/04/07 17:01:06 | 2:56:15 | 335 | 335 | 0 | 0 | - |
| Validation | 13 | 1 | RPE differentiation | 1 to 4 | 2020/04/08 11:27:10 | 2020/04/08 14:21:01 | 2:53:51 | 335 | 335 | 0 | 0 | - |
| Validation | 13 | 2 | RPE differentiation | 5 to 8 | 2020/04/08 15:04:27 | 2020/04/08 17:58:07 | 2:53:40 | 335 | 335 | 0 | 0 | - |
| Validation | 14 | 1 | RPE differentiation | 8 to 5 | 2020/04/09 10:48:02 | 2020/04/09 13:37:57 | 2:49:55 | 335 | 335 | 0 | 0 | - |
| Validation | 14 | 2 | RPE differentiation | 4 to 1 | 2020/04/09 13:49:26 | 2020/04/09 16:40:07 | 2:50:41 | 335 | 335 | 0 | 0 | - |
| Validation | 15 | 1 | RPE differentiation | 1 to 4 | 2020/04/10 10:39:41 | 2020/04/10 13:35:35 | 2:55:54 | 335 | 334 | 1 | 0 | Defective labware error |
| Validation | 15 | 2 | RPE differentiation | 5 to 8 | 2020/04/10 13:47:27 | 2020/04/10 16:37:57 | 2:50:30 | 335 | 335 | 0 | 0 | - |
| Validation | 16 | 1 | RPE differentiation | 8 to 5 | 2020/04/11 11:07:36 | 2020/04/11 13:55:56 | 2:48:20 | 335 | 335 | 0 | 0 | - |
| Validation | 16 | 2 | RPE differentiation | 4 to 1 | 2020/04/11 14:24:46 | 2020/04/11 17:13:04 | 2:48:18 | 335 | 335 | 0 | 0 | - |
| Validation | 17 | 1 | RPE differentiation | 1 to 4 | 2020/04/12 10:59:29 | 2020/04/12 13:44:04 | 2:44:35 | 335 | 335 | 0 | 0 | - |
| Validation | 17 | 2 | RPE differentiation | 5 to 8 | 2020/04/12 14:00:25 | 2020/04/12 16:45:06 | 2:44:41 | 335 | 335 | 0 | 0 | - |
| Validation | 18 | 1 | RPE differentiation | 8 to 5 | 2020/04/13 11:17:36 | 2020/04/13 13:55:07 | 2:37:31 | 335 | 335 | 0 | 0 | - |
| Validation | 18 | 2 | RPE differentiation | 4 to 1 | 2020/04/13 14:17:37 | 2020/04/13 16:54:16 | 2:36:39 | 335 | 335 | 0 | 0 | - |
| Validation | 19 | 1 | RPE differentiation | 1 to 4 | 2020/04/14 09:59:29 | 2020/04/14 12:27:42 | 2:28:13 | 335 | 335 | 0 | 0 | - |
| Validation | 19 | 2 | RPE differentiation | 5 to 8 | 2020/04/14 12:40:42 | 2020/04/14 15:09:17 | 2:28:35 | 335 | 335 | 0 | 0 | - |
| Validation | 20 | 1 | RPE differentiation | 8 to 5 | 2020/04/15 12:34:37 | 2020/04/15 14:13:10 | 1:38:33 | 77 | 77 | 0 | 0 | - |
| Validation | 20 | 2 | RPE differentiation | 4 to 1 | 2020/04/15 14:48:03 | 2020/04/15 16:24:19 | 1:36:16 | 77 | 77 | 0 | 0 | - |
| Validation | 21 | 1 | RPE differentiation | 1 to 4 | 2020/04/16 11:04:38 | 2020/04/16 12:41:13 | 1:36:35 | 77 | 77 | 0 | 0 | - |
| Validation | 21 | 2 | RPE differentiation | 5 to 8 | 2020/04/16 13:16:09 | 2020/04/16 14:52:17 | 1:36:08 | 77 | 77 | 0 | 0 | - |
| Validation | 22 | 1 | RPE differentiation | 8 to 5 | 2020/04/17 10:50:08 | 2020/04/17 12:27:07 | 1:36:59 | 77 | 77 | 0 | 0 | - |
| Validation | 22 | 2 | RPE differentiation | 4 to 1 | 2020/04/17 12:45:59 | 2020/04/17 14:22:17 | 1:36:18 | 77 | 77 | 0 | 0 | - |
| Validation | 23 | 1 | RPE differentiation | 1 to 4 | 2020/04/18 11:04:36 | 2020/04/18 12:41:11 | 1:36:35 | 77 | 77 | 0 | 0 | - |
| Validation | 23 | 2 | RPE differentiation | 5 to 8 | 2020/04/18 13:16:05 | 2020/04/18 14:52:21 | 1:36:16 | 77 | 77 | 0 | 0 | - |
| Validation | 24 | 1 | RPE differentiation | 8 to 5 | 2020/04/19 10:58:06 | 2020/04/19 12:36:06 | 1:38:00 | 77 | 77 | 0 | 0 | - |
| Validation | 24 | 2 | RPE differentiation | 4 to 1 | 2020/04/19 12:41:16 | 2020/04/19 14:19:30 | 1:38:14 | 77 | 77 | 0 | 0 | - |
| Validation | 25 | 1 | RPE differentiation | 1 to 4 | 2020/04/20 11:04:36 | 2020/04/20 14:01:25 | 2:56:49 | 77 | 76 | 1 | 0 | Microscope error |
| Validation | 25 | 2 | RPE differentiation | 5 to 8 | 2020/04/20 14:36:20 | 2020/04/20 16:13:27 | 1:37:07 | 77 | 77 | 0 | 0 | - |
| Validation | 26 | 1 | RPE maintenance | 8 to 5 | 2020/04/21 11:06:08 | 2020/04/21 12:39:17 | 1:33:09 | 77 | 77 | 0 | 0 | - |
| Validation | 26 | 2 | RPE maintenance | 4 to 1 | 2020/04/21 12:42:56 | 2020/04/21 14:19:57 | 1:37:01 | 77 | 77 | 0 | 0 | - |
| Validation | 28 | 1 | RPE maintenance | 1 to 4 | 2020/04/23 11:12:10 | 2020/04/23 12:51:25 | 1:39:15 | 77 | 77 | 0 | 0 | - |
| Validation | 28 | 2 | RPE maintenance | 5 to 8 | 2020/04/23 12:54:42 | 2020/04/23 14:31:29 | 1:36:47 | 77 | 77 | 0 | 0 | - |
| Validation | 30 | 1 | RPE maintenance | 8 to 5 | 2020/04/25 11:45:49 | 2020/04/25 13:23:25 | 1:37:36 | 77 | 77 | 0 | 0 | - |
| Validation | 30 | 2 | RPE maintenance | 4 to 1 | 2020/04/25 13:28:38 | 2020/04/25 15:06:25 | 1:37:47 | 77 | 77 | 0 | 0 | - |
| Validation | 32 | 1 | RPE maintenance | 1 to 4 | 2020/04/27 10:43:29 | 2020/04/27 12:22:34 | 1:39:05 | 77 | 77 | 0 | 0 | - |
| Validation | 32 | 2 | RPE maintenance | 5 to 8 | 2020/04/27 12:25:00 | 2020/04/27 14:01:32 | 1:36:32 | 77 | 77 | 0 | 0 | - |

**Table S6**

| Protein | Sample name | Protein expression [ng/mL] | | | | |
|---|---|---|---|---|---|---|
| | | Replicate 1 | Replicate 2 | Replicate 3 | Mean | SEM |
| VEGF | hiPSC | N.D. | N.D. | N.D. | N.D. | N.D. |
| | H-RPE | 1.43 | 1.38 | 1.47 | 1.43 | 0.03 |
| | Pre-optimized | 6.53 | 7.36 | 8.33 | 7.41 | 0.52 |
| | Optimized-1 | 4.76 | 5.52 | 4.78 | 5.02 | 0.25 |
| | Optimized-2 | 7.75 | 9.17 | 8.63 | 8.52 | 0.41 |
| | Optimized-3 | 8.56 | 7.61 | 5.72 | 7.30 | 0.83 |
| | Optimized-4 | 6.45 | 6.63 | 8.71 | 7.26 | 0.72 |
| | Optimized-5 | 6.92 | 8.33 | 7.28 | 7.51 | 0.42 |
| | hiPSC medium | N.D. | | | | |
| | RPE medium | N.D. | | | | |
| PEDF | hiPSC | N.D. | N.D. | N.D. | N.D. | N.D. |
| | H-RPE | 0.36 | 0.43 | 0.43 | 0.41 | 0.02 |
| | Pre-optimized | 1.50 | 1.41 | 1.41 | 1.44 | 0.03 |
| | Optimized-1 | 1.31 | 1.21 | 1.32 | 1.28 | 0.04 |
| | Optimized-2 | 1.74 | 1.70 | 1.60 | 1.68 | 0.04 |
| | Optimized-3 | 1.50 | 1.33 | 1.34 | 1.39 | 0.06 |
| | Optimized-4 | 1.14 | 1.30 | 1.33 | 1.25 | 0.06 |
| | Optimized-5 | 1.26 | 1.20 | 1.43 | 1.29 | 0.07 |
| | hiPSC medium | N.D. | | | | |
| | RPE medium | N.D. | | | | |
